# Navigating directed evolution efficiently: optimizing selection conditions and selection output analysis

**DOI:** 10.1101/2024.05.23.595501

**Authors:** Paola Handal-Marquez, Hoai Nguyen, Vitor B. Pinheiro

## Abstract

Directed evolution is a powerful tool that can bypass gaps in our understanding of the sequence-function relationship of proteins and still isolate variants with desired activities, properties, and substrate specificities. The rise of directed evolution platforms for polymerase engineering has accelerated the isolation of xenobiotic nucleic acids (XNAs) synthetases and reverse transcriptases (RTs) capable of processing a wide array of unnatural XNAs which have numerous therapeutic and biotechnological applications. Still, the current generation of XNA polymerases functions with significantly lower efficiency than the natural counterparts and retains a significant level of DNA polymerase activity which limits their *in vivo* applications. Although directed evolution approaches are continuously being developed and implemented to improve XNA polymerase engineering, the field lacks an in-depth analysis of the effect of selection parameters, library construction biases and sampling biases. Focusing on the directed evolution pipeline for DNA and XNA polymerase engineering, this work sets out a method for understanding the impact of selection conditions on selection success and efficiency. We also explore the influence of selection conditions on fidelity at the population and individual mutant level. Additionally, we explore the sequencing coverage requirements in directed evolution experiments, which differ from genome assembly and other *-omics* approaches. This analysis allowed us to identify the sequencing coverage threshold for the accurate and precise identification of significantly enriched mutants. Overall, this study introduces a robust methodology for optimizing selection protocols, which effectively streamlines selection processes by employing small libraries and cost-effective NGS sequencing. It provides valuable insights into critical considerations, thereby enhancing the overall effectiveness and efficiency of directed evolution strategies applicable to enzymes other than the ones considered here.

## 1 Introduction

Protein evolution can be represented as an adaptive walk on a sequence-functional landscape, a concept pioneered by Sewall Wright in 1932 and commonly referred to as the “fitness landscape.” Here, sequences (genotypes) are mapped to a quantitative measure of fitness such as enzymatic activity, thermostability or other physicochemical properties (Eigen, 1985; Kauffman and Levin, 1987; Kauffman, 1992) (phenotypes). In this framework, closely related sequences are proximal in the fitness map and sequences can occupy peaks (high fitness) or valleys (low fitness) on the landscape (Kauffman and Levin, 1987; Kondrashov and Kondrashov, 2015; Macken and Perelson, 1989; Wittmund et al., 2022). In natural evolution, the term “adaptive walk” describes the evolutionary changes in the sequences on the fitness landscape taken to reach the global (maximum) fitness peak within a particular environment.

This natural process can be mimicked *in vitro* through directed evolution, which encompasses two complementary approaches. On one hand, a stepwise process of mutation-screening-learning that reaches functional maximum through the sequential accumulation of beneficial mutations. On the other hand, an approach where strong genotype-phenotype linkages allow ultra-high-throughput strategies that sample the genotype space more widely in searching for the functional maximum (Tizei et al., 2016). The latter approach proves especially valuable in situations where understanding of the structure-sequence-function relationships of the target protein is limited, yet it still facilitates the isolation of variants with desired activities, properties, and substrate specificities.

Despite partial understanding of the factors governing polymerase specificity and selectivity, the rise of directed evolution *in vitro* selection platforms has enabled the engineering of variants with novel and improved properties. Emulsion-based selection platforms have successfully isolated variants with improved thermostability (Ghadessy et al., 2001; Povilaitis et al., 2016), variants capable of incorporating nucleotide analogues (Larsen et al., 2016; Loakes et al., 2009; Pinheiro et al., 2012; Ramsay et al., 2010), DNA polymerase variants capable of reverse transcription (RTs) (Ellefson et al., 2016; Houlihan et al., 2020; Pinheiro et al., 2012) and RNA polymerase variants with altered promoter recognition (Abil et al., 2017). Selection platforms based on phage display have also been implemented to expand the substrate spectrum of thermophilic DNA polymerases (Chen et al., 2016).

These selection platforms establish a strong phenotype-genotype link through the emulsification of individual cells expressing a unique variant, substrates, and products. This minimizes cross-reactivity and cross-catalysis allowing the partitioning of libraries based on enzyme function (substrate recognition, product formation, and/or synthesis rate) of individual variants. Recovered selection outputs can be subcloned for subsequent rounds of selection, screened, or deep sequenced to recover the genotype of enriched variants for further characterization.

As the engineering challenge increases towards more distant substrates, the likelihood of failure increases but nothing can be learned from those - it is not clear whether this is a limitation of the methodology, selection conditions, library, or the target substrate. Therefore, tools that can give us better insight into the selection process itself are essential to better address selection challenges, such as the recovery of false positives. False positives are variants that are recovered due to random and non-specific processes (background) or due to viable alternative but non-desired phenotypes (parasites) (Tizei et al., 2016). For instance, a selection parasite in CSR would be a variant able to use the low cellular concentrations of dNTPs present in the emulsion instead of the analogues provided.

Selection parameters such as cofactor concentration, play a crucial role in shaping the activity of polymerases in directed evolution, potentially influencing the cooperative interplay between polymerase and exonuclease activities, or leading to increased parasite recovery. However, determining the optimal selection parameters for biasing evolution towards the desired variants on a polymerase library of unknown function, as often the case when engineering new-to-nature substrate specificities, is a non-trivial task.

Here, we propose a pipeline that incorporates Design of Experiments (DoE) for screening and benchmarking selection parameters using a small protein library. This approach enables the parameters and concentration ranges of selection criteria to be optimized, thereby enhancing the efficacy of the selection process and achieving optimal results with larger and more complex libraries.

To validate this strategy, we used a small, focused library targeting a B-family polymerase metal-coordinating residue (D404) (Kropp et al., 2017) and neighboring residues in *Thermococcus kodakarensis* DNA polymerase (KOD DNAP), to explore the influence of CSR selection parameters on the outputs of selection. Selection parameters (factors) investigated included nucleotide concentration, nucleotide chemistry, selection time, Mg^2+^ and/or Mn^2+^ concentration, as well as other commonly used PCR additives. Selection outputs (responses) analyzed included recovery yield, variant enrichment, and variant fidelity.

The result was the rapid optimization of selection parameters, maximizing the efficiency of selection. In addition, further analysis on the balance of synthesis efficiency and fidelity (a window into the polymerase/exonuclease equilibrium) can be used to gain biological insight into polymerase mechanisms. Our data also confirmed that cost-effective, precise, and accurate identification of active variants is possible even at low coverages.

Together, the method described here establishes a more systematic pipeline for the engineering of XNA polymerases and that can also be applied for other enzymes. It shows how much selection impacts the library in a single round of selection and how robust CSR (and presumably other emulsion-based strategies) is as a directed evolution platform.

## 2 Materials and methods

### 1.1 Library design and construction

The 2-point saturation mutagenesis library targeting the catalytic D404 and its vicinal L403 residue was generated using mutagenic primers KOD_Sat_403-404_F1 and KOD_Sat_403-404_R1 (Table S1). The 5-point saturation mutagenesis library targeting L403, D404, F405, L408 and Y409 was generated using primers KOD_5ptSat_408-409_F1 and KOD_5ptSat_403-405_R1. Both libraries were assembled through inverse PCR (iPCR) on the pET23-KOD-Exo-(Supplementary Information S1 Table S2) plasmid in Q5 High-Fidelity DNA Polymerase (NEB) reactions of 28 cycles following the manufacturer’s standard protocol. Amplified products were digested with DpnI (NEB) and purified with the Monarch® PCR & DNA Cleanup Kit (NEB). An aliquot (1 µg) of each library was blunt-end ligated in 100 µl reactions containing 1x T4 DNA ligase buffer, T4 DNA ligase (4 units/µl, final concentration) and T4 Polynucleotide Kinase (0.1 units/µl, final concentration) overnight at room temperature. Ligations were purified through phenol:chloroform:isoamyl alcohol extraction and ethanol precipitation. 1 µg of each purified ligated library was transformed in 10-beta competent *E. coli* cells (NEB). The cells were freshly prepared to maximise transformation efficiency. Briefly, 50 mL of fresh LB media per library was inoculated with an overnight culture of 10-beta cells and washed 3 times with 30 mL of 1mM HEPES buffer ((4-(2-hydroxyethyl)-1-piperazineethanesulfonic acid) pH 7.0, Sigma-Aldrich) once it reached OD_600_ of 0.4. Cells were pelleted and resuspended in 400 µl of HEPES buffer, mixed with the ligation mixtures and transferred to chilled 2 mm electroporation cuvettes (BioRad). Cells were transformed using a Gene Pulser II electoporator (BioRad) at 12.5 kV/cm, 200 Ω and 25 µF. The cells were resuspended in 5 mL of fresh LB media, incubated at 37°C for 1 h, pelleted through centrifugation at 4000 rpm (3250 *g*), resuspended in 1 mL of LB and finally plated in 24.5 cm x 24.5 cm LB plates with 100 µg/µl of Ampicillin. After overnight incubation at 37°C, libraries were scraped and resuspended in 5 mL of LB supplemented with 50 µg/mL ampicillin and 25% filter-sterilized glycerol. Resuspended cells were then split into 5 cryovial tubes and stored at -80°C. Libraries were aliquoted in 10 mL (OD600 ∼ 0.2) and re-grown for 2-3 h to extract plasmids using the GeneJET Plasmid Miniprep Kit (Thermo Fisher Scientific) for sequencing and for subcloning into the expression strain, BL21(DE3) (NEB) following the same transformation procedure described. The KODΔ mutant was generated using primers PH_Delta_KOD_F1/R1 (Table S1) through iPCR and cloned into the final BL21(DE3) (NEB) strain following the same procedure as the library construction.

### 1.2 Design of Experiments (DoE) implementation

An initial design (D1) was performed to test the main effects of several parameters including nucleotide, magnesium, manganese, BSA, betaine, PEG1000 and formamide concentrations as well as extension time and nucleotide type on selection efficiency and mutant behavior. A summary of the selection parameters be found in **Table 1**. A linear model with D optimality was used to design 12 unique selection parameter combinations for testing using the skpr package (version 1.7.1) in RStudio (version 4.3.1). All designs were run in blocks of 12 randomized experiments. Design 2 (D2) explored only nucleotide type, manganese, betaine and formamide concentrations. A central composite design in JMP Pro 16 with 36 unique selection parameter combinations that tested main effects and 2-way interactions was generated. We additionally included 12 negative control selections with no nucleotides and random concentrations of the other factors (scattered across the experimental design blocks), which were used for noise correction during quantification. Since all 12 negative controls from D2 showed negligible signal, in subsequent designs, we included only one negative control per block with no nucleotides nor other additives. Design 3 (D3) and Design 4 (D4) explored nucleotide, magnesium, manganese, betaine and PEG1000 concentrations. A custom design with D optimality in JMP Pro 16 with 24 unique selection parameter combinations that explored main effects, 2-factor interactions, and quadratic effects was generated. Detailed experimental conditions from each design can be found in **Supplementary Information S3.**

### 1.3 Compartmentalized self-replication (CSR)

Libraries or KODΔ were aliquoted from glycerol stocks in 5 mL of LB supplemented with 100 µg/mL ampicillin (OD600 ∼ 0.2), grown to OD_600_ 0.7 and induced with 1 mM IPTG for 4 h at 30°C. A total of 2x10^8^ cells were isolated and pelleted through centrifugation at 4000 rpm (3250 *g*) for 5 minutes. For design 1 (D1), selections were spiked with 10% KODΔ, resulting in a mixture were 90% of the total cell density corresponded to the 2-point library and the remaining 10% to KODΔ. D2, D3 and D4, were not spiked with KODΔ to facilitate sample handling and downstream quantification of the reactions, but negative controls without nucleotides were included. Pellets were resuspended in selection mixtures containing the proposed concentrations of each component by the models as well as 1x Thermopol buffer, 0.005 mg/ml RNAse A and 1 µM of each selection primer (CSR_Sel_SHORT_F3/R3, Supplementary Information S1 Table S1). Following recommendations for emulsion composition previously published (Pinheiro et al., 2014), the resuspended cells were aliquoted in a 2 mL round bottomed microcentrifuge tube containing a 5-mm steel bead and were then overlaid with 500 µl of an oil mix composed of 73% TEGOSOFT DEC (Evonik), 20% mineral oil (Sigma-Aldrich) and 7% ABIL WE09 (Evonik). The microcentrifuge tubes were transferred to a TissueLyser II (Qiagen) for emulsification at 15 Hz for 10 s followed by 17 Hz for 7 s. Emulsions were transferred to PCR strips and into a thermocycler with the following parameters: 5 min at 94°C followed by 20 cycles of 1 min at 94°C, 1 min at 61°C and 0 – 80 s at 68°C. Emulsions were collected in 2 mL tubes and 100 µL of 1X TBS buffer (10 mM Tris•Cl pH 7.4, 20 mM NaCl, 0.02% (v/v) Tween20) and 1 mL of water saturated 1-Hexanol were added. Emulsions were vortexed and centrifuged at 13,000 *g* for 10 min. The oil-hexanol phase was discarded and 700 µL of water saturated 1-hexanol was added, samples vortexed and centrifuged. The bottom aqueous phase was then recovered and purified through phenol:chloroform:isoamyl alcohol extraction and ethanol precipitation. Pellets were resuspended in 16 µL of elution buffer and mixed with 1x Cutsmart buffer, 1 µl DpnI, 1 µl ExoI (NEB) and incubated for 1 h at 37°C followed by heat inactivation of 20 min at 80°C. Selections were then purified using the Monarch® PCR & DNA Cleanup Kit (NEB), eluting in 10 µl of the Monarch® DNA Elution Buffer. Selection generally yields products below the limit of detection with common DNA quantification methods; thus, the products undergo a recovery PCR using primers that hybridize to overhangs introduced by selection primers, ensuring that only selection products are amplified (Figure 1B). Taq DNAP is typically used for recovery because of its ability to amplify even single-copy templates. Recovery PCR products can then be further amplified in innest PCR reactions to introduce overhangs for cloning or to prepare amplicons for NGS using ultra high fidelity DNAPs such as KOD Xtreme (EMD Millipore). Recovery PCR reactions were carried out with 1 µl of the selection product in Taq DNA polymerase (NEB) reactions with recovery primers (CSR_Rec_SHORT_F1/R1, Supplementary Information S1 Table S1) and 20 – 28 cycles. 1 µl of ExoI and 1 µl rSAP (NEB) were added to the recovery products and incubated for 15 min at 37°C followed by heat inactivation of 20 min at 80°C. Recovery products were then purified with the Monarch® PCR & DNA Cleanup Kit (NEB) and eluted in 10 µl of the Monarch® DNA Elution Buffer. In-nest PCR reactions of 28 cycles were then carried out using either the spCSR_Innest_MUT_F3/R3 primers (Supplementary Information S1 Table S1) primers or the NGS amplicon generation primers (Supplementary Information S1 Table S1) directly, and 2 µl of the purified recovery products in 50 µl KOD XTREME (Sigma-Aldrich) following the manufacturer’s recommended protocol. Reactions carried out with the spCSR_Innest_MUT primers were subcloned into the pET23-KOD-Exo-(Supplementary Information S1 Table S2) through Type IIS cloning and transformed into electrocompetent *E.coli* cells as previously described. Libraries were grown in 10 mL LB for 2-3 h and plasmids extracted using the GeneJET Plasmid Miniprep Kit (Thermo Fisher Scientific). From each library, 10 ng were then amplified using the NGS amplicon generation primers (Supplementary Information S1 Table S1) as follows: D1 2-pt library pre-selection was amplified with primers WP2_D1_Seq_F1_R0 and WP2_D1_Seq_R1; D1 2-pt selections 2, 5, 7, 8 and 11 were amplified with forward primers WP2_D1_Seq_F1_lib2, WP2_D1_Seq_F1_lib5, WP2_D1_Seq_F1_lib7, WP2_D1_Seq_F1_lib8, and WP2_D1_Seq_F1_lib11 respectively together with reverse primer WP2_D1_Seq_R1; D4 5-pt library pre-selection as well as selections 4, 8 and 20, were amplified with primers WP2_D4_Seq_F1 and WP2_D4_Seq_R1. NGS amplicon reactions were carried out in 50 µl KOD XTREME (Sigma-Aldrich) following the manufacturer’s recommended protocol. All reactions were then digested with 1 µl ExoI and purified using the Monarch DNA Gel Extraction Kit (NEB), eluting in 10 µl of elution buffer (10 mM Tris•Cl pH 7.4). D1 selections were pooled together for sequencing.

### 1.4 Selection output quantification, factor analysis with Boruta and Lasso regression

Recovery and innest PCR products from each selection were quantified through densitometric measurement of band intensity from agarose gel electrophoresis using ImageJ. Gels were pre-processed using ImageJ’s Rolling Ball background subtraction (20 pixels) method. PCR products were also quantified with by absorbance at 260 nm using the SpectroStar Nano (BMG Labtech, UK) and/or through dye-based Qubit fluorometric (Thermo Fisher Scientific) using the 1X dsDNA HS Assay Kit (Thermo Fisher Scientific). Background noise was removed by dividing gel quantifications by the average yield of negative control reactions. Spectrophotometric quantifications were noise-adjusted by subtracting the average yield of negative control reactions. Quantifications were normalized to the smallest yield (0%) and largest yield (100%) identified in each quantification method. Input data for factor analysis, containing the corresponding factors, levels and selection output quantification, can be found in S1_DOE_Analysis.xlsx. Both, Boruta and Lasso regression analysis were carried out using the DOE_Factor_Analysis.R **Supplementary Information S2** (RStudio version 4.3.1). In short, for the Boruta implementation, the Boruta package (version 8.0.0) was used and the datasets were split into predictors (factors) and responses (selection output quantifications). The Boruta algorithm was executed with 100 runs (“maxRuns”). The Lasso regression was carried out using the glmnet package (version 4.1.7) on the predictors and responses. Cross-validation was implemented to find the optimal value of the regularization parameter (*λ*) and find the best performing model. The model was iterated 100 times to obtain stable coefficient scores. The absolute average coefficient scores were used as a measure of feature importance. Performance metrics (R-squared, Mean Squared Error (MSE), Mean Absolute Error (MAE), Bayesian Information Criterion (BIC), and Akaike Information Criterion (AIC)) were also calculated. Coefficient scores and performance metrics can be found in *DOE_Analysis.xlsx* file (**Supplementary Information S3**).

#### 2.1.1 Next-generation sequencing (NGS) data pre-processing

NGS-based amplicon sequencing was carried out for all libraries by Azenta Life Sciences. The 2-point saturation mutagenesis library (theoretical library size of 400 mutants) was sequenced using the Amplicon-EZ service, expecting 400k reads pre-quality filtering. The 5 selections from Design 1 were pooled together and sequenced using the Amplicon-EZ service, expecting ∼80k sequences for each selection pre-quality filtering. The 5-point saturation mutagenesis library (theoretical library size of 3.2 million mutants) and the 3 selections from design 3 were independently sequenced using the Custom Short-Read Amplicon service, expecting 4 million reads for the input library and 2 million reads for each selection. Sequencing data, which can be found in the NCBI SRA database (BioProject: PRJNA1096033), was pre-processed using the Galaxy public server (usegalaxy.org) (Afgan et al., 2022). The total number of reads and the impact of the pre-processing workflow can be found on **Supplementary Information S1 Tables S3-S4**.

### 1.5 Calculation of enrichment and fidelity scores

Sequencing reads were trimmed at the mutation site and mutant frequencies before and after selection were calculated using the *DOE_NGS_Analysis.ipynb* Julia notebook (Julia 1.7.2) (**Supplementary Information S4**). Enrichment scores and the E-test for comparing two Poisson means (Krishnamoorthy and Thomson, 2004) were calculated using the same script by implementing the following equation:

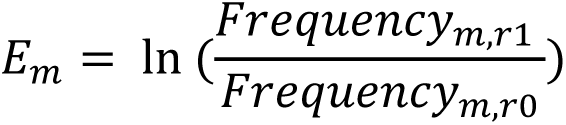

where *m* corresponds to unique mutants and r0 and r1 pre- and post-selection respectively. Calculation of overall and mutant-specific insertions, deletions and substitutions were performed using the *DOE_NGS_Analysis.ipynb* Julia notebook (Julia 1.7.2) (**Supplementary Information S4**). In short, the code aligns each read, compares it to the reference sequence and tallies the number of errors by type. The total error number is then divided by the number of bases analyzed. For mutant-specific fidelity, the same approach is implemented on mutant-specific sequence subsets. All the calculated scores can be found in the *DOE_Analysis.xlsx* file (**Supplementary Information S4**). Correlation and statistical analysis were carried out in GraphPad Prism (version 10.1.1).

### 1.6 Expression of polymerase variants for PCR with 2’F-dATP

The mutants: m1 (L403F), m2 (L403V), m3 (D404S), m4 (L403V; D404S), m5 (Y409S), were constructed on the pET23-KOD-Exo- (Supplementary Information S1 and Table S2). Mutants m1-m4 were generated using the forward mutagenic primers: KOD_L403F_fw, KOD_L403V_fw, KOD_D404S_fw, KOD_L403V-D404S_fw respectively, together with reverse primer KOD_mut_rv1 (Supplementary Information S1 and Table S1). Mutant m5 was generated with forward mutagenic primer KOD_Y409S_H5 and reverse primer KOD_mut_rv3 (Supplementary Information S1 and Table S1). Mutants were generated through inverse PCR (iPCR) on the pET23-KOD-Exo-(Table S2) plasmid in Q5 High-Fidelity DNA Polymerase (NEB) reactions of 30 cycles following the manufacturer’s standard protocol. Amplified products were treated with 0.4U/µl DpnI for 1 h at 37°C prior to their purification with the GeneJET PCR Purification Kit (Thermo Scientific). Blunt-end ligation was carried out with 100 ng of DNA ligated in 20 µl reactions with 40U/µl T4 DNA ligase, 1U/µl T4 PNK in 1x T4 DNA ligase buffer for 2 h at room temperature. From the ligation products, 5 µl were transformed in NEB^®^ 5-alpha competent *E. coli* cells following the recommended High Efficiency Transformation Protocol (C2987, New England Biolabs) described by the commercial strain provider. Successful cloning was confirmed through Sanger Sequencing.

Mutants were re-transformed into BL21(DE3) (NEB) cells, and an isolated transformant was then grown overnight and later subcultured in 50 mL of fresh LB media. The cells were induced at OD_600_ = 0.8 for 4 h at 30°C. Cells were pelleted, frozen at -20°C and resuspended in 4 mL/gr of pellet of B-PER Reagent (Thermo Fisher Scientific) with 2 µl of lysozyme (50 mg/mL, Sigma-Aldrich) and 2 µl of DNase I (2500U/mL, NEB) for each mL of B-PER reagent used, and 1 mM of Pefabloc (Sigma-Aldrich). Cells were incubated for 15 min at room temperature. Lysed cells were centrifuged at 15,000 *g* for 5 min and soluble fraction was isolated and resuspended in 50 mM of Tris•HCl (pH 7.9), 50 mM of KCl, 0.1 mM of EDTA, 1 mM of DTT, and 0.5 mM of PMSF. Resuspended cells were incubated at 80°C for 30 min, pelleted, and cleared supernatant isolated. Purified protein was concentrated using an Amicon Ultra-4 Centrifugal 50 kDa Filter Units (Sigma Aldrich NV). Protein was quantified against a standard curve made with Bovine Serum Albumin (BSA). PCR reactions with standardized protein concentrations were carried out using the CSR_Sel_SHORT_F3/R3 (Supplementary Information S1 and Table S1) primers and corresponding reaction additives (Supplementary Information S3). The thermocycling parameters used were: 35 cycles of 1 min at 94°C, 1 min at 61°C and 80 s at 68°C.

### 1.7 Sequencing coverage analysis

The sequencing coverage analysis was carried out using the *DOE_NGS_Analysis.ipynb* Julia notebook (Julia 1.7.2) (**Supplementary Information S4**). Here we define coverage as:

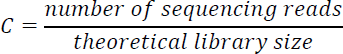

In short, random sets of sequences within a range of sequencing coverages (0.1, 0.2, 0.5, 0.8, 1, 2, 5, 10, 20, 50, 60) for *N* trials (typically 10, with replacement) are extracted from a pre-(R0) and post-(R1) selection datasets. For each pair of random sets of sequences, the enrichment analysis described earlier is carried out. Then, we calculated the probability of isolating a mutant 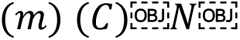 trials using the following equation:

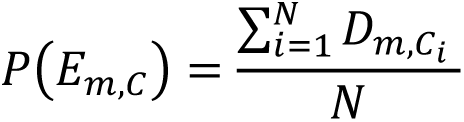

where *D_m_*,*c_i_* represents the binary outcome of significant enrichment detection of mutant *m* at the specific coverage size *C_i_* in the *i*-th trial (0 for not detected, 1 for detected) and *N* the total number of trials. Average precision calculations were carried out using the following formula:

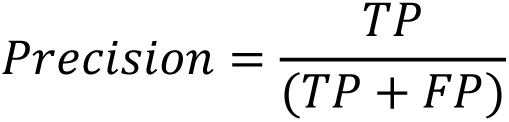

where *TP*, represents the average probability of true positives and *FP* the average probability of false positives.

## 3 Results

CSR consists of a feedback loop where polymerase libraries are expressed, and functional variants replicate their own encoding gene (self-replication) within discrete, spatially separated compartments that also contain their respective substrates. The compartmentalization, a water-in-oil emulsion, establishes a strong phenotype-genotype link, which results in better adapted mutants being able to better amplify (i.e. generate more copies of) their own genes. Next Generation Sequencing (NGS) of the libraries pre-selection and post-selection can then be used to calculate the enrichment in frequency of individual mutants and to identify beneficial (sequences that are more frequent post-selection) and detrimental (sequences that are less frequent post-selection) mutations.

To test the effect of multiple CSR parameters on the output of selection, we chose to implement a Design of Experiments (DoE) pipeline. DoE provides a systematic and efficient approach to experimental work by optimizing the allocation of resources and minimizing the number of experimental runs needed (Gilman et al., 2021). Continuous and categorial parameters suspected to influence an outcome or “response” are known as “factors”. Each factor has assigned values, or “levels”, based on *a priori* knowledge (Farooq et al., 2016). Figure 1A summarises CSR as well as potential experimental factors including library size, nucleotide chemistry, the concentration of cofactors and additives, and the reaction time.

There are several responses with varying levels of complexity that can be analyzed to determine whether a selection round was successful and/or efficient in isolating the desired phenotype. Since in CSR, active variants replicate their own encoding genes, selection efficiency can be estimated from the recovered product yield. While this quantification will not provide precise information of which variants were isolated, it can provide an overall view of the effect of a factor on polymerase activity at the population level. When recovered products are sequenced, mutant identification and abundance quantification pre- and post-selection can be used to determine enrichment scores which can be used as a proxy for polymerase replication efficiency. Mutant quantification, in turn, can be used to determine if selection was successful in partitioning the library, but one must know in advance which mutants are active or inactive. Alternatively, inactive variants (e.g., mutants harboring a deletion in their active site) can be introduced during selection and their enrichment scores used as a cutoff. Other responses that can be explored include individual mutant behavior such as fidelity.

**Figure 1.**
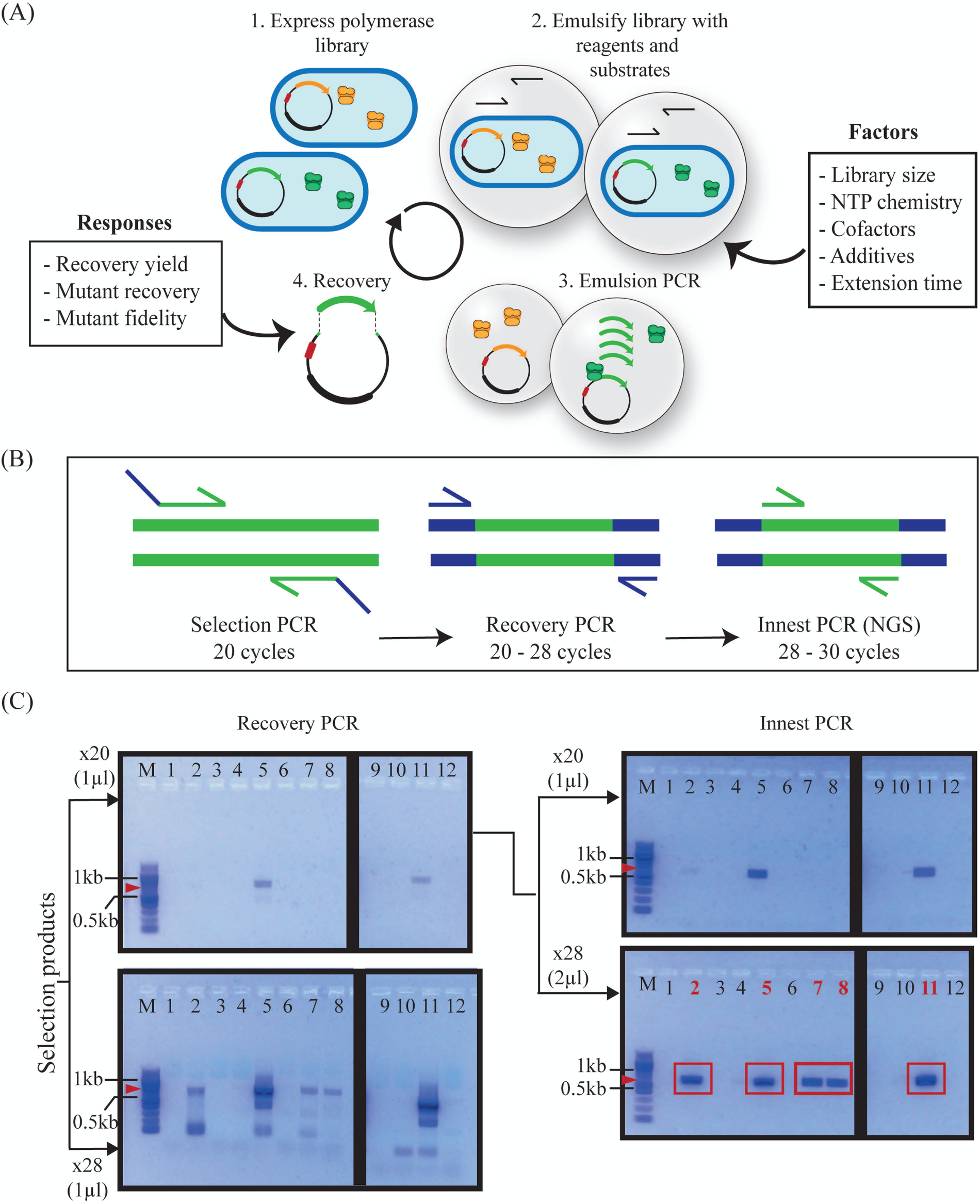
Compartmentalised self-replication (CSR) optimization pipeline and recovery. (A) An expressed polymerase library is emulsified with different substrates and reaction additives (factors). Cells are lysed within the emulsion and polymerase variants access the provided substrates to amplify their own encoding genes. The proportion of highly active variant genes within the provided selection conditions (e.g., green) should increase whereas that of variants with lower activity (e.g., orange) should decrease. Amplified products are recovered and quantified, cloned and/or sequenced. These measurements (responses) are used to determine the efficiency of selection and determine the influence of factors on selection. (B) Pipeline for recovering selection products, including recovery PCR using primers that hybridize to overhangs introduced by selection primers and subsequent innest PCR reactions to introduce overhangs for cloning or to prepare amplicons for NGS. PCR parameters were empirically optimized to minimize the number of cycles needed for visualization, while maintaining significant differences to background (control reactions). (C) Agarose gel electrophoresis of selection products from DoE Design 1 (D1) post-recovery PCR and innest PCR with different cycling parameters. The x28 innest PCR reactions amplified from recovery PCR of x20 cycles were selected for cloning and NGS as these parameters lead to maximum yields with minimum background. Red arrows indicate the expected molecular weight of the PCR product (664 bp), and red rectangles denote correctly sized products (664 bp).

To optimize and analyze CSR through DoE efficiently, it is crucial to confirm whether the expected genotypes of functional variants have been enriched (and non-functional variants depleted). By incorporating prior knowledge of the expected enriching and depleting genotypes, we can streamline the assessment of selection outcomes and refine selection parameters accordingly. Optimization of selection parameters for small, targeted libraries can be directly ported to larger libraries, maximising the likelihood of a successful selection.

To achieve this streamlined search, we constructed two polymerase libraries of different sizes, a 2-point saturation mutagenesis library of 400 variants and a 5-point saturation mutagenesis library of 3.2x10^6^ variants, targeting the catalytic site and neighboring residues. Both libraries are within the sampling capacities of CSR, where, typically, 10^8^ to 10^9^ PCR-competent compartments per milliliter are generated. The 2-point and 5-point saturation mutagenesis libraries were constructed on an exonuclease deficient KOD DNAP (D141A/E143A) background (KOD exo-). D404 (catalytic aspartate) and neighboring L403 were targeted in both libraries. F405, L408 and Y409 (involved in substrate discrimination (Cozens et al., 2012)) were additionally targeted in the 5-point library. Since D404 is the only optimal solution for polymerase functionality at that residue, recovery of this genotype functions as a benchmark for selection efficiency.

### 3.1 Uncovering key factors: initial screening and main effects analysis

A typical DoE pipeline, when no prior knowledge of a process is available, starts with a screening phase, where multiple factors are explored assuming a simple linear model to determine the main effects, if any, of each factor on the response being analyzed (Antony, 2023). The factors can then be narrowed down for further characterization where more models can be tested (e.g. factor interactions and quadratic effects). A final optimization stage can then be implemented to identify more complex interactions between factors that can further optimize the process (Antony, 2023).

An initial DoE-guided compartmentalized self-replication campaign, hereinafter referred to as D1, was designed following a D-optimal linear model with 9 factors (nucleotide, magnesium, manganese, BSA, betaine, PEG1000 and formamide concentrations, extension time and nucleotide chemistry) of 2 levels (Table 1). With this model, we created a design comprised of 12 distinct selection experiments to screen the main effects of these factors. The full list of parameter combinations can be found in **Supplementary Information S3**. The 12 unique reaction mixtures were prepared and used to resuspend a mixture of 2x10^8^ induced cells containing 90% of the 2-point library and 10% of a non-functional 91 amino acid deletion variant of KOD DNAP, KODΔ. Recovery of KODΔ (391 bp) can easily be identified from real selection products (664 bp) through agarose gel electrophoresis following PCR. KODΔ recovery can highlight unsuccessful selections or excessive amplification steps leading to a high accumulation of background. The 12 selections were carried out and selection products were extracted and purified.

**Table 1.**
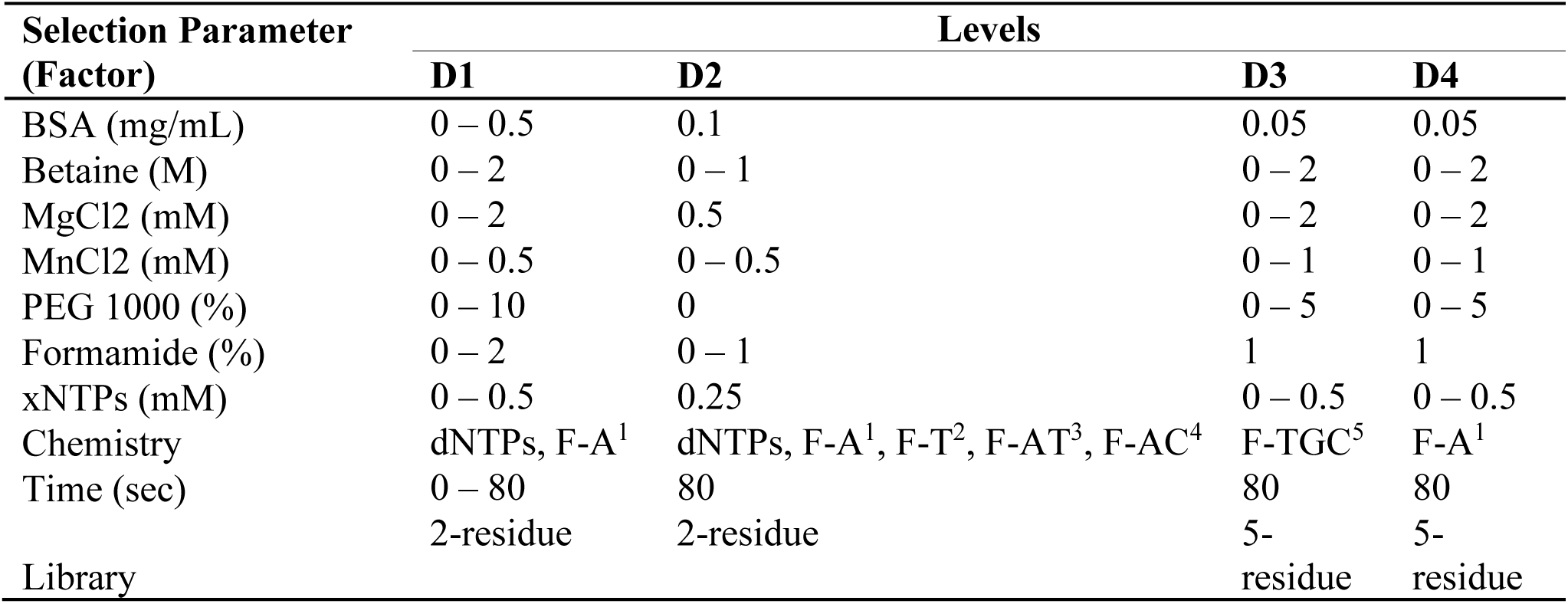
Summary of CSR factors and levels tested in each design. ^1^F-A: 2’F-dA, dT, dG, dC; ^2^F-T: 2’F-dT, dA, dG, dC; ^3^F-AT: 2’F-dA, 2’F-dT, dG, dC; ^4^F-AC: 2’F-dA, dT, dG, 2’F-dC; ^5^F-TGC: dA, 2’F-dT, 2’F-dG, 2’F-dC. Detailed description of each design can be found in Supplementary file S3.

The selection and purification steps typically result in low product yields and often at concentrations that cannot be accurately determined. Thus, an initial recovery PCR step is carried out using primers that hybridize to regions introduced by overhangs in the selection primers (Figure 1B) using Taq DNAP (NEB) as it is capable of amplifying from single copies of template. Since Taq DNAP has lower fidelity than other commercially available polymerases we limit the cycle number to 20 – 28 cycles. An in-nest PCR, using KOD Xtreme (an ultra-high fidelity DNAP), is then carried out on purified recovery products to further amplify the signal and generate amplicons for cloning or for NGS with primers that hybridize within the gene. As shown in Figure 1C, 28-cycle recovery PCRs result in the recovery of the KODΔ mutant as well as other low molecular weight products. Thus, we proceeded with the products from 20-cycle recovery PCRs and further amplified them with in-nest PCRs. Again, we tested different cycling parameters and input template concentrations and found that 28 cycles and 2 µl of input template enabled the successful recovery of 5/12 selections.

Densitometric measurement from agarose gel electrophoresis and spectrophotometric measurements of recovered DNA, corresponding to relative amplicon quantity, were obtained for each recovery PCR (Rec) or in-nest PCR (Innest) reactions with either 20 or 28 cycles. The relative recovery of each reaction with either gel electrophoresis (G) or spectrophotometer (S) quantification is shown in Figure 2A. Selections 2, 5, 7, 8 and 11 yielded products across all quantifications. Products from selections 7 and 8 were not detected with the Rec20G or Innest20G quantification. The parameters of these positive selections are shown in Figure 2B.

By analyzing and interpreting the variations in DNA recovery, we evaluated the relationship between selection parameters and selection efficiency (the efficiency of isolating highly active variants). High recovery yields indicate the successful isolation of a highly active population, while low recovery suggests poor recovery of active variants. The pattern and quantification of recovery across multiple selection conditions is what guides the identification of important factors in selection.

**Figure 2.**
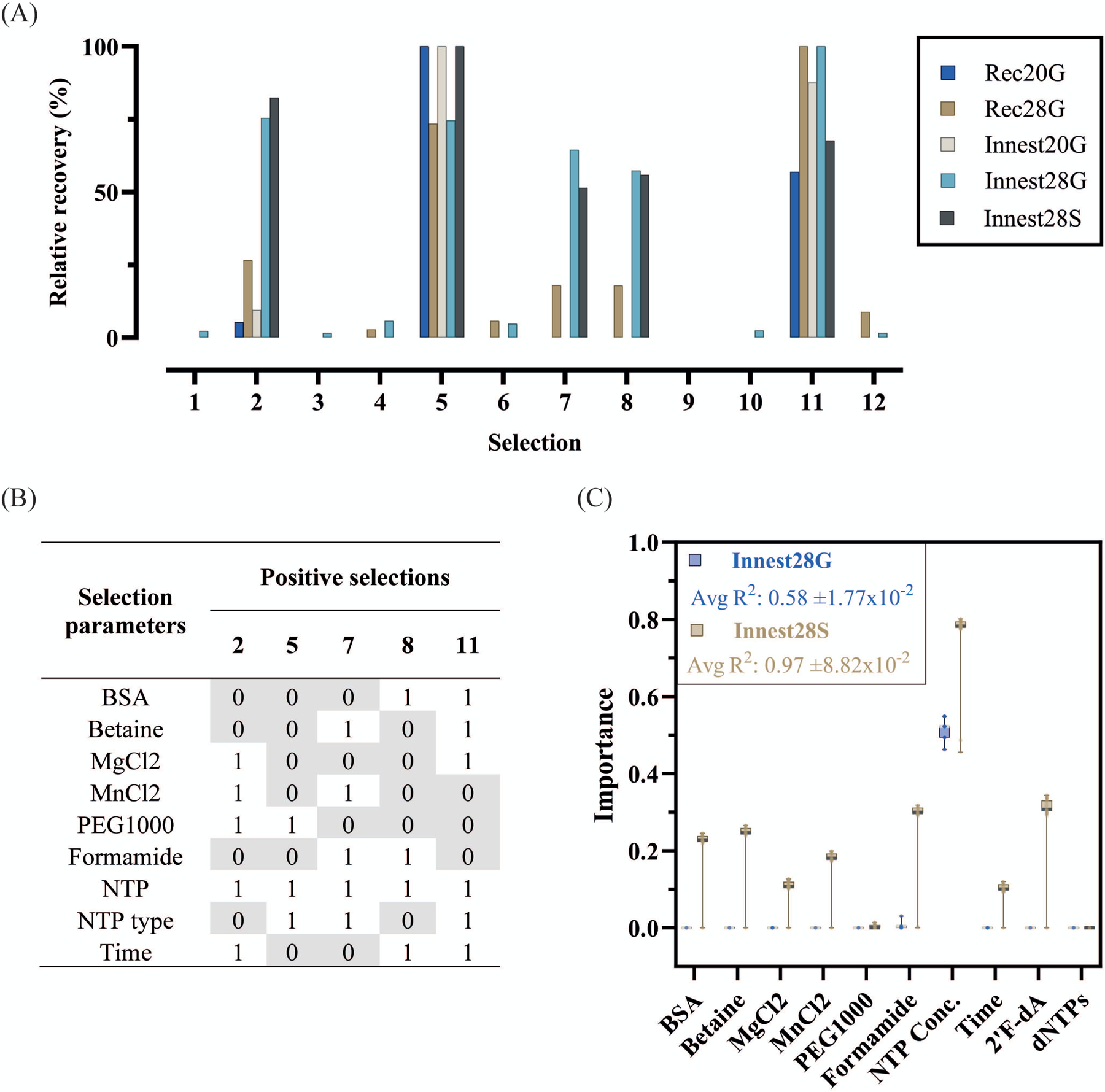
Quantification and analysis of DoE-CSR D1 selection products. (A) Selection products from recovery (Rec) and innest (Innest) PCRs with x20 (e.g., Rec20) or x28 (e.g., Rec28) cycles, were quantified through densiometric measurement of band intensity from agarose gels (G, e.g. Rec28G) and through spectrophotometric measurements by absorbance at 260 nm (S, e.g., Rec28S). The product yields were normalized to the smallest yield (0%) and largest yield (100%) identified in each quantification method. (B) Selection parameters and their corresponding binary level (0 = 0 concentration or minutes, 1 maximum concentration or minutes) of positive selections from D1 (Selections 2, 5, 7, 8, 11). The complete list of factors and levels for all selections and actual concentrations/times can be found on supplementary file X. (C) Feature importance analysis was carried out using Lasso regression model to determine the impact of selection factors on Innest28G and Innest28S responses. The coefficient values of each factor derived from 100 runs of the model were averaged and their absolute value plotted as a measure of factor importance. The average R^2^ and RMSE values for the models with Innest28G as the response were 5.8 ±0.017 and 0.62 ±0.013 respectively and the average R^2^ and RMSE values for the models with Innest28S as the response were 0.97±0.088 and 0.14±0.11 respectively.

Initially, we applied the Boruta algorithm to identify relevant factors for DNA recovery post-selection. Boruta is a well-established wrapper method in machine learning, based on random forest classification to identify and remove irrelevant features in datasets (Kursa et al., 2010). The analysis, conducted on all available samples, aimed to determine the recovery stage with the highest predictive power and the earliest identification of factors in the selection pipeline.

No significant factors were identified for Rec20 and Innest20 samples and we observed significant variability on the Boruta feature selection for these samples. This suggests that the DNA quantification methods were not sensitive enough for the 20-cycle samples. For Rec28, Innest28G and Innest28S, only nucleotide concentration was identified as significant. While the additional number of PCR cycles and purification steps can reduce the precision, these provided a more robust and stable signal, enabling better Boruta algorithm performance.

To cross-validate Boruta findings, we employed Lasso regression modeling, which automatically selects relevant features while preventing overfitting and handling multicollinearity (Muthukrishnan and Rohini, 2017). Figure 2C displays absolute coefficient scores of each feature’s importance with Innest28G and Innest28S responses, with detailed scores in **Supplementary Information S3**.

Using the Innest28G response, nucleotide concentration positively impacted recovery, while formamide had a slight negative effect. In contrast, Innest28S showed a better fit, identifying BSA, betaine, magnesium, manganese, and formamide concentrations, as well as 2F-ATP presence, as negatively influencing selection yield. Nucleotide concentration and time positively affected recovery, while PEG1000 was not a significant predictor.

Overall, nucleotide concentration was consistently identified as the only significant factor that contributed positively to selection recovery efficiency. Because of the nucleotides naturally present in cells, CSR selections can yield amplification even in the absence of exogenous dNTPs. Variants that can make use of the reduced dNTP concentrations are seen as “parasites” in XNA synthesis; they are difficult to track and to remove from the population. By introducing a “zero-concentration” of nucleotides, we hoped to identify parameters that benefit parasites. Instead, these extreme conditions may have overshadowed the effects of other factors - oversimplifying the underlying problem. Nonetheless, D1 was still useful in reducing the range of factors for further optimization.

### 3.2 In-depth factor analysis and interaction exploration

In the next design, D2, we decided to fix nucleotide concentration at 0.25 mM as in the original CSR protocol and reduce the number of factors to manganese, betaine, and formamide concentrations and nucleotide chemistry (dNTP mixtures with 2F-ATP, 2F-TTP, 2F-ATP/2F-TTP or 2F-ATP/2F-CTP substitutions). D2, was generated using response surface methodology (RSM) with a central composite design (CCD) (Table 1).

The 36 distinct selection experiments of D2 were chosen to explore main effects and 2-way interactions of the 4 factors. Instead of a KODΔ spike, for D2 we opted to use a “negative control selection” - selection reactions prepared without exogenous nucleotide triphosphates. As with the initial analysis, 28-cycle recovery PCR (Rec28G) and in-nest PCR (Innest28S) were used for quantification (Figure 3A).

**Figure 3.**
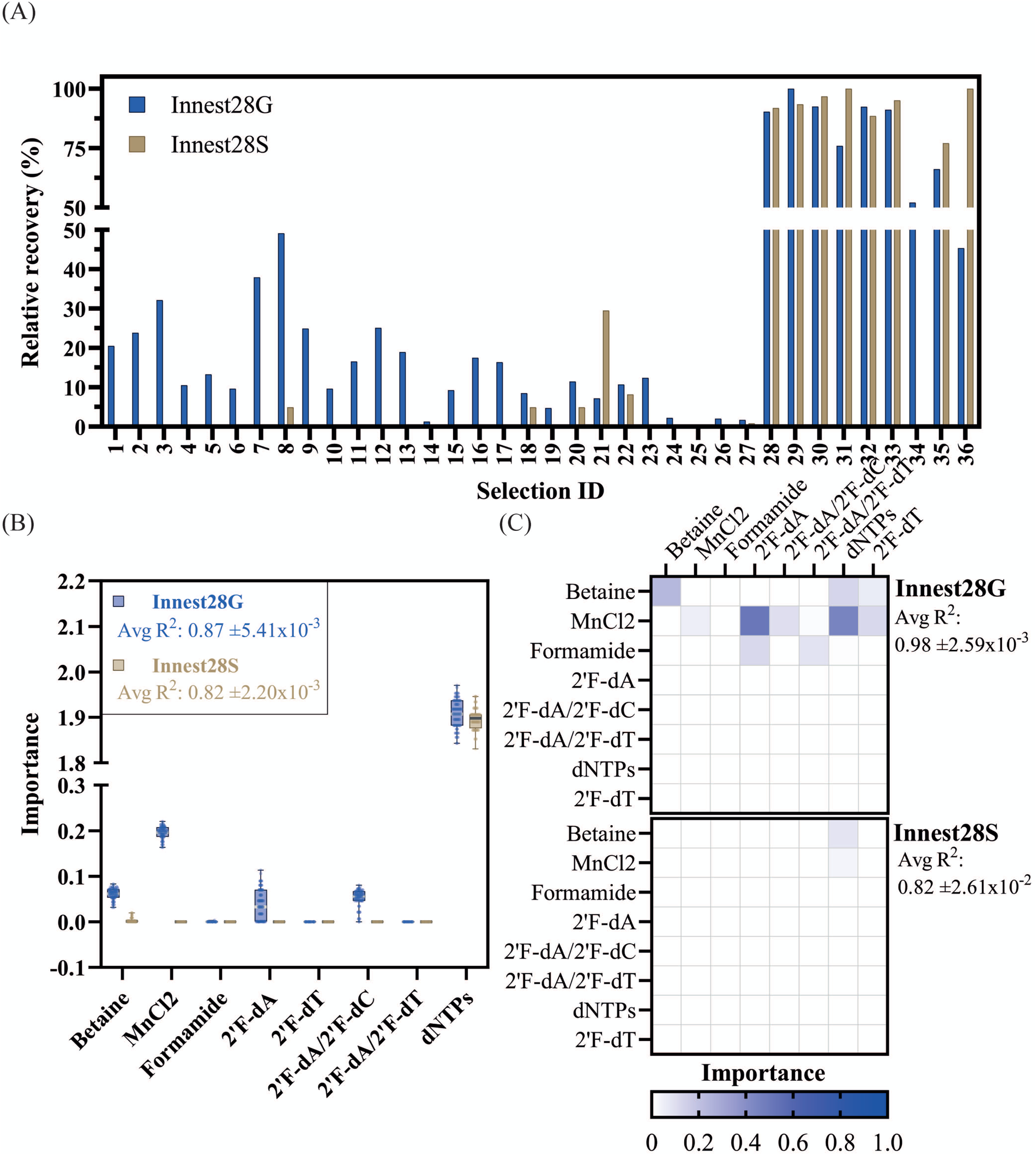
Quantification and analysis of DoE-CSR D2 selection products. (A) Products from the x28 innest PCR reactions from the D2 selections were quantified through densiometric measurement of band intensity from agarose gels (G) and through spectrophotometric measurements by absorbance at 260 nm (S). Background noise was removed by dividing gel quantifications by the average yield of 12 negative control reactions. Spectrophotometric quantifications were noise-adjusted by subtracting the average yield of the 12 negative control reactions. Quantifications were normalized to the smallest yield (0%) and largest yield (100%) identified in each quantification method. (B) Feature importance analysis was carried out using a Lasso regression model to determine the impact of selection factors on each response (Innest28G or Innest28S). The coefficient values of each factor derived from 100 runs of the model were averaged and their absolute value plotted as a measure of factor importance. The average R^2^ and RMSE values for the models with Innest28G as the response were 0.87±0.0054, 0.35±0.0074 respectively and the average R^2^ and RMSE values for the models with Innest28S as the response were 0.82±0.0022, 0.41±0.0026 respectively. (C) Lasso regression modelling with interaction terms was carried out to identify factor interactions. The corresponding feature importance metrics are displayed in a 2D plot for clarity. The average R^2^ and RMSE values for the models with Innest28G as the response were 0.98±0.0026, 0.15 ±0.0079 respectively and the average R^2^ and RMSE values for the models with Innest28S as the response were 0.82±0.026, 0.41±0.0026 respectively.

To explore potential interactions across factors and understand their impact on the response variable, we employed a Lasso regression analysis. When examining the main effects of each feature with the Rec28G response variable, several factors appeared to have a positive influence on selection yield (Figure 3B). Notably, the concentration of betaine and the inclusion of fully natural nucleotides showed a significant positive effect on the yield, as well as, to a lesser extent, the presence of 2’F-dA in the nucleotide mix. Conversely, the presence of manganese and the combination of 2’F-dA/2’F-dC in the nucleotide mix had a notable negative impact on the yield.Analysis of Innest28S reactions identified some of the same correlations. However, model homoscedasticity and normality of residuals were suboptimal, suggesting a poorer fit than Rec28G dataset.

In our pursuit of further model refinement, we sought to enhance the understanding of quadratic effects and interaction terms. Interestingly, for the Rec28G response variable, we additionally identified the positive effects of the interaction between betaine and natural substrate incorporation, betaine and 2’F-dT incorporation, manganese and 2’F-dT incorporation and a positive quadratic effect of betaine on recovery yield, indicating a non-linear relationship. Additionally, we discovered a negative effect of formamide as well as its interaction with 2’F-dA and the combination of 2’F-dA/2’F-dT and a negative quadratic effect of manganese as well as its interaction with 2’F-dA, 2’F-dA/2’F-dC, dNTPs incorporation.

Formamide is a widely used organic PCR additive that lowers melting temperature by binding to the major and minor DNA grooves but has a narrow effective range and can lead to reduced amplification in high concentrations (Chakrabarti and Schutt, 2001). In D2 we tested 0, 0.5% and 1% concentrations, which are below the recommended concentrations (1.25-10%) (Sarkar et al., 1990). Nonetheless, formamide interactions with the emulsions used are not well characterized. For instance, if formamide weakens compartment integrity, it would compromise selection by allowing exchange of genetic information between compartments. Interestingly, manganese also appeared to have a negative effect on selection. Family B polymerases can use Mg^2+^ and Mn^2+^ to catalyze nucleotidyl transfer and presence of Mn^2+^ has been shown to promote the incorporation of non-natural nucleotides (Chen et al., 2016; Dunn and Chaput, 2016; Kropp et al., 2017; Pinheiro et al., 2012). In family B-RB69 DNAP-catalyzed reactions, Mn^2+^ enhances ground-state binding of both correct and incorrect dNTPs and promotes misincorporation by increasing the rate of incorporation (k_pol_ or V_max_), reducing substrate selectivity and fidelity (Vashishtha and Konigsberg, 2016). Thus, while it was expected that the presence of manganese ions would result in increased recovery yield, these results indicate that its inclusion, at the tested concentrations, may hinder selection efficiency.

Overall, the analysis captures some of our expected results: higher levels of substitution interfere with enrichment and that betaine works as a PCR enhancer in the selection conditions.

### 3.3 Mutant enrichment and fidelity analysis of D1 positive selections

The balance between exonuclease and polymerase active sites as well as the translocation process are dynamic processes that can be altered through directed evolution (Kuroita et al., 2005; Pinheiro, 2019). Therefore, we sought to explore the influence of selection parameters on mutant enrichment and mutant fidelity, offering insights into the overall polymerase incorporation rate and exonuclease activity, respectively.

In parallel with the detailed factor analysis, we deep sequenced successful selections from D1 for analysis. All 400 variants out of the 400 possible 2-point mutant variants were identified in the library pre-selection. Post-selection, 229 unique variants were isolated across all selections (Figure 4A). Of these, 218 variants show a significant change in abundance with only 19 variants showing significant enrichment across all selections (FD, MD, ID, HD, YD, QD, VD, CD, KD, SD, LD, AD, ED, ND, DD, TD, WD, PD, GD) (Figure 4B). As expected, catalytic D404 is an essential residue for enrichment and residue 403 can tolerate variation. LE showed significant enrichment in 2 selections, albeit to a notably lesser extent than the top 19 mutants with the catalytic D404 reversal. Notably, the wild-type sequence L403 does not consistently provide the highest enrichment in the different selection conditions, with L403F mutant consistently outperforming the wild-type in all tested selection conditions.

**Figure 4.**
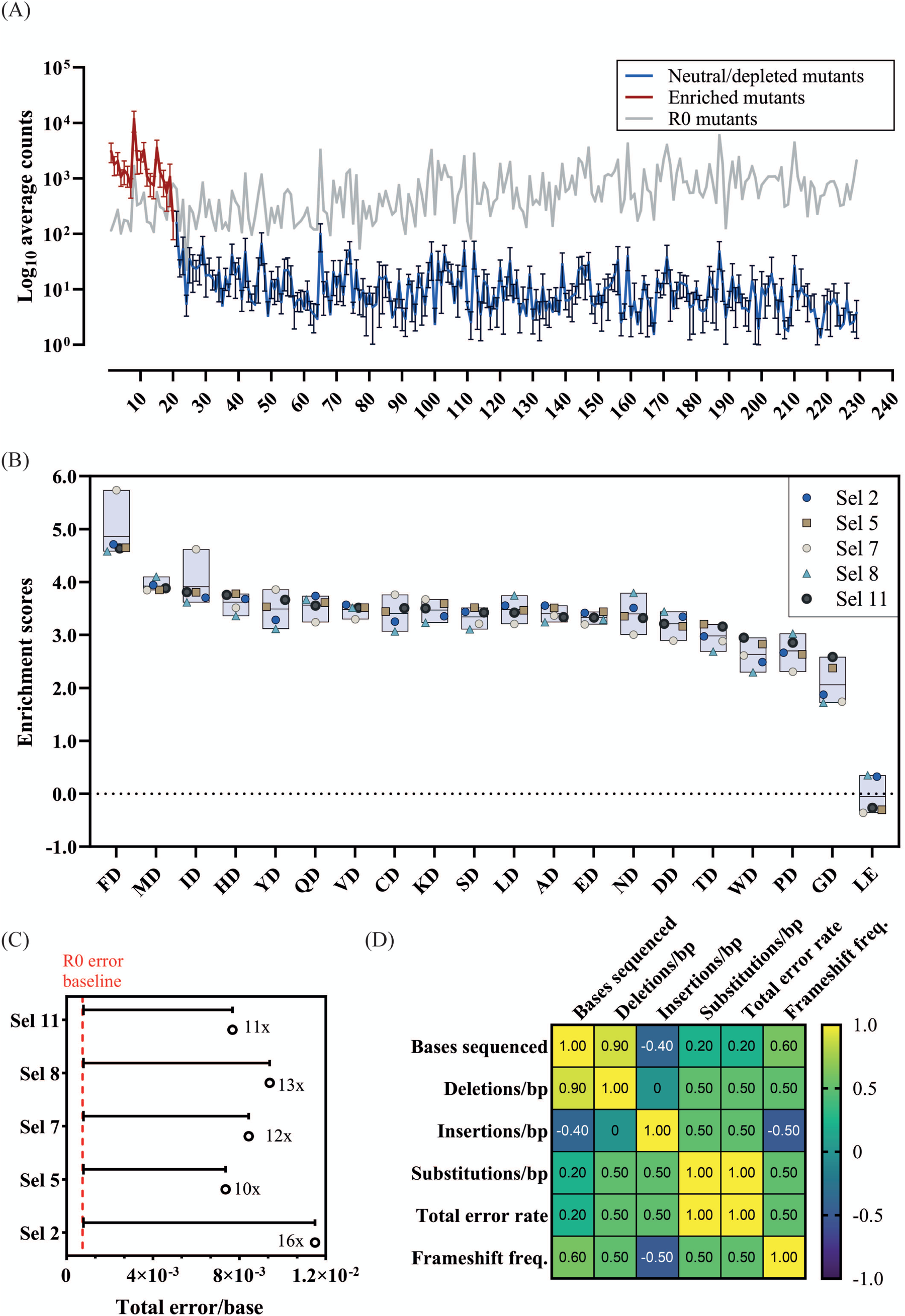
Mutant enrichment and population fidelity analysis of successful D1 selections. (A) The log average count and standard deviation of each mutant across all 5 positive D1 selections was plotted. Significantly enriched mutants are colored in red, neutral, or significantly depleted mutants are colored in blue, and the corresponding log count of each mutant pre-selection (R0) is shown in grey. (B) ) The enrichment scores of significantly enriched mutants across selections. (C) Overall fidelity scores (insertion, deletion, and substitution rates) by polymerase variants in each selection normalized to the fidelity pre-selection (R0), 7.49x10^-4^. The fold numbers correspond to the fidelity relative to the R0. (D) Correlation plot of bases sequenced and error rates by type.

To investigate the impact of selection parameters on mutant fidelity we took advantage of the principle of CSR, where each mutant amplifies their corresponding gene. Thus, we calculated the error rate of each mutant by quantifying the number of errors on their source read. The resulting error rates were normalized to those of the R0 to account for sequencing errors.

**Table 2.**
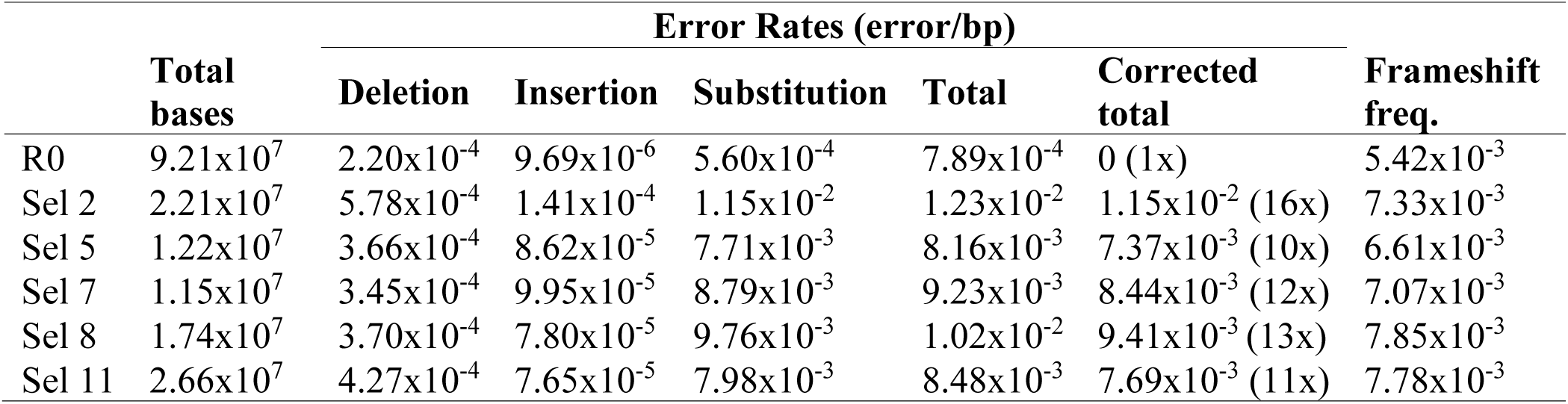
Quantification of deletion, insertion, and substitution rates (errors/bp) of KOD DNAP mutants across each selection from D1. Individual and total error rates do not consider PCR and NGS error rates. Corrected error rates are calculated by subtracting the total error rate pre-selection (R0) from the total error rates of each selection. Frameshift frequency (freq.) is calculated as the number of sequences that insertions or deletions that lead to frameshifts divided by the total number of sequences analyzed.

As shown in Figure 4C and Table 2, the error rate of Sel 2 was the highest (1.15x10^-2^ errors/base), followed by selection 8 (9.41x10^-3^). These two selections contained the 2’F-dATP substrate in the nucleotide mix. Those error rates are significantly higher than for the other 3 selections. Both mutant-independent analysis using a Kruskal-Wallis’ test and a mutant-specific Friedman test, confirm that the error rates of selection 5 (7.37 x 10^-3^), 7 (8.44x10^-3^) and 11 (7.69x10^-3^), which only contained natural substrates, are significantly different (p<0.01) to selections 2 and 8.

To rule out that sequencing depth was not affecting the fidelity estimates, we looked at a Spearman correlation between the bases sequenced and the different errors observed (Figure 4D). Insertions and deletions, being rarer, show a clear correlation with sequencing depth. Comparing the number of bases sequenced with substitution rates or the total error rates shows only a weak correlation (0.2).

The frequency of sequences that contained insertions and deletions (InDels) resulting in frameshifts that were excluded from the general InDel counts were also analyzed. A negative correlation between sequencing coverage and insertion incorporation (-0.40) as well as a strong positive correlation between sequencing coverage and deletion incorporation (0.90) were identified. Nonetheless, InDels correspond to a small proportion of the errors identified, and thus the overall error rate does not appear to be significantly correlated with the sequencing coverage.

Following the library-level analysis, we also focused our analysis on the error profile of the top 20 most-enriched variants (Figure 4B). Enriched variants are well-sampled; thus, the focused analysis should minimize the impact of selection background and NGS errors.

The patterns observed at the library level were recapitulated in the focused analysis (Figure 5A). Therefore, all subsequent analysis focused on the 20 enriched variants.

**Figure 5.**
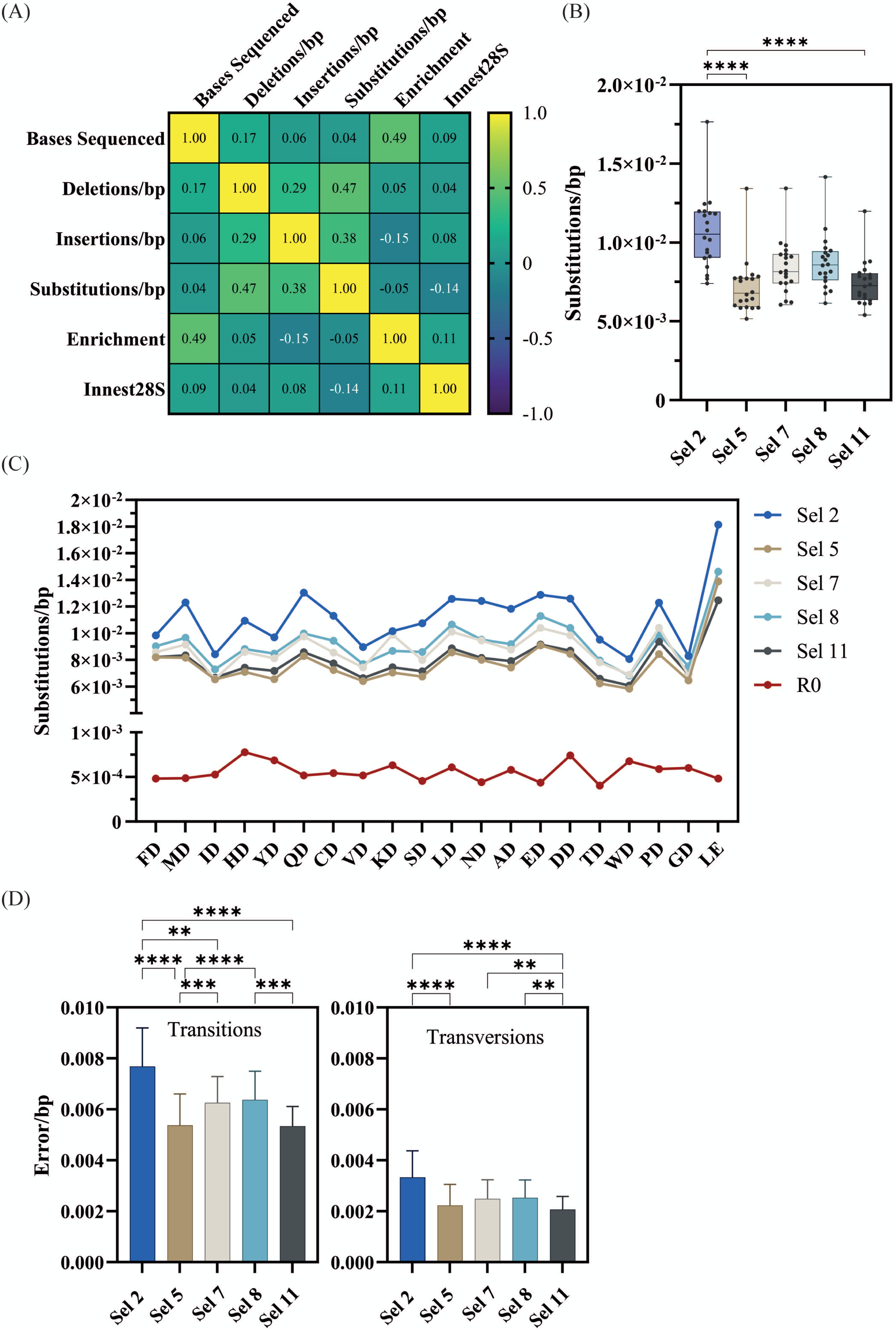
Fidelity analysis of significantly enriched mutants from successful D1 selections. (A) Correlation plot of bases sequenced error rates by type and recovery yield (Innest28S). (B) Overall substitution error rates of significantly enriched mutants across selections. Significance comparisons (p<0.01) were determined by the Kruskal-Wallis test followed by Dunn’s multiple comparison test with Bonferroni correction. (C) Mutant-specific substitution error rates across selections as well as pre-selection (R0). (D) The overall transition and transversion error rates of significantly enriched mutants across significantly enriched mutants across selections and significance comparisons determined by the Friedman test followed by the multiple comparisons Dunn’s test with Bonferroni correction.

Next, we focused further analysis on substitution rates as this was the most abundant error type. As shown in Figure 5B, mutants in Sel 2 had the highest error rates which were significantly different to those in Sel 5 and Sel 11. Figure 5C shows the substitution rates of each mutant across each selection. All mutants appear to follow a general fidelity trend, and almost identical patterns in Sel 5 and Sel 11.

Transition and transversion rates were determined (Figure 5D), identifying that C→T transitions were the most common (Figure 6), regardless of selection condition. This transition is the most common in *E. coli* (Beletskii and Bhagwat, 1996), presumably due to unrepaired cytosine deamination. A→G and T→C transitions were also found to be abundant, consistent with a study showing a bias for these transitions in WT KOD DNAP (McInerney et al., 2014). In Sel 2 and Sel 8, the preferential incorporation of dC with template thymine, could have been exacerbated due to the poor recognition of 2’F-dA and reduced stabilization of the closed conformation (Johnson, 2008; Pinheiro, 2019), resulting in the increased T→C and A→G transitions.

**Figure 6.**
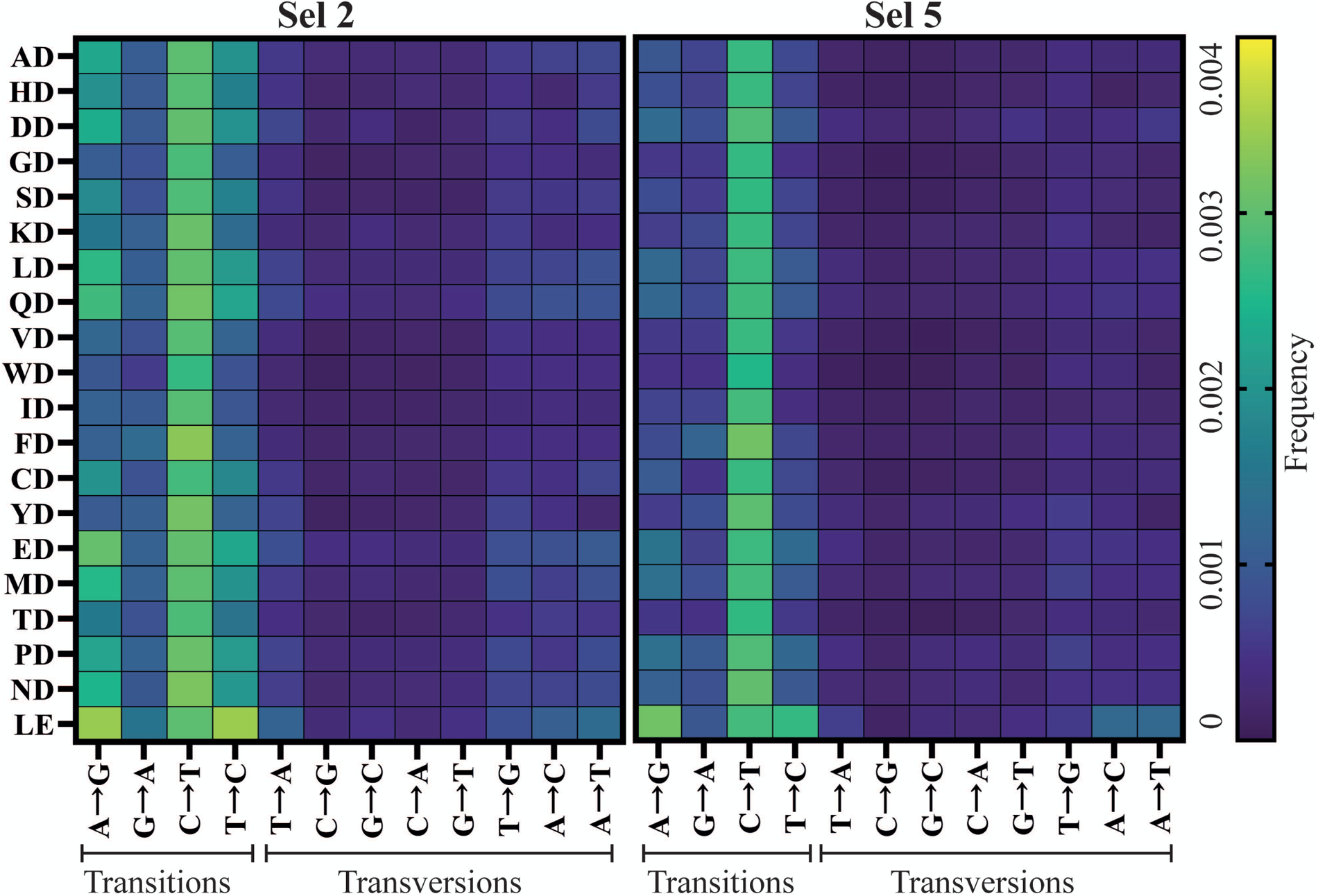
Mutant-specific fidelity analysis of top enriched mutants from successful D1 selections. Frequency of transition and transversion incorporation of selection 2 (Sel 2) and selection 5 (Sel 5) mutants.

### 3.4 Factor screening in a large sequence-functional space

Since library composition can affect the optimum selection conditions, we decided to investigate the influence of selection parameters on a library targeting 5 residues in the active site. As Y409 was targeted for mutagenesis in this 5-point library and given its “steric gate” role (Gardner and Jack, 1999; Pinheiro et al., 2012),we decided to pursue a more stringent selection: substituting 3 of the natural triphosphates for their 2’F-analogues (i.e. 2F-TTP, 2F-GTP, 2F-CTP and dATP).

We sought to assess the main effects, 2-factor interactions and quadratic effects of nucleotide, magnesium, manganese, betaine and PEG1000 in Design 3 (D3). Despite PCR cycling optimization, selection conditions were too stringent and recovery was not detectable above background.

Thus, we proceeded with Design 4 (D4) which was identical to D3 but included only one modified nucleotide (2’F-ATP) in the nucleotide mix, which had already been successfully incorporated by mutants from the 2-point mutant library in D1 and D2. 24 selections in 2 blocks of 12 reactions were carried out (**Supplementary Information S3**). Products from recovery PCR reactions of 25 cycles did not yield quantifiable amplification products but the subsequent in-nest PCR amplification of 30 cycles (Innest30Q and Innest30G) resulted in strong positive signals in 5/24 selections (Figure 7A).

We employed a Lasso regression analysis looking at the main effects of the 5 factors. With the Innest30G response variable, we observed a skew (S-shape) in the distribution of residuals and thus a departure from the optimal normal distribution, indicating that the Lasso model may not entirely accommodate potential non-linear relationships or higher-order interactions present in the data. We did not identify any relevant interactions between the factors or substantial quadratic effects in relation to any of the response variables that could improve the model. Similarly, no important features were identified with the Innest30Q response variable with a simple or more complex model. The weak relationship between selection parameters and recovery could be due to several reasons.

Firstly, the larger library size may have introduced additional complexity and variability during selection, making it challenging for the feature selection analysis to discern meaningful patterns. Additionally, while selection conditions resembled previous designs, they may have failed to sufficiently enrich active mutants in the larger library, potentially obscuring true recovery, and enrichment patterns. Finally, given the data’s complexity, a more sophisticated model might be necessary to capture underlying relationships and enhance predictive accuracy.

#### 3.4.1 Mutant enrichment and fidelity analysis of D4 positive selections

We opted to sequence the successful 5-point mutant library selections (Sel 4, Sel 8, and Sel 20). Sequence coverage for both R0 (0.5x coverage) and R1 (0.2x coverage assuming no selection) libraries was low. Nevertheless, many of the enriched variants were frequently observed (> 1000 reads) in R1. Selection 20 (0.25 mM NTP mix (2’F-dA, dT, dG, dC), 0.5 mM MgCl_2_, 0.25 mM MnCl_2_, no Betaine and no PEG 1000) showed the highest post-selection diversity and the weakest enrichment, indicating that selection stringency was low.

**Figure 7.**
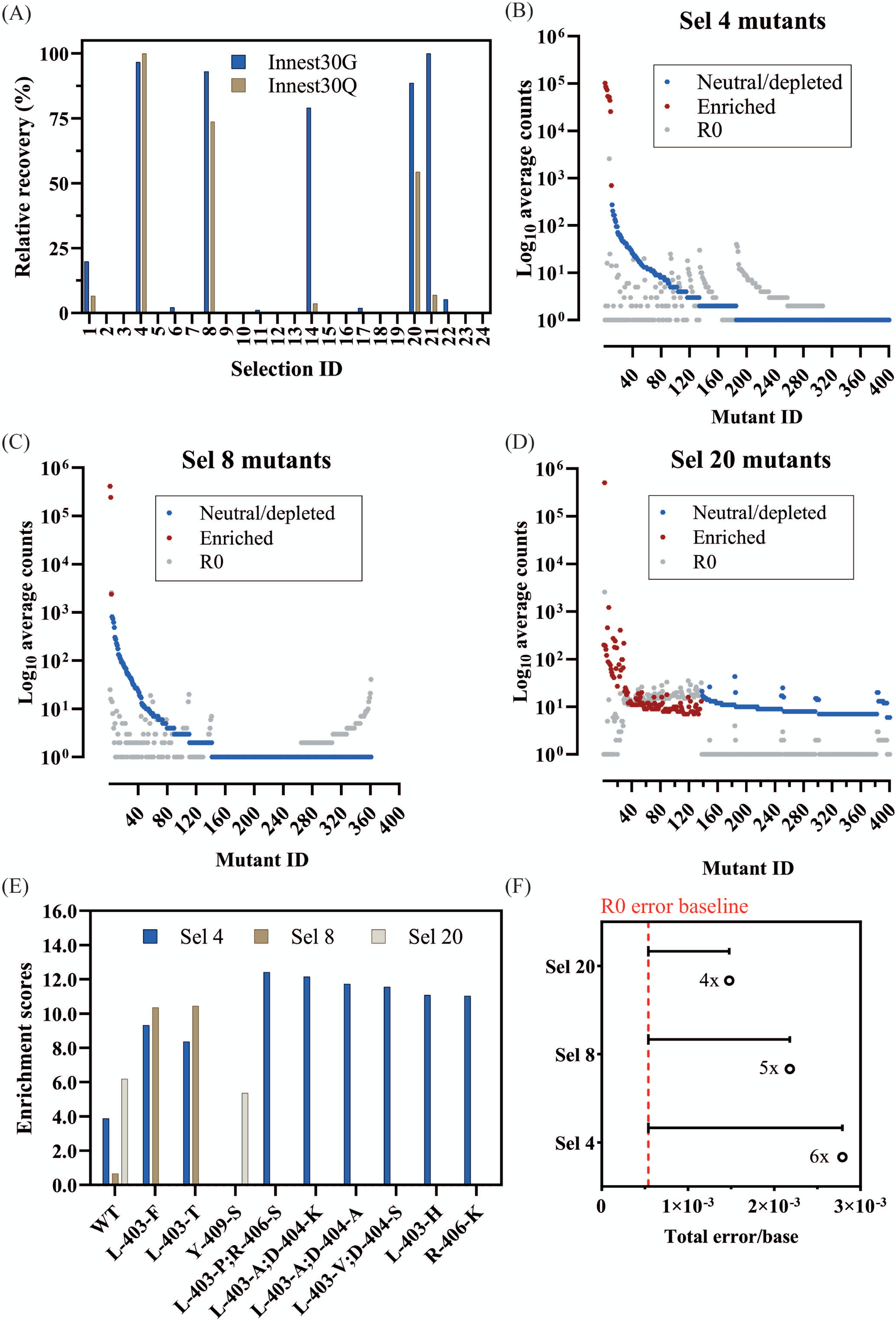
Quantification and analysis of DoE-CSR D4 selection products. (A) Products from the x30 innest PCR reactions from the D4 selections were quantified through densiometric measurement of band intensity from agarose gels (Innest30G) and through dye-based Qubit fluorometric quantification (Innest30Q). Product yields were normalized to the smallest yield (0%) and largest yield (100%). Top 400 mutant log counts from D4 positive selection 4 (B), selection 8 (C) and selection 20 (D) are shown. Significantly enriched mutants (red), neutral, or significantly depleted mutants (blue), and the corresponding log count of each mutant pre-selection (grey) are highlighted. Mutants with lower counts post-selection may appear enriched if their frequency relative to the total counts is higher. (E) The enrichment scores of significantly enriched mutants across selections with >1000 counts post-selection. (F) Overall fidelity scores (insertion, deletion, and substitution rates) by polymerase variants in each selection normalized to the fidelity pre-selection (R0). The fold numbers correspond to the fidelity relative to the R0.

**Table 3.**
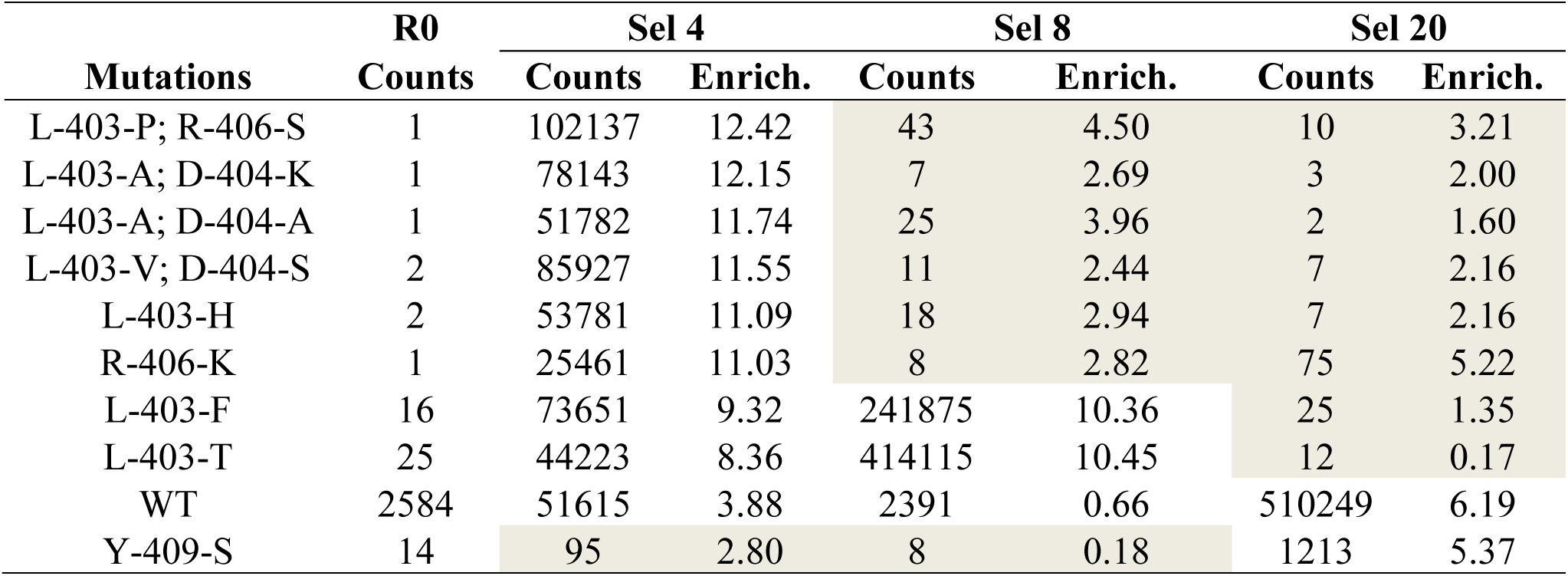
Mutants and their corresponding counts and enrichment scores (Enrich.) identified to have been significantly enriched in at least one of the positive selections from D4. The counts of each mutant pre-selection (R0) are also shown. Highlighted in tan correspond to mutants that were not identified as significantly enriched in the corresponding selection.

Due to the low sequencing coverage, analysis of mutant enrichment focused on significantly enriched mutants with >1000 reads. As shown in Figure 7E and Table 3, there was limited mutant overlap across selections, with the wild-type significantly enriched across all 3 selections. We expect this limited overlap is the result of the stringent frequency filter used in the analysis but PCR biases introduced during amplification cannot be ruled out. Mutants found in the 2-residue library are found in both selections 4 and 8 (L403F, L403T) with selection 4 conditions also isolating several frequent double mutants: L403P+R406S, L403V+D404S, and L403A+D404A. Selection 20 identified Y409S as well as a large repertoire of other significantly enriched mutants, but due to their lower observation frequency, they were not considered for further analysis.

Looking at the fidelity of the overall library (Figure 7F and Table 4), Sel 20 showed the lowest rates of misincorporation. Given Sel 20 least stringent selection conditions, the lower error may simply reflect the lack of amplification during selection.

Focusing on the fidelity of the most enriched variants (Table 3), we see mutant-specific differences (Figure 8B), with WT, Y409S and L403H showing misincorporation rates above R0 but in line with the previously determined in D1 (Figure 5C).

It is not possible to separate the impact of the low sequence coverage, selection conditions and library composition. Repeated isolation of variants in different conditions suggest that if higher sequencing coverage could be obtained, a pattern similar to the earlier experiments would be observed. Nonetheless, for this library, selection conditions from selection 8 were closest to optimum: there was clear enrichment (unlike the less stringent selection 20) and enriched sequences did not include sequences expected to be inactive (as in selection 4).

**Figure 8.**
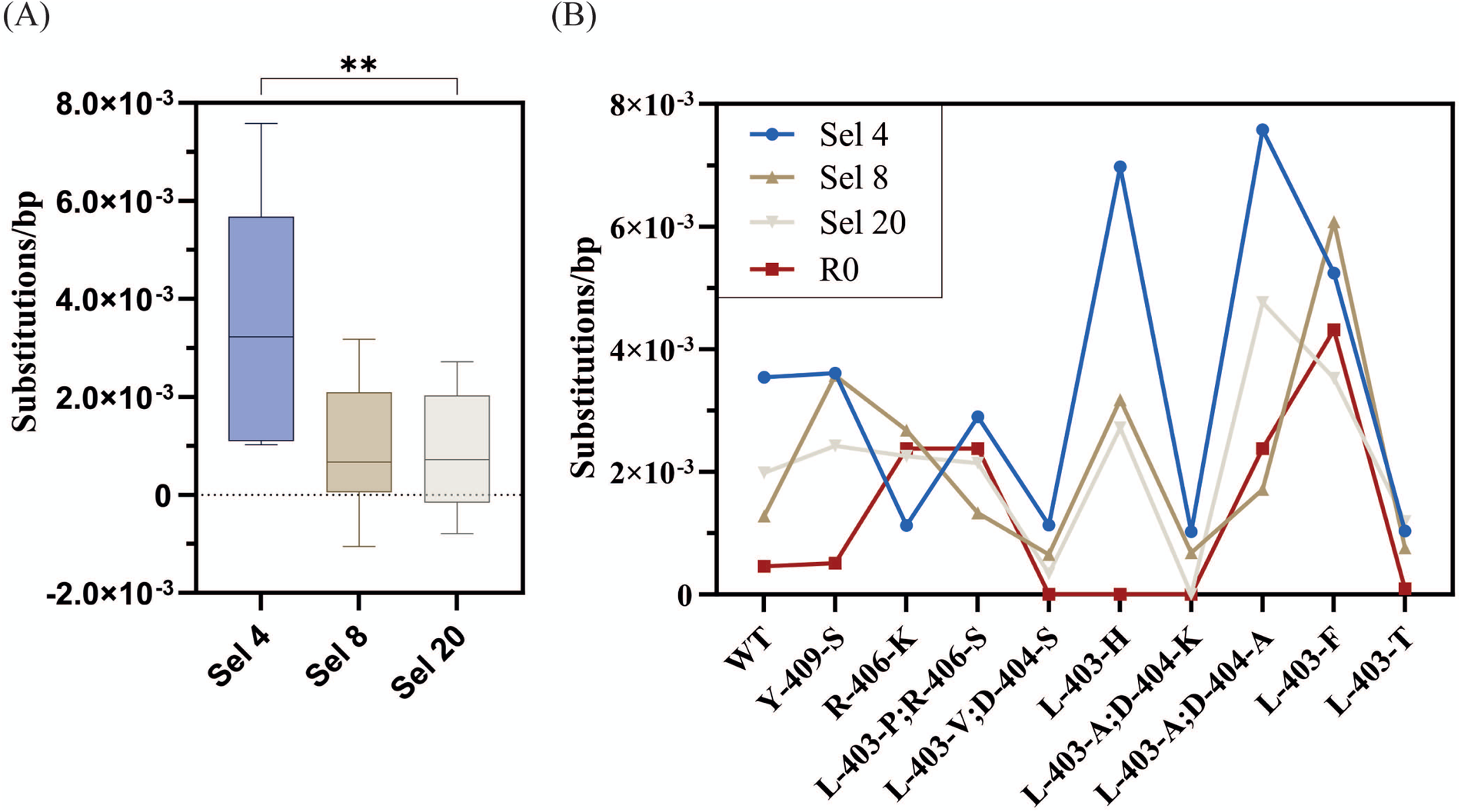
Fidelity analysis of significantly enriched mutants from successful D4 selections. The overall substitution error rates of significantly enriched mutants across selections and significance comparisons determined by the Friedman test followed by the multiple comparisons Dunn’s test with Bonferroni correction. (C) Mutant-specific substitution error rates across selections as well as pre-selection (R0).

**Table 4.**
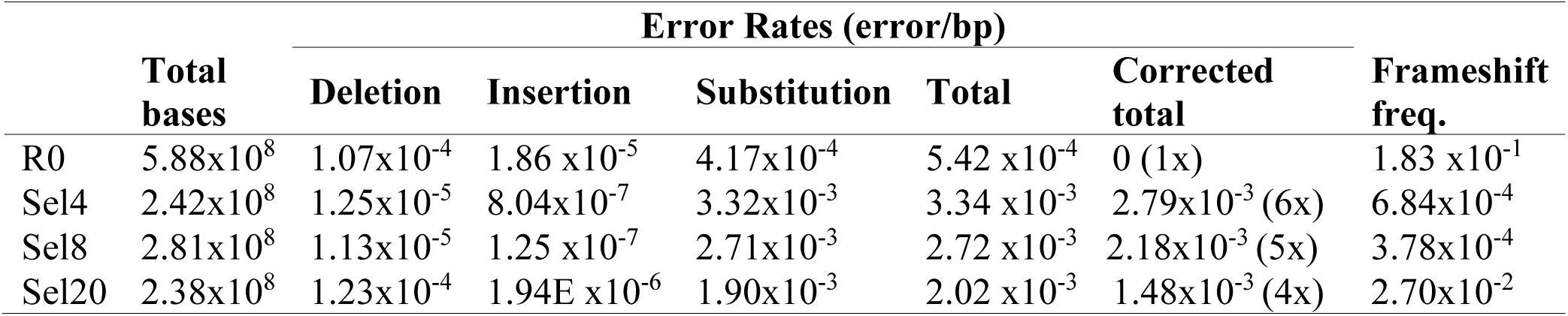
Quantification of deletion, insertion, and substitution rates (errors/bp) of KOD DNAP mutants across each selection from D4. Individual and total error rates do not consider PCR and NGS error rates. Corrected error rates are calculated by subtracting the total error rate pre-selection (R0) from the total error rates of each selection. Frameshift frequency (freq.) is calculated as the number of sequences that insertions or deletions that lead to frameshifts divided by the total number of sequences analyzed.

#### 3.4.2 Incorporation 2’F-dATP by KOD DNAP mutants in PCR

We selected a total of 5 variants for further characterization from the significantly enriched variants identified in the D4 selections (Figure 9A-B). M1 (L403F) was significantly enriched in both Sel 4 and Sel8 conditions, and one of the variants that enriched better than wild-type in D1 conditions. M4 (L403V and D404S) was identified from Sel4 but also tested as individual mutations: m2 (L403V) and m3 (D4043S). Finally, m5 (Y409S), enriched in Sel 20, was selected since it is an already characterized residue known to be involved in substrate discrimination (Cozens et al., 2012).

**Figure 9.**
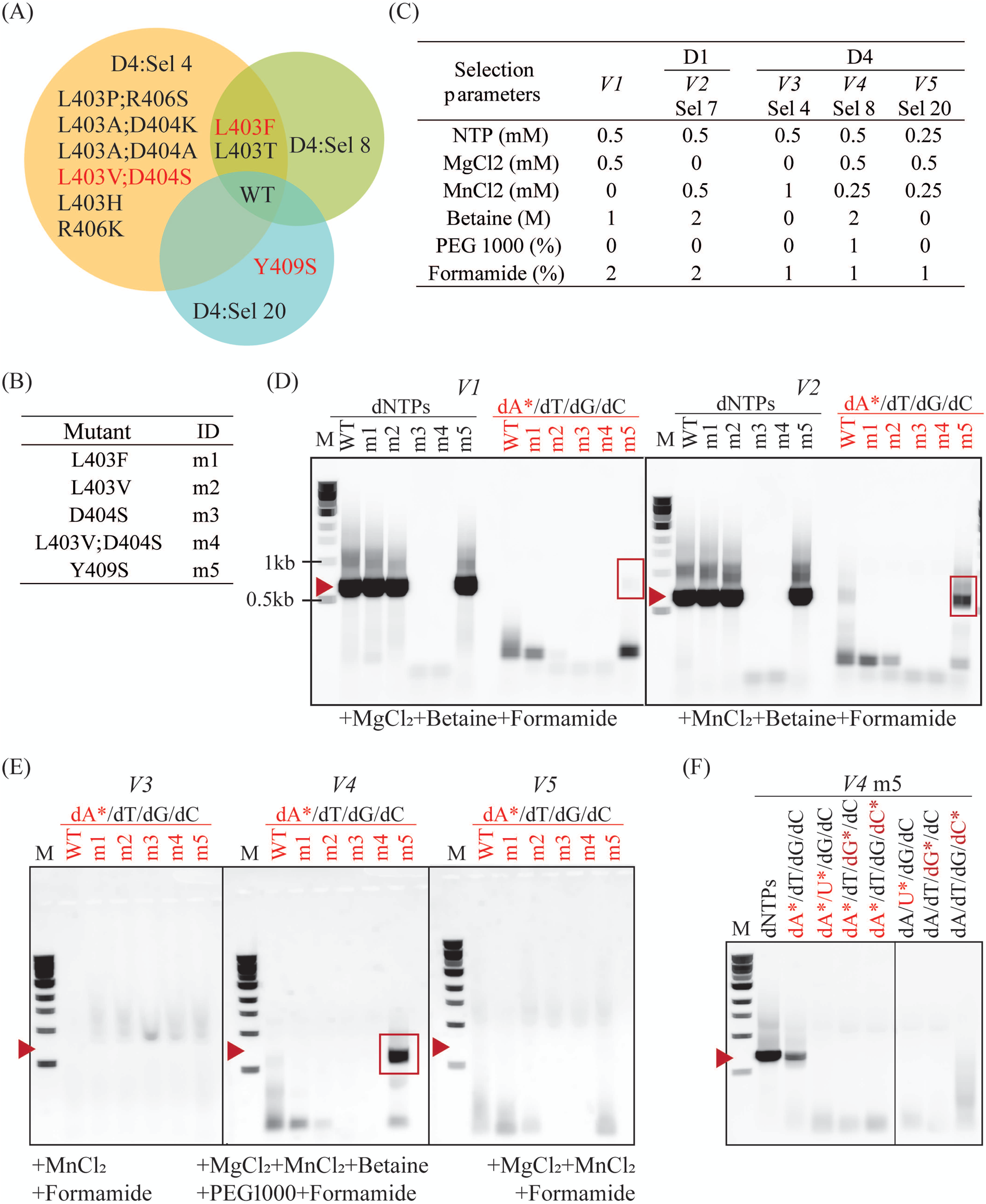
Incorporation of 2’f-dA in PCR by enriched KOD DNAP variants. (A) Venn diagram showing mutations isolated in successful D4 and D1 selections. Mutants labelled in red were selected for further characterization. (B) Mutants selected and their corresponding ID in experiments (C-F). (C) PCR conditions mimicking selection parameters from D1 Sel 7 and D4 Sel 4, 8 and 20, labelled from *V2 – V5*. *V1* corresponds to selection parameters that enabled the isolation of a mesophilic HNA synthetase (Handal-Marquez et al., 2022). (D) PCR products from each mutant in *V1* and *V2* reaction conditions with dNTPs or 2’f-dA substitution (dA*). Red highlights indicate reactions with 2’f-dA, red arrows indicate the expected molecular weight of the PCR product (664 bp), and red rectangles denote correctly sized products (664 bp). (E) PCR products from reactions with *V3*, *V4* and *V5* conditions. (F) PCR products from reactions with *V4* conditions and dNTPs, or substitutions with 2’Fluoro-modified analogues.

We opted for testing their activity in PCR with the amplification of the same 664bp-fragment generated during selection and using some of the parameters from successful selection conditions (Figure 9C). As expected, mutations to D404 (as seen in m3 and m4) abolish DNA synthesis activity and account for the lack of DNA amplification. PCR amplification biases coupled to insufficient sequencing depth are sufficient to explain how m4 could be identified in Sel 4. All other variants successfully amplified DNA in conditions previously used for HNA synthetase selection, V1 (Handal-Marquez et al., 2022), and used in D1 selections (V2). However, only m5 (and WT to a much lower extent) could sustain PCR in the presence of 2’F-dA.

V3, V4 and V5 reaction conditions correspond to those in selections 4, 8, and 20 from Design 4, respectively. No mutant efficiently incorporates 2’F-dA, aside from m5 in V4 (Figure 9D). M5 performed best in V4 (Sel 8), even though it was originally isolated in Sel 20 conditions (V5). Like with D404S, biases and under-sampling can account for the misclassification. In addition, it is possible that the PCR conditions using purified enzymes, differ significantly from the reaction conditions within the emulsion during selection. Comparing all the conditions tested, it is clear that betaine is an essential enhancer for the synthesis of DNA containing 2’F-dA, recapitulating the results obtained in D2. These results confirm the importance of betaine as a selection and PCR enhancer, as previously observed. However, they also highlight that while these conditions may effectively isolate functional variants, fine-tuning reaction parameters for optimal performance by purified variants remains essential.

Lastly, we tested if m5 could incorporate other 2’F-modified substrates in the V4 reaction conditions. As shown in Figure 9E, m5 cannot carry out PCR with 2’F-U, 2’F-dG or 2’F-dC substitutions or in combination with 2’F-dA. It is possible that m5 is only capable of 2’F-dA incorporation and selection with the other analogues may identify alternative mutations that are better suited for their incorporation.

### 3.5 Sequencing coverage analysis in directed evolution experiments

Sequencing coverage requirements are well established for genomic and transcriptomic analysis (Petrackova et al., 2019; Sims et al., 2014) but remain unexplored in directed evolution. Here we define sequencing coverage as the number of sequencing reads divided by the theoretical library size, as traditional definitions are not applicable.

For small libraries like a 2-point saturation mutagenesis library (that has a theoretical size of 400 possible variants), achieving 60x coverage is readily feasible. However, as the number of targeted sites increases, the library size grows significantly, making deep sequencing considerably more expensive. For instance, achieving 60x coverage with the 5-point saturation mutagenesis library would require at least 192 million reads. Even 10x coverage becomes prohibitively expensive when looking to investigate multiple libraries and selection conditions.

Therefore, we decided to use data obtained from sequencing the 2-residue library (selection 7), that reached coverages of 60x, to investigate the impact of coverage on the identification of the expected top 20 enriched variants. Detection of enriching sequences is expected to depend on their frequency post-selection and on the depth of sampling of the population. To test that assumption and to determine which other factors are relevant for the detection of enrichment, we investigated the impact of sampling from the 60x-coverage data (seen as the “truth” set) on the identification of enriched sequences. Sampling from the 60x coverage data allows low coverage to be simulated and we looked at the probability of an enriched sequence being a “true” enriched variant, as well as the probability that enriched variants are detected. For each condition, ten trials were used to obtain a measure of reproducibility in the identification.

**Figure 10.**
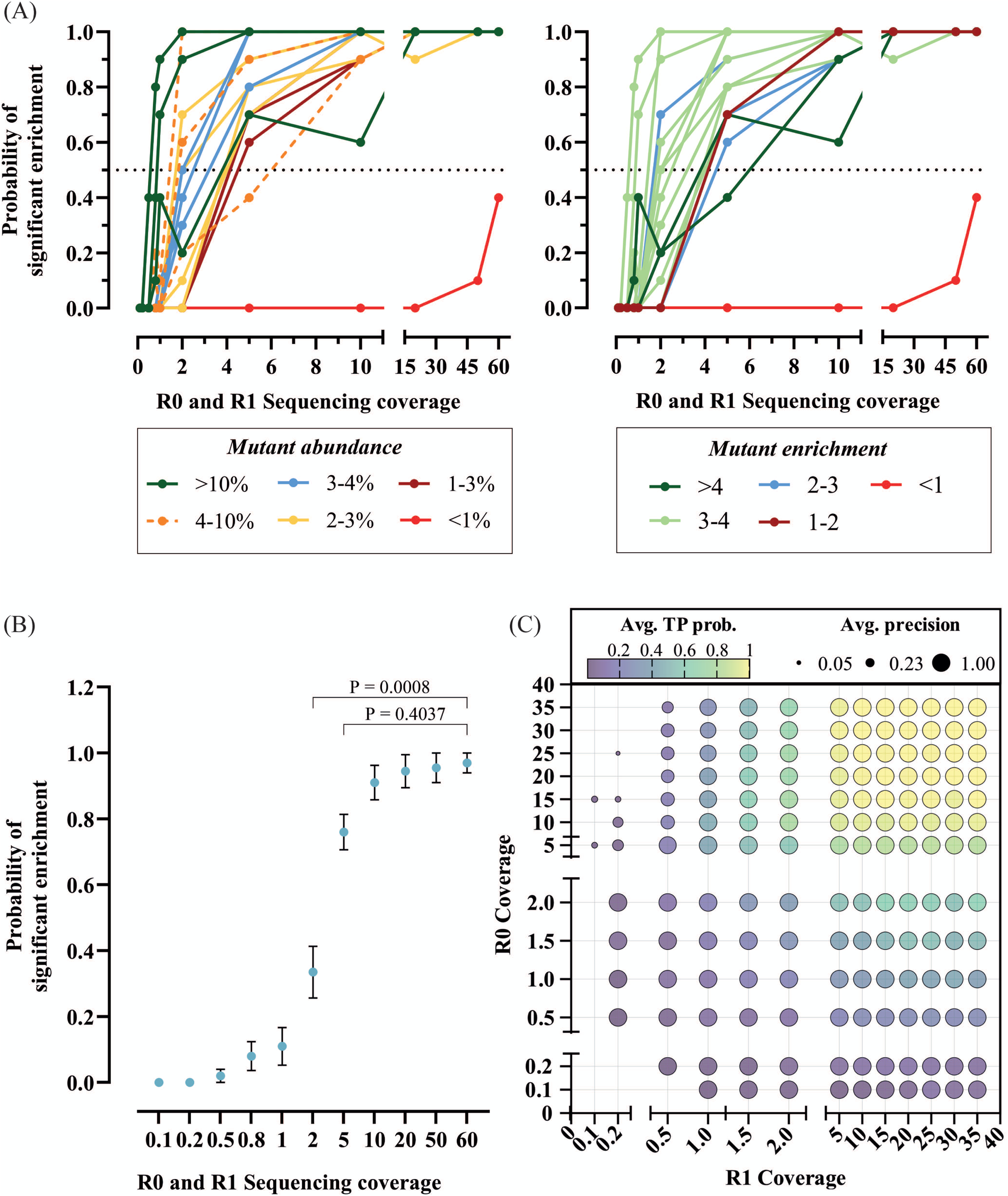
Sequencing coverage analysis in directed evolution. (A) Ten subsets of sequencing reads at varying coverages pre-(R0) and post-(R1) selection from D1 Sel 7 were extracted, and the probability of isolating significantly enriched mutants across these subsets was calculated and plotted. Mutants are color-coded by frequency (left) and enrichment scores (right). (B) Average probability of identifying expected significantly enriched mutants across trials, with significance comparisons conducted against 60x coverage using Friedman test followed by Dunn’s test with Bonferroni correction. Only significant differences were found between 60x and 2x or lower coverages. (C) 2D bubble plot illustrating the impact of sequencing coverages pre-(R0) and post-(R1) selection on true positive (TP) probability and precision, where precision is calculated as TP divided by the sum of TP and false positives (FP). Low probability scores indicate reduced likelihood of identifying all significantly enriched mutants, while precision measures the accuracy of positive predictions.

First, we tested the rate of mutant detection against mutant abundance. As shown in Figure 10A (left), enriched sequences with abundance above 2% post-selection could be accurately identified in 80% of the simulations. Some variation on the detection frequency arises due to low abundance pre-selection, preventing its correct classification despite high abundance post-selection and significant enrichment. Figure 10A (right), shows the rate of identification against mutant enrichment. As expected, high enrichment (>3) scores enable the detection of significantly enriched mutants at lower sequencing coverages. Unexpectedly, igh sequencing coverages can also lead to the recovery of false positives, such as mutant FE (labelled in red).

Overall, as shown in Figure 10B, the probability of significant enrichment only differs between the maximum coverage tested, 60x, and 0.1x, 0.2x, 0.5x, 0.8x, 1x and 2x, suggesting that coverages over 2x do not significantly change the probability of recovering significantly enriched mutants.

As it would be expected, high balanced coverage identifies all enriching mutants correctly (high precision) and robustly (high accuracy). As coverage is reduced, the probability of identifying all enriching mutants drops significantly when coverage falls below 5x (Figure 10C). Nonetheless, the enriched mutants detected in the sparser data remain accurate - i.e. enriching variants picked out in analysis are true but the list may be incomplete. In the 5-residue library where coverage was low (< 0.5%) but balanced (R0 coverage ≈ R1 coverage), the Y409S mutant is one such example. It was picked up on only one of the selections but upon further testing (Figure 9) it was confirmed to have enhanced activity.

When coverage is unbalanced, particularly when R0 coverage is high and R1 coverage is low, the likelihood of identifying enriched variants drops and the recovery of false positives increases. These results further support that while selection can narrow down the list of putative functionally relevant variants, they need to be ultimately biochemically tested.

## 4 Discussion and conclusion

Protein evolution can be visualized as an adaptive walk on a fitness landscape, where fitness values are distributed over the sequence space (genotypes) (Wright, 1932). In this framework, sequences can occupy peaks (high fitness) or valleys (low fitness) on the landscape. Valley crossing can be the rate-limiting step for protein engineering and may require the screening and sampling of a large proportion of the vast sequence space. Tools like directed evolution facilitate the exploration of larger proportions of the sequence space simultaneously and accelerate the discovery of relevant functional variants. While directed evolution have enabled the engineering of polymerase variants with desired properties and substrate specificities, the effects of the selection parameters on the efficiency of selection and on the behavior of mutants has not been previously investigated.

Replicative DNA-dependent DNA polymerases are complex, multifunctional enzymes displaying polymerase and exonuclease activities with complex dynamics (Kropp et al., 2017; Pinheiro, 2019). These functions may vary depending on the identity of their substrates and products and thus variant activity is context-dependent, adding yet another layer of complexity to directed evolution experiments. Additionally, selection is a multi-layered, complex process and while recovery yield can inform if a selection was successful, it does not inform on how well it performed.

We demonstrate here that a DoE-guided approach towards selection optimization is possible and can be used to optimize selection conditions. This approach facilitates selection parameter analysis without requiring empirical characterization of all enriched variants. If no mutants are known to display the target function to any degree, an alternative approach that introduces a negative and tractable control is available. For this, the library can be mixed with 10% of non-functional variants (e.g., catalytic site deletion mutants) that can be easily detectable. This method enables the comparison of observed enrichment against this negative control, facilitating the monitoring of non-functional variant depletion during selection. The reduction in non-functional variants serves as a reliable indicator of selection success, validating the efficacy of the directed evolution process. Methods such as digital PCR (dPCR) could be incorporated into the pipeline to facilitate the quantification of selection products directly from selection, without the additional amplification steps that may introduce PCR biases, and identify, with greater accuracy, differences across experimental runs.

The implementation of the DoE-guided polymerase directed evolution allowed us to determine important selection conditions for optimal recovery as well as their impact on polymerase fidelity. For instance, betaine was detected as an important factor for DNA replication as well as 2’F-dA and 2’F-dT incorporation whereas formamide and manganese appeared to have a negative effect on selection recovery. Additionally, significant recovery was only identified with 2’F-dA and dNTP nucleotide mixes, indicating that the isolation of variants capable of incorporating the remaining 2’F-modified analogues may require alternative selection conditions. These insights would have been considerably more challenging to determine without prior selection benchmarking, highlighting the strengths of our proposed pipeline.

For large sequence spaces, selection optimization is more complex. To bypass this limitation, a fraction of the entire population (a subset of the input library) can be used to benchmark selection parameters and enrichment cut-offs. Alternatively, an initial round of selection with low stringency can be implemented to reduce the population of inactive variants and amplify the signal in subsequent DoE experimental runs.

In addition, our proposed pipeline efficiently explores multiple protein functions and aids in identifying optimal parameters for isolating variants with desired phenotypes. For example, notable differences in mutant enrichment patterns were observed between D1 selections 8 and 2, both utilizing 2’F-dA in the nucleotide mix. Selection 2 exhibited the highest substitution error rates, indicating that its parameters may enrich variants capable of more proficient 2’F-dA incorporation at the cost of fidelity, compared to those from Selection 8 conditions.

Machine learning can be used to infer more complex interactions between enrichment, fidelity, and selection conditions. Our dataset was not sufficiently large for robust analysis by multiple ML algorithms (data not shown) but more advanced imputation methods with neural networks (Choudhury and Pal, 2019) remain possible. Still, correlation analysis between error rates and enrichment suggests these two parameters are not directly related and may depend on other parameters from individual selections.

Lastly, we wanted to investigate the sequencing coverage required for protein engineering through directed evolution. We found that a coverage of 5x was sufficient to isolate 80% of all the significantly enriched mutants with 0% chance of false positives. We also found that with coverages as low as 0.5x, while the probability of obtaining significantly enriched variants is low, the probability of false positives remains close to zero. We also observed that it is important to have equal coverage pre- and post-selection, particularly at low (<5x) coverages to obtain accurate mutant enrichment patterns.

In summary, we carried out an in-depth analysis of selection and developed an approach to investigate different levels of selection efficiency. We demonstrated the importance of selection parameters and their effect on the population-and the individual mutant-level. Optimizing selection parameters is crucial for the discovery of desired variants. We also demonstrated that a simple metric for sequencing coverage should be used to standardize sequencing in directed evolution experiments and that low sequencing coverage is enough to obtain precise and accurate mutant enrichment patterns.

## Supporting information

Supplementary_information, scripts and analyses

## 6 Author Contributions

PHM: conceptualization, formal analysis, methodology, investigation, data curation, writing, editing and funding acquisition. HN: expression polymerase variants and PCR assays with 2’-Fluoro-analogues. VBP: supervision, conceptualization, methodology, editing, funding acquisition.

## Acknowledgments

The authors thank Dr. Ross Kent from Synthace for helping with the generation of the JMP DoE.

## 9 Data Availability Statement

NGS sequencing data generated and analyzed can be found in the NCBI SRA database (BioProject: PRJNA1096033).

