## Supplementary_information, scripts and analyses for "Navigating directed evolution efficiently: optimizing selection conditions and selection output analysis": S1.docx

Supplementary Material

| **Category** | **Name** | **Sequence** |
| --- | --- | --- |
| **Library construction** | KOD_Sat_403-404_F1 | CATAGTGTACNNKNNKTTTAGATCCCTGTACC |
|  | KOD_Sat_403-404_R1 | TTCTCCCACAACCCTCTC |
|  | KOD_5ptSat_408-409_F1 | AGATCCNNKNNKCCCTCAATCATC |
|  | KOD_5ptSat_403-405_R1 | KNNKNNKNNGTACACTATGTTCTCCCAC |
|  | PH_Delta_KOD_F1 | TCCCTGTACCCCTCAATC |
|  | PH_Delta_KOD_R1 | ATCTTCCATCGAGTAGCG |
| **Selection** | CSR_Sel_SHORT_F3 | cgagctgatcatcaacGAAATAACCACAGCCTGG |
|  | CSR_Sel_SHORT_R3 | cagctcatgctatatctCCGTTACGCTCTCTGC |
|  | CSR_Rec_SHORT_F1 | CGAGCTGATCATCAAC |
|  | CSR_Rec_SHORT_R1 | CAGCTCATGCTATATCC |
|  | spCSR_Innest_MUT_F3 | aaaGCTCTTCtGAAGGTCACATAtGAGCTTGGGAAG |
|  | spCSR_Innest_MUT_R3 | aaaGCTCTTCtCCTTGCATAcCCGTAGTAACCGTAG |
| **NGS Amplicon generation** | WP2_D1_Seq_F1_R0 | atgGCTACTCGATGGAAGATG |
|  | WP2_D1_Seq_F1_lib2 | atgGCTACTCGATGGAAGATG |
|  | WP2_D1_Seq_F1_lib5 | tgaGCTACTCGATGGAAGATG |
|  | WP2_D1_Seq_F1_lib7 | gatGCTACTCGATGGAAGATG |
|  | WP2_D1_Seq_F1_lib8 | actGCTACTCGATGGAAGATG |
|  | WP2_D1_Seq_F1_lib11 | cagGCTACTCGATGGAAGATG |
|  | WP2_D1_Seq_R1 | CCTCTAGGAGGTCTCCTAG |
|  | WP2_D4_Seq_F1 | gatGCTACTCGATGGAAGATG |
|  | WP2_D4_Seq_R1 | CCTCTAGGAGGTCTCCTAG |
| **Generation of polymerase variants for PCR** | KOD_D404S_fw | ACATAGTGTACCTATCTTTTAGATCC |
|  | KOD_L403F_fw | ACATAGTGTACTTCGATTTTAGATCC |
|  | KOD_L403V-D404S_fw | ACATAGTGTACGTATCTTTTAGATCC |
|  | KOD_L403V_fw | ACATAGTGTACGTAGATTTTAGATCC |
|  | KOD_mut_rv1 | TCTCCCACAACCCTC |
|  | KOD_Y409S_H5 | AGATCCCTGTCTCCCTCAATCATCATC |
|  | KOD_mut_rv3 | AAAATCTAGGTACACTATG |

**Table S1.** Sequences of the oligonucleotides and templates used in this study. All sequences are written in the 5’🡪3’ direction. N:A/C/G/T; K:G/T

**pET23-KOD-Exo-**

TGGCGAATGGGACGCGCCCTGTAGCGGCGCATTAAGCGCGGCcgcTGTGGTGGTTACGCGCAGCGTGACCGCTACACTTGCCAGCGCCCTAGCGCCCGCTCCTTTCGCTTTCTTCCCTTCCTTTCTCGCCACGTTCGCCGGCTTTCCCCGTCAAGCTCTAAATCGGGGGCTCCCTTTAGGGTTCCGATTTAGTGCTTTACGGCACCTCGACCCCAAAAAACTTGATTAGGGTGATGGTTCACGTAGTGGGCCATCGCCCTGATAGACGGTTTTTCGCCCTTTGACGTTGGAGTCCACGTTCTTTAATAGTGGACTCTTGTTCCAAACTGGAACAACACTCAACCCTATCTCGGTCTATTCTTTTGATTTATAAGGGATTTTGCCGATTTCGGCCTATTGGTTAAAAAATGAGCTGATTTAACAAAAATTTAACGCGAATTTTAACAAAATATTAACGTTTACAATTTCAGGTGGCACTTTTCGGGGAAATGTGCGCGGAACCCCTATTTGTTTATTTTTCTAAATACATTCAAATATGTATCCGCTCATGAGACAATAACCCTGATAAATGCTTCAATAATATTGAAAAAGGAAGAGTATGAGTATTCAACATTTCCGTGTCGCCCTTATTCCCTTTTTTGCGGCATTTTGCCTTCCTGTTTTTGCTCACCCAGAAACGCTGGTGAAAGTAAAAGATGCTGAAGATCAGTTGGGTGCACGAGTGGGTTACATCGAACTGGATCTCAACAGCGGTAAGATCCTTGAGAGTTTTCGCCCCGAAGAACGTTTTCCAATGATGAGCACTTTTAAAGTTCTGCTATGTGGCGCGGTATTATCCCGTATTGACGCCGGGCAAGAGCAACTCGGTCGCCGCATACACTATTCTCAGAATGACTTGGTTGAGTACTCACCAGTCACAGAAAAGCATCTTACGGATGGCATGACAGTAAGAGAATTATGCAGTGCTGCCATAACCATGAGTGATAACACTGCGGCCAACTTACTTCTGACAACGATCGGAGGACCGAAGGAGCTAACCGCTTTTTTGCACAACATGGGGGATCATGTAACTCGCCTTGATCGTTGGGAACCGGAGCTGAATGAAGCCATACCAAACGACGAGCGTGACACCACGATGCCTGCAGCAATGGCAACAACGTTGCGCAAACTATTAACTGGCGAACTACTTACTCTAGCTTCCCGGCAACAATTAATAGACTGGATGGAGGCGGATAAAGTTGCAGGACCACTTCTGCGCTCGGCCCTTCCGGCTGGCTGGTTTATTGCTGATAAATCTGGAGCCGGTGAGCGTGGGTCTCGCGGTATCATTGCAGCACTGGGGCCAGATGGTAAGCCCTCCCGTATCGTAGTTATCTACACGACGGGGAGTCAGGCAACTATGGATGAACGAAATAGACAGATCGCTGAGATAGGTGCCTCACTGATTAAGCATTGGTAACTGTCAGACCAAGTTTACTCATATATACTTTAGATTGATTTAAAACTTCATTTTTAATTTAAAAGGATCTAGGTGAAGATCCTTTTTGATAATCTCATGACCAAAATCCCTTAACGTGAGTTTTCGTTCCACTGAGCGTCAGACCCCGTAGAAAAGATCAAAGGATCTTCTTGAGATCCTTTTTTTCTGCGCGTAATCTGCTGCTTGCAAACAAAAAAACCACCGCTACCAGCGGTGGTTTGTTTGCCGGATCAAGAGCTACCAACTCTTTTTCCGAAGGTAACTGGCTTCAGCAGAGCGCAGATACCAAATACTGTaCTTCTAGTGTAGCCGTAGTTAGGCCACCACTTCAAGAACTCTGTAGCACCGCCTACATACCTCGCTCTGCTAATCCTGTTACCAGTGGCTGCTGCCAGTGGCGATAAGTCGTGTCTTACCGGGTTGGACTCAAGACGATAGTTACCGGATAAGGCGCAGCGGTCGGGCTGAACGGGGGGTTCGTGCACACAGCCCAGCTTGGAGCGAACGACCTACACCGAACTGAGATACCTACAGCGTGAGCTATGAGAAAGCGCCACGCTTCCCGAAGGGAGAAAGGCGGACAGGTATCCGGTAAGCGGCAGGGTCGGAACAGGAGAGCGCACGAGGGAGCTTCCAGGGGGAAACGCCTGGTATCTTTATAGTCCTGTCGGGTTTCGCCACCTCTGACTTGAGCGTCGATTTTTGTGATGCTCGTCAGGGGGGCGGAGCCTATGGAAAAACGCCAGCAACGCGGCCTTTTTACGGTTCCTGGCCTTTTGCTGCGTTATCCCCTGATTCTGTGGCCTTTTGCgCgtCTGCGTTATCCCCTGATTCTGatgttctttcCTGCGTTATCCCCTGATTCTGtggataaccgtattaccgcctttgagtgagCTGCGTTATCCCCTGATTCTGCTGATACCGCTCGCCGCAGCCGAACGACCGAGCGCAGCGAGTCAGTGAGCGAGGAAGCGGAAtAtCGCCTGATGCGGTATTTTCTCCTTACGCATCTGTGCGGTATTTCACACCGCAATGGTGCACTCTCAGTACAATCTGCTCTGATGCCGCATAGTTAAGCCAGTATACACTCCGCTATCGCTACGTGACTGGGTCATGGCTGCGCCCCGACACCCGCCAACACCCGCTGACGCGCCCTGACGGGCTTGTCTGCTCCCGGCATCCGCTTACAGACAAGCTGTGACCGTCTCCGGGAGCTGCATGTGTCAGAGGTTTTCACCGTCATCACCGAAACGCGCGAGGCAGCTGCGGTAAAGCTCATCAGCGTGGTCGTGAAGCGATTCACAGATGTCTGCCTGTTCATCCGCGTCCAGCTCGTTGAGTTTCTCCAGAAGCGTTAATGTCTGGCTTCTGATAAAGCGGGCCATGTTAAGGGCGGTTTTTTCCTGTTTGGTCACTGATGCCTCCGTGTAAGGGGGATTTCTGTTCATGGGGGTAATGATACCGATGAAACGAGAGAGGATGCTCACGATACGGGTTACTGATGATGAACATGCCCGGTTACTGGAACGTTGTGAGGGTAAACAACTGGCGGTATGGATGCGGCGGGACCAGAGAAAAATCACTCAGGGTCAATGCCAGCGCTTCGTTAATACAGATGTAGGTGTTCCACAGGGTAGCCAGCAGCATATGGTGCAGGGCGCTGACTTCCGCGTTTCCAGACTTTACGAAACACGGAAACCGAAGACCATTCATGTTGTTGCTCAGGTCGCAGACGTTTTGCAGCAGCAGTCGCTTCACGTTCGCTCGCGTATCGGTGATTCATTCTGCTAACCAGTAAGGCAACCCCGCCAGCCTAGCCGGGTCCTCAACGACAGGAGCACGATCATGCGCACCCGTGGCCAGGACCCAACGCTGCCCGAGATCTCGATCCCGCGAAATTAATACGACTCACTATAGGGAGACCACAACGGTTTCCCTCTAGAAATAATTTTGTTTAACTTTAAGAAGGAGATATACCATGGATCCTCTAGAGTCGACCTGCAGGCATGCAAGCTTGCGGCcacacAGGAGATAGTCATACATGAAACACAAAGAGGAGAAATTAACTATGAGAGGATCTCACCATCACCATCACCATACGGATCCAAGCGGCCTGGTGCCGCGCGGCAGCATGATCCTCGACACTGACTACATAACCGAGGATGGAAAGCCTGTCATAAGAATTTTCAAGAAGGAAAACGGCGAGTTTAAGATTGAGTACGACCGGACTTTTGAACCCTACTTCTACGCCCTCCTGAAGGACGATTCTGCCATTGAGGAAGTCAAGAAGATAACCGCCGAGAGGCACGGGACGGTTGTAACGGTTAAGCGGGTTGAAAAGGTTCAGAAGAAGTTCCTaGGGAGACCAGTTGAGGTCTGGAAACTCTACTTTACTCATCCGCAGGACGaaCCAGCGATAAGGGACAAGATACGAGAGCATCCAGCAGTTATTGACATCTACGAGTACGACATACCCTTCGCCAAGCGCTACCTCATAGACAAGGGATTAGTGCCAATGGAAGGCGACGAGGAGCTGAAAATGCTCGCCTTCGCGATTGCGACTCTCTACCATGAGGGCGAGGAGTTCGCCGAGGGGCCAATCCTTATGATAAGCTACGCCGACGAGGAAGGGGCCAGGGTGATAACTTGGAAGAACGTGGATCTCCCCTACGTTGACGTCGTCTCGACGGAGAGGGAGATGATAAAGCGCTTCCTCCGTGTTGTGAAGGAGAAAGACCCGGACGTTCTCATAACCTACAACGGCGACAACTTCGACTTCGCCTATCTGAAAAAGCGCTGTGAAAAGCTCGGAATAAACTTCGCCCTCGGAAGGGATGGAAGCGAGCCGAAGATTCAGAGGATGGGCGACAGGTTTGCCGTCGAAGTGAAGGGACGGATACACTTCGATCTCTATCCTGTGATAAGACGGACGATAAACCTGCCCACATACACGCTTGAGGCCGTTTATGAAGCCGTCTTCGGTCAGCCGAAGGAGAAGGTTTACGCTGAGGAAATAACCACAGCCTGGGAAACCGGCGAGAACCTTGAGAGAGTCGCCCGCTACTCGATGGAAGATGCGAAGGTCACATACGAGCTTGGGAAGGAGTTCCTTCCGATGGAGGCCCAGCTTTCTCGCTTAATCGGCCAGTCCCTCTGGGACGTCTCCCGCTCCAGCACTGGCAACCTCGTTGAGTGGTTCCTCCTCAGGAAGGCCTATGAGAGGAATGAGCTGGCCCCGAACAAGCCCGATGAAAAGGAGCTGGCCAGAAGACGGCAGAGCTATGAAGGAGGCTATGTAAAAGAGCCCGAGAGAGGGTTGTGGGAGAACATAGTGTACCTAGATTTTAGATCCCTGTACCCCTCAATCATCATCACCCACAACGTCTCGCCGGATACGCTCAACAGAGAAGGATGCAAGGAATATGACGTTGCCCCACAGGTCGGCCACCGCTTCTGCAAGGACTTCCCAGGATTTATCCCGAGCCTGCTaGGAGACCTCCTAGAGGAGAGGCAGAAGATAAAGAAGAAGATGAAGGCCACGATTGACCCGATCGAGAGGAAGCTCCTCGATTACAGGCAGAGGGCAATCAAGATCCTGGCAAACAGCTACTACGGTTACTACGGCTATGCAAGGGCGCGCTGGTACTGCAAGGAGTGTGCAGAGAGCGTAACGGCCTGGGGAAGGGAGTACATAACGATGACCATCAAGGAGATAGAGGAAAAGTACGGCTTTAAGGTAATCTACAGCGACACCGACGGATTTTTTGCCACAATACCTGGAGCCGATGCTGAAACCGTCAAAAAGAAGGCTATGGAGTTCCTCAAGTATATCAACGCCAAACTTCCGGGCGCGCTTGAGCTCGAGTACGAGGGCTTCTACAAACGCGGCTTCTTCGTCACGAAGAAGAAGTATGCGGTGATAGACGAGGAAGGCAAGATAACAACGCGCGGACTTGAGATTGTGAGGCGTGACTGGAGCGAGATAGCGAAAGAGACGCAGGCGAGGGTTCTTGAAGCTTTGCTAAAGGACGGTGACGTCGAGAAGGCCGTGAGGATAGTCAAAGAAGTTACCGAAAAGCTGAGCAAGTACGAGGTTCCGCCGGAGAAGCTGGTGATCCACGAGCAGATAACGAGGGATTTAAAGGACTACAAGGCAACCGGTCCCCACGTTGCCGTTGCCAAGAGGTTGGCCGCGAGAGGAGTCAAAATACGCCCTGGAACGGTGATAAGCTACATCGTGCTCAAGGGCTCTGGGAGGATAGGCGACAGGGCGATACCGTTCGACGAGTTCGACCCGACGAAGCACAAGTACGACGCCGAGTACTACATTGAGAACCAGGTTCTCCCAGCCGTTGAGAGAATTCTGAGAGCCTTCGGTTACCGCAAGGAAGACCTGCGCTACCAGAAGACGAGACAGGTTGGTTTGAGTGCTTGGCTGAAGCCGAAGGGAACTTGATCGATGCTCCGAGATGAGGTAGGATGGCTGGCTTACGGTGTTACTGCTGAGGAATGAgccatcCTCGAGCACCACCACCACCACCACTGAGATCCGGCTGCTAACAAAGCCCGAAAGGAAGCTGAGTTGGCTGCTGCCACCGCTGAGCAATAACTAGCATAACCCCTTGGGGCCTCTAAACGGGTCTTGAGGGGTTTTTTGCTGAAAGGAGGAACTATATCCGGAT

**Table S2.** Sequences of all the plasmids used in this study.

| **Pipeline step** | **Sequences output R0** | **Sequences output R1** | | | | |
| --- | --- | --- | --- | --- | --- | --- |
|  |  | **Sel 2** | **Sel 5** | **Sel 7** | **Sel 8** | **Sel 11** |
| **Total reads** | 466,836 | 462,439 | | | | |
| **Total paired reads** | 433794 (93%) | 374,592 (81%) | | | | |
| **Quality filtering** | 417398 (89%) | 355,811 (77%) | | | | |
| **Filtering by 3’ and 5’ sequence and translating** | 211,965* (45%) | 55,228* (12%) | 29,657* (6%) | 28,297* (6%) | 42,943* (9%) | 64,898* (14%) |
| **Unique sequences** | 450 | 338 | 310 | 295 | 314 | 335 |
| **Coverage** | 530x | 138x | 74x | 71x | 107x | 162x |

**Table S3.** Analysis by next generation sequencing of the Design 1 recovered **sequences.** Total read number obtained and the impact of the analysis pipeline are shown. *Number of sequences used in downstream analysis.

| **Pipeline step** | **Sequences output R0** | **Sequences output R1** | | |
| --- | --- | --- | --- | --- |
|  |  | **Sel 4** | **Sel 8** | **Sel 20** |
| **Total reads** | 2,508,732 | 753,984 | 911,540 | 833,199 |
| **Total paired reads** | 2,128,712 (85%) | 695,596 (92%) | 830,478 (91%) | 790,840 (95%) |
| **Quality filtering** | 2,030,557 (81%) | 667,972 (89%) | 790,840 (87%) | 689,774 (83%) |
| **Filtering by 3’ and 5’ sequence and translating** | 1,400,528 (56%) | 577,132 (77%) | 668,635 (73%) | 566,024 (68%) |
| **Unique sequences** | 531,097 | 1,428 | 1,058 | 16,300 |
| **Coverage** | 0.5x | 0.2x | 0.2x | 0.2x |

**Table S4.** Analysis by next generation sequencing of the Design 4 recovered sequences. Total read number obtained and the impact of the analysis pipeline are shown. *Number of sequences used in downstream analysis.
