## Supplementary_information, scripts and analyses for "Navigating directed evolution efficiently: optimizing selection conditions and selection output analysis": S2_DOE_Factor_Analysis.R

```
# Load packages
library(glmnet)
library("readxl")
library(dplyr)
library(tidyr)

# Import datasets

#D1
D1_Rec = read_excel("C:\\Users\\PH\\OneDrive - KU Leuven\\PhD\\Polymerase_Landscape_Mapping\\05_Writing\\01_Manuscripts\\Selection_landscape_mapping\\SI\\D1_Rec.xlsx")
D1_Rec = data.frame(D1_Rec)
head(D1_Rec) #sel recovery
D1_Rec$NTP_Type = factor(D1_Rec$NTP_Type, levels = c('DNA', '2FATP'), labels = c(1, 2))

#D2
D2_Rec = read_excel("C:\\Users\\PH\\OneDrive - KU Leuven\\PhD\\Polymerase_Landscape_Mapping\\05_Writing\\01_Manuscripts\\Selection_landscape_mapping\\SI\\D2_Rec.xlsx")
D2_Rec = data.frame(D2_Rec)
head(D2_Rec)
D2_Rec$NTP_Type = factor(D2_Rec$NTP_Type)

#D4
D4_Rec = read_excel("C:\\Users\\PH\\OneDrive - KU Leuven\\PhD\\Polymerase_Landscape_Mapping\\05_Writing\\01_Manuscripts\\Selection_landscape_mapping\\SI\\D4_Rec.xlsx")
D4_Rec = data.frame(D4_Rec)
D4_Rec = D4_Rec[, c("Betaine","MgCl2","MnCl2","PEG1000","NTP_Conc","Block","Y2_Innest30_Norm","Y2_Innest30_Gel_Norm")]
head(D4_Rec)

#D1 Mutant Fidelity of Positive selections
fidelity_positive = read.csv("C:\\Users\\PH\\OneDrive - KU Leuven\\PhD\\Polymerase_Landscape_Mapping\\05_Writing\\01_Manuscripts\\Selection_landscape_mapping\\SI\\individual_files\\NGS_Analysis\\D1_Fidelity_Matched.csv")
fidelity_positive = data.frame(fidelity_positive)
fidelity_positive$ID <- ifelse(is.na(fidelity_positive$ID), "NA", fidelity_positive$ID)
fidelity_positive$ID <- as.factor(fidelity_positive$ID)
# reshaping
fidelity_positive <- fidelity_positive %>%
  pivot_longer(cols = -ID,
               names_to = c(".value", "suffix"),
               names_sep = "_lib") %>%
  rename(
    deletions = deletions,
    insertions = insertions,
    substitutions = substitutions,
    Sel = Sel
  )

D1_sel_parameters <- D1_Rec %>%
  mutate(Sel = row_number())
fidelity_positive <- merge(fidelity_positive, D1_sel_parameters, by = "Sel", all.x = TRUE)
head(fidelity_positive)
fidelity_positive_sels <- subset(fidelity_positive, Sel != 0)
head(fidelity_positive_sels)


#D1 Mutant Fidelity Positive and Engineered Negative selections
fidelity_complete_df = read.csv("C:\\Users\\PH\\OneDrive - KU Leuven\\PhD\\Polymerase_Landscape_Mapping\\05_Writing\\01_Manuscripts\\Selection_landscape_mapping\\SI\\individual_files\\NGS_Analysis\\D1_Mutant_Fidelity_FailedSelections_Engineered_1.csv")
fidelity_complete_df = data.frame(fidelity_complete_df)
fidelity_complete_df$ID <- ifelse(is.na(fidelity_complete_df$ID), "NA", fidelity_complete_df$ID)
fidelity_complete_df$ID <- as.factor(fidelity_complete_df$ID)
columns_to_remove <- grep("^(mean|std|lower_quantile|upper_quantile|range)", colnames(fidelity_complete_df))
fidelity_complete_df <- fidelity_complete_df[, -columns_to_remove]

# reshaping
fidelity_complete <- fidelity_complete_df
fidelity_complete <- fidelity_complete %>%
  pivot_longer(cols = -ID,
               names_to = c(".value", "suffix"),
               names_sep = "_lib") %>%
  rename(
    deletions = deletions,
    insertions = insertions,
    substitutions = substitutions,
  )
names(fidelity_complete)[names(fidelity_complete) == "suffix"] <- "Sel"
fidelity_complete <- merge(fidelity_complete, D1_sel_parameters, by = "Sel", all.x = TRUE)
head(fidelity_complete)
fidelity_complete_sels <- subset(fidelity_complete, Sel != 0)
head(fidelity_complete_sels)

# Sub-setting data to include only Enriched mutants
fidelity_enriched = read.csv("C:\\Users\\PH\\OneDrive - KU Leuven\\PhD\\Polymerase_Landscape_Mapping\\05_Writing\\01_Manuscripts\\Selection_landscape_mapping\\SI\\individual_files\\NGS_Analysis\\D1_Fidelity_Enriched.csv")
enriched_muts <- as.character(trimws(fidelity_enriched$ID))

enriched_pos <- fidelity_positive_sels[as.character(fidelity_positive_sels$ID) %in% enriched_muts, ]
head(enriched_pos)

enriched_all <- fidelity_complete_sels[fidelity_complete_sels$ID %in% enriched_muts, ]
head(enriched_all)

# To file for machine learning modelling
write.csv(fidelity_positive_sels, file = "fidelity_positive_sels.csv", row.names = FALSE)
write.csv(fidelity_complete_sels, file = "fidelity_complete_sels.csv", row.names = FALSE)
write.csv(enriched_pos, file = "enriched_pos_sels.csv", row.names = FALSE)
write.csv(enriched_all, file = "enriched_complete_sels.csv", row.names = FALSE)

############################################################################################################

# Correlation across response variables
library(ggcorrplot)
# Compute correlation analysis across response variables
cor_matrix_D1 <- cor(D1_Rec[10:14], method = "kendall")
ggcorrplot(cor_matrix_D1, method = "circle")

cor_matrix_D2 <- cor(D2_Rec[14:15], method = "kendall")
ggcorrplot(cor_matrix_D2, method = "circle")

cor_matrix_fidelity <- cor(fidelity_positive[4:7], method = "kendall")
ggcorrplot(cor_matrix_fidelity, method = "circle")
write.csv(cor_matrix_fidelity, file = "fidelity_positive_sels_cor.csv", row.names = TRUE)

cor_matrix_fidelity_enriched <- cor(enriched_pos[4:7], method = "kendall")
ggcorrplot(cor_matrix_fidelity_enriched, method = "circle")
write.csv(cor_matrix_fidelity_enriched, file = "fidelity_positive_enriched_cor.csv", row.names = TRUE)

#############################################################################################################

# Boruta factor selection
library(Boruta)
library(randomForest)

# Predictors and response variables
predictors_D1 <- D1_Rec[, c("BSA","Betaine","MgCl2","MnCl2","PEG1000","Formamide","NTP_Conc","NTP_Type","Time")]
response_variables_D1 <- c("Y1_Rec20", "Y1_Rec28", "Y2_Innest20", "Y2_Innest28", "Y2_Innest28_S")

predictors_D2 <- D2_Rec[, c("Betaine","MnCl2","Formamide","NTP_Type")]
response_variables_D2 <- c("Y1_Rec28_Norm","Y2_Innest28_S")

predictors_D4 <- D4_Rec[, c("Betaine","MgCl2","MnCl2","PEG1000","NTP_Conc","Block")]
response_variables_D4 <- c("Y2_Innest30_Norm", "Y2_Innest30_Gel_Norm")

predictors_fidelity <- fidelity_positive_sels[, c("ID","bases_sequenced","BSA","Betaine","MgCl2","MnCl2","PEG1000","Formamide","NTP_Conc","NTP_Type","Time")]
predictors_pos_enriched <- enriched_pos[, c("ID","bases_sequenced","BSA","Betaine","MgCl2","MnCl2","PEG1000","Formamide","NTP_Conc","NTP_Type","Time")]
predictors_fidelity_complete <- fidelity_complete_sels[, c("ID","BSA","Betaine","MgCl2","MnCl2","PEG1000","Formamide","NTP_Conc","NTP_Type","Time")]
predictors_all_enriched <- enriched_all[, c("ID","BSA","Betaine","MgCl2","MnCl2","PEG1000","Formamide","NTP_Conc","NTP_Type","Time")]

response_variables_fidelity <- c("deletions", "insertions","substitutions","Y2_Innest28","Y2_Innest28_S")


run_boruta_analysis <- function(data, predictors, response_variables, maxRuns = 100) {
  # Load required package
  require(Boruta)

  # Initialize an empty list to store selected features for each response variable
  selected_features_list <- list()

  # Loop through each response variable
  for (response_var in response_variables) {
    # Convert the response variable to numeric if it is not already
    if (!is.numeric(data[[response_var]])) {
      data[[response_var]] <- as.numeric(data[[response_var]])
    }

    # Run the Boruta analysis for the current response variable
    boruta_model <- Boruta(x = predictors, y = data[[response_var]], maxRuns = maxRuns)

    # Extract confirmed relevant variables
    selected_features <- getSelectedAttributes(boruta_model, withTentative = FALSE)
    selected_features <- paste_with_none(selected_features)

    # Store selected features for the current response variable
    selected_features_list[[response_var]] <- selected_features
  }

  # Combine all selected features for each response variable
  all_selected_features <- data.frame(Response = response_variables,
                                      features = unlist(selected_features_list),
                                      stringsAsFactors = FALSE)

  return(all_selected_features)
}

# Helper function to concatenate selected features
paste_with_none <- function(lst, collapse = ", ") {
  if (length(lst) > 0) {
    return(paste(lst, collapse = collapse))
  } else {
    return("none")
  }
}


# Boruta Factor Selection - D1
data_D1 <- D1_Rec
selected_features_D1 <- run_boruta_analysis(data_D1, predictors_D1, response_variables_D1)
print(selected_features_D1)

# Boruta Factor Selection - D2
data_D2 <- D2_Rec
selected_features_D2 <- run_boruta_analysis(data_D2, predictors_D2, response_variables_D2)
print(selected_features_D2)

# Boruta Factor Selection - D4
data_D4 <- D4_Rec
selected_features_D4 <- run_boruta_analysis(data_D4, predictors_D4, response_variables_D4)
print(selected_features_D4)

# Boruta Factor Selection - D1 Mutant Fidelity
data_fidelity <- fidelity_positive_sels
fidelity_features <- run_boruta_analysis(data_fidelity, predictors_fidelity, response_variables_fidelity)
print(fidelity_features)

data_enrich_pos <- enriched_pos
fidelity_features <- run_boruta_analysis(data_enrich_pos, predictors_pos_enriched, response_variables_fidelity)
print(fidelity_features)

data_fidelity_complete <- fidelity_complete_sels
fidelity_complete_features <- run_boruta_analysis(data_fidelity_complete, predictors_fidelity_complete, response_variables_fidelity)
print(fidelity_complete_features)

data_enriched_complete <- enriched_all
fidelity_complete_features <- run_boruta_analysis(data_enriched_complete, predictors_all_enriched, response_variables_fidelity)
print(fidelity_complete_features)

#############################################################################################################

# Lasso regression
# Load packages
library(glmnet)
library("readxl")
library(dplyr)
library(caret)

# Function to perform Lasso regression

# Define the run_lasso_regression function
run_lasso_regression <- function(data, predictor_variables, response_variable) {
  # Convert the expanded data to matrix format
  x_matrix <- model.matrix(~., data = data[, c(predictor_variables)])
  # Convert response variable to numeric
  y_vector <- as.numeric(data[[response_variable]])

  # Fit the Lasso regression model with cross-validation
  lasso_model_cv <- cv.glmnet(x_matrix, y_vector, family = "gaussian", alpha = 1)
  # Access the coefficients from the optimal lasso_model_cv
  lasso_coefficients <- coef(lasso_model_cv, s = "lambda.min")
  # Extract the non-zero coefficients (important features)
  important_features <- which(lasso_coefficients[-1] != 0)

  # If only one important feature is selected, add a dummy feature
  if (length(important_features) == 1) {
    x_matrix <- cbind(x_matrix, dummy = 0)
    important_features <- c(important_features, ncol(x_matrix))  # Add the dummy column index
  }
  # Fit the final Lasso model with the optimal lambda value
  lasso_model_final <- glmnet(x_matrix[, important_features, drop = FALSE], y_vector,
                              family = "gaussian", alpha = 1, lambda = lasso_model_cv$lambda.min)

  # Extract the coefficients of the final Lasso model
  final_lasso_coefficients <- coef(lasso_model_final)
  # Set a threshold to exclude coefficients close to zero
  threshold <- 1e-9
  final_important_features <- which(abs(final_lasso_coefficients[-1]) > threshold)

  # Filter the coefficients and feature names using the updated selection
  final_important_coefficients <- final_lasso_coefficients[-1][final_important_features]
  final_lasso_feature_names <- rownames(final_lasso_coefficients)[-1] # Exclude the intercept (first row)
  final_lasso_feature_names <- final_lasso_feature_names[final_important_features]

  # Predict the probabilities using the selected lambda and final important features
  lasso_probs <- predict(lasso_model_final, newx = x_matrix[, important_features, drop = FALSE], type = "response")

  # Calculate the residuals
  lasso_residuals <- y_vector - lasso_probs
  # Calculate Mean Squared Error (MSE)
  mse <- mean(lasso_residuals^2)
  # Calculate Root Mean Squared Error (RMSE)
  rmse <- sqrt(mse)
  # Calculate Mean Absolute Error (MAE)
  mae <- mean(abs(lasso_residuals))
  # Calculate R-squared for the current model
  r_squared <- 1 - sum(lasso_residuals^2) / sum((y_vector - mean(y_vector))^2)

  # Calculate degrees of freedom
  df <- sum(abs(final_lasso_coefficients[-1]) > threshold)

  # Calculate the Residual Sum of Squares (RSS)
  rss <- sum(lasso_residuals^2)

  # Calculate the number of data points (n)
  n <- nrow(x_matrix)

  # Calculate BIC and AIC
  bic <- n * log(rss / n) + df * log(n)
  aic <- n * log(rss / n) + 2 * df

  # Create a dataframe with the performance metrics, coefficients, BIC, and AIC
  performance_metrics <- data.frame(
    Metric = c(final_lasso_feature_names, "r_squared", "mse", "rmse", "mae", "BIC", "AIC"),
    Score = c(final_important_coefficients, r_squared, mse, rmse, mae, bic, aic),
    stringsAsFactors = FALSE
  )

  performance_metrics <- performance_metrics %>%
    pivot_wider(names_from = Metric, values_from = Score)

  # Diagnostic plots for Lasso regression
  # 1. Residuals vs. Fitted Values (independence)
  plot(lasso_probs, lasso_residuals, main = "Residuals vs. Fitted Probabilities",
       xlab = "Fitted Probabilities", ylab = "Residuals")

  # 2. Histogram of Residuals (normality)
  hist(lasso_residuals, main = "Histogram of Residuals", xlab = "Residuals")

  # 3. Q-Q Plot of Residuals (normality)
  qqnorm(lasso_residuals, main = "Q-Q Plot of Residuals")
  qqline(lasso_residuals)

  # Return a list with the necessary outputs, including additional performance metrics and important features
  return(performance_metrics)
}


# stability selection / consensus Lasso

# Load necessary libraries
library(glmnet)
library(tidyverse)
library(reshape2)
library(tidyr)

# Define the run_lasso_regression_stability_selection function

run_lasso_regression_stability_selection <- function(data, predictor_variables, response_variable, num_models = 100, threshold = 0.8) {
  all_outputs <- list()

  for (i in 1:num_models) {
    lasso_model_output <- tryCatch(run_lasso_regression(data, predictor_variables, response_variable),
                                   error = function(e) list(final_lasso_dataframe = NULL,
                                                            performance_metrics = NULL))  # Capture any error and return NULL

    all_outputs[[i]] <- lasso_model_output
  }

  # Remove NULL entries from the list
  all_outputs <- all_outputs[sapply(all_outputs, Negate(is.null))]

  # Extract performance metrics and concatenate into a single data frame
  all_metrics <- bind_rows(all_outputs, .id = "Model")

  # Check if the number of rows in all_metrics is less than num_models
  if (nrow(all_metrics) < num_models) {
    missing_rows <- num_models - nrow(all_metrics)
    missing_data <- matrix(0, ncol = ncol(all_metrics), nrow = missing_rows)
    colnames(missing_data) <- colnames(all_metrics)
    all_metrics <- rbind(all_metrics, as.data.frame(missing_data))
  }

  # Calculate average values for metrics
  #average_metrics <- all_metrics %>%
    #summarize(across(c(names(all_metrics)), mean, na.rm = TRUE))

  return(all_outputs = all_metrics)
}


#### Lasso regression - D1

# Select the columns for z-score normalization (standardization)
predictors_zc_D1 <- c("BSA","Betaine","MgCl2","MnCl2","PEG1000","Formamide","NTP_Conc","Time")
# Standardize the selected numerical features
data_zs_D1 <- data_D1 %>%
  mutate(across(all_of(predictors_zc_D1), scale)) %>%
  mutate(across(c(Y2_Innest28, Y2_Innest28_S), scale)) %>%
  select(all_of(predictors_zc_D1), NTP_Type, Y2_Innest28, Y2_Innest28_S)
predictors_zs_D1 <- names(data_zs_D1)[1:(length(names(data_zs_D1)) - 2)]

#run lasso
lasso_D1_innest28 <- run_lasso_regression_stability_selection(data_zs_D1, predictors_zs_D1, response_variable = "Y2_Innest28")
lasso_D1_innest28_S <- run_lasso_regression_stability_selection(data_zs_D1, predictors_zs_D1, response_variable = "Y2_Innest28_S")

# Export important features as CSV
write.csv(lasso_D1_innest28, file = "SI/individual_files/D1_lasso_innest28G.csv", row.names = FALSE)
write.csv(lasso_D1_innest28_S, file = "SI/individual_files/D1_lasso_innest28S.csv", row.names = FALSE)


#### Lasso regression - D2

# Create dummy variables using one-hot encoding
dummy_data <- dummyVars("~ NTP_Type", data = D2_Rec) %>%
  predict(newdata = D2_Rec)

# Combine one-hot encoded data with original numeric columns
data_zs_D2 <- cbind(D2_Rec[, c("Betaine","MnCl2","Formamide")], dummy_data, D2_Rec[, c("Y1_Rec28_Norm","Y2_Innest28_S")])

# Select the columns for z-score normalization (standardization)
norm_zc_D2 <- c("Betaine","MnCl2","Formamide","Y1_Rec28_Norm","Y2_Innest28_S")
# Standardize the selected numerical features
data_zs_D2 <- data_zs_D2 %>%
  mutate(across(all_of(norm_zc_D2), scale)) %>%
  select("Betaine","MnCl2","Formamide", starts_with("NTP_Type."),"Y1_Rec28_Norm","Y2_Innest28_S")

predictors_zs_D2 <- names(data_zs_D2)[1:(length(names(data_zs_D2)) - 2)]

lasso_D2_innest28 <- run_lasso_regression_stability_selection(data_zs_D2, predictors_zs_D2, response_variable = "Y1_Rec28_Norm")
lasso_D2_innest28_S <- run_lasso_regression_stability_selection(data_zs_D2, predictors_zs_D2, response_variable = "Y2_Innest28_S")

# Export important features as CSV
write.csv(lasso_D2_innest28, file = "SI/individual_files/D2_lasso_innest28_1.csv", row.names = FALSE)
write.csv(lasso_D2_innest28_S, file = "SI/individual_files/D2_lasso_innest28S_1.csv", row.names = FALSE)


#### Lasso regression - D2: quadratic effects and interactions

expand_predictors <- function(data) {
  data <- data %>%
    mutate(
      F1_sq = F1^2,
      F2_sq = F2^2,
      F3_sq = F3^2,
      F1_F2 = F1 * F2,
      F1_F3 = F1 * F3,
      F1_F4 = F1 * F4,
      F1_F5 = F1 * F5,
      F1_F6 = F1 * F6,
      F1_F7 = F1 * F7,
      F1_F8 = F1 * F8,
      F2_F3 = F2 * F3,
      F2_F4 = F2 * F4,
      F2_F5 = F2 * F5,
      F2_F6 = F2 * F6,
      F2_F7 = F2 * F7,
      F2_F8 = F2 * F8,
      F3_F4 = F3 * F4,
      F3_F5 = F3 * F5,
      F3_F6 = F3 * F6,
      F3_F7 = F3 * F7,
      F3_F8 = F3 * F8
    )

  return(data)
}

predictor_vars <- c(
  "F1", "F2", "F3",
  "F4", "F5", "F6", "F7", "F8",
  "F1_sq", "F2_sq", "F3_sq",
  "F1_F4", "F1_F5", "F1_F6", "F1_F7", "F1_F8",
  "F2_F4", "F2_F5", "F2_F6", "F2_F7", "F2_F8",
  "F3_F4", "F3_F5", "F3_F6", "F3_F7", "F3_F8"
)

# Change the column names
# Define the new column names as a named vector

data_zs_D2_v2 <- data_zs_D2
names(data_zs_D2_v2)[names(data_zs_D2_v2) %in% predictors_zs_D2] <- c("F1", "F2", "F3", "F4", "F5", "F6", "F7", "F8")
expanded_data <- expand_predictors(data_zs_D2_v2)

lasso_D2_innest28_2_coded <- run_lasso_regression_stability_selection(expanded_data, predictor_vars, response_variable = "Y1_Rec28_Norm")
lasso_D2_innest28_S_2_coded <- run_lasso_regression_stability_selection(expanded_data, predictor_vars, response_variable = "Y2_Innest28_S")

# mapping feature names
lasso_D2_innest28_2 <- lasso_D2_innest28_2_coded
lasso_D2_innest28_S_2 <- lasso_D2_innest28_S_2_coded

coded_terms <- c("F1", "F2", "F3", "F4", "F5", "F6", "F7", "F8")
original_names <- c("Betaine","MnCl2","Formamide","NTP.Type.A","NTP.Type.AC","NTP.Type.AT","NTP.Type.dNTPs","NTP.Type.T")
names_to_rename1 <- names(lasso_D2_innest28_2_coded)
names_to_rename2 <- names(lasso_D2_innest28_S_2_coded)


map_names <- function(names_to_rename, coded_terms, original_names) {
  mapped_names <- character(length(names_to_rename))
  for (i in seq_along(names_to_rename)) {
    col <- names_to_rename[i]
    if (col == "Model") {
      mapped_names[i] <- "Model"
    } else if (startsWith(col, "F")) {
      parts <- strsplit(col, "_")[[1]]
      new_parts <- character(length(parts))
      for (j in seq_along(parts)) {
        part <- parts[j]
        if (startsWith(part, "F")) {
          index <- as.integer(substring(part, 2))
          if (index <= length(original_names)) {
            new_parts[j] <- original_names[index]
          } else {
            new_parts[j] <- part
          }
        } else {
          new_parts[j] <- part
        }
      }
      mapped_names[i] <- paste(new_parts, collapse = "_")
    } else {
      mapped_names[i] <- col
    }
  }
  return(mapped_names)
}

new_names1 <- map_names(names_to_rename1, coded_terms, original_names)
new_names2 <- map_names(names_to_rename2, coded_terms, original_names)


# mapped feature names
colnames(lasso_D2_innest28_2) <- new_names1
colnames(lasso_D2_innest28_S_2) <- new_names2

# Export important features as CSV
write.csv(lasso_D2_innest28_2, file = "SI/individual_files/D2_lasso_innest28_2.csv", row.names = FALSE)
write.csv(lasso_D2_innest28_S_2, file = "SI/individual_files/D2_lasso_innest28S_2.csv", row.names = FALSE)


#### Lasso regression - D4

# Select the columns for z-score normalization (standardization)
norm_zc_D4 <- c("Betaine","MgCl2","MnCl2","PEG1000","NTP_Conc","Y2_Innest30_Norm","Y2_Innest30_Gel_Norm")
# Standardize the selected numerical features
data_zs_D4 <- data_D4 %>%
  mutate(across(all_of(norm_zc_D4), scale)) %>%
  select(all_of(norm_zc_D4))

predictors_zs_D4 <- names(data_zs_D4)[1:(length(names(data_zs_D4)) - 2)]

lasso_D4_innest30 <- run_lasso_regression_stability_selection(data_zs_D4, predictors_zs_D4, response_variable = "Y2_Innest30_Gel_Norm")
lasso_D4_innest30_Q <- run_lasso_regression_stability_selection(data_zs_D4, predictors_zs_D4, response_variable = "Y2_Innest30_Norm")

# Export important features as CSV
write.csv(lasso_D4_innest30, file = "SI/individual_files/D4_lasso_innest30G.csv", row.names = FALSE)
write.csv(lasso_D4_innest30_Q, file = "SI/individual_files/D4_lasso_innest30Q.csv", row.names = FALSE)

#############################################################################################################
# Analysis of fidelity across mutants and selection conditions

# Normality test

for (col in colnames(fidelity_complete_df)[-1]) {  # Skip the first column
  if (is.numeric(fidelity_complete_df[[col]])) {
    p_value <- shapiro.test(fidelity_complete_df[[col]])$p.value

    if (p_value < 0.05) {
      cat("Column:", col, "is not normally distributed (p-value:", p_value, ")\n")
    } else {
      cat("Column:", col, "is normally distributed (p-value:", p_value, ")\n")
    }
  }
}

# Load required package
library(tidyr)
library(dplyr)
library(tidyverse)
library(ggpubr)
library(rstatix)

# Split data into separate dataframes for each metric
data_df <- fidelity_enriched   #fidelity_complete_df  , fidelity_enriched

del_data <- data_df %>%
  select(ID, starts_with("deletions"))
del_data <- del_data[c("ID","deletions_lib0","deletions_lib2","deletions_lib5","deletions_lib7","deletions_lib8","deletions_lib11")]
ins_data <- data_df %>%
  select(ID, starts_with("insertions"))
ins_data <- ins_data[c("ID","insertions_lib0","insertions_lib2","insertions_lib5","insertions_lib7","insertions_lib8","insertions_lib11")]
subs_data <- data_df %>%
  select(ID, starts_with("substitutions"))
subs_data <- subs_data[c("ID","substitutions_lib0","substitutions_lib2","substitutions_lib5","substitutions_lib7","substitutions_lib8","substitutions_lib11")]


data <- del_data   #del_data  ins_data   subs_data

data <- data %>%
  pivot_longer(-ID, names_to = "selection", values_to = "score")

data %>%
  group_by(selection) %>%
  get_summary_stats(score, type = "common")

ggboxplot(data, x = "selection", y = "score", add = "jitter")

# friedman test
res.fried <- data %>% friedman_test(score ~ selection |ID)
res.fried

pwc <- data %>%
  wilcox_test(score ~ selection, paired = TRUE, p.adjust.method = "bonferroni")
pwc

library(rstatix)

# Extract significant comparisons based on adjusted p-values
significant_comparisons <- pwc %>%
  filter(p.adj < 0.05)

# Print the significant comparisons
print(significant_comparisons, n = Inf)

# Visualization: box plots with p-values
pwc <- pwc %>% add_xy_position(x = "selection")
ggboxplot(data, x = "selection", y = "score", add = "point") +
  stat_pvalue_manual(pwc, hide.ns = TRUE) +
  labs(
    subtitle = get_test_label(res.fried,  detailed = TRUE),
    caption = get_pwc_label(pwc)
  )
```
