## Supplementary_information, scripts and analyses for "Navigating directed evolution efficiently: optimizing selection conditions and selection output analysis": S4_DOE_NGS_Analysis.ipynb

```
# %% [markdown]
# KOD DNAP DIRECTED EVOLUTION NGS DATA ANALYSIS
This notebook contains a simple workflow that will: 
- Demultiplex a large MiSeq dataset based on experimental IDs
- Trim and filter reads by length
- Translate reads to protein sequences
- Identify and count unique sequences pre- and post-selection
- Calculate enrichment and fitness scores of unique sequences post-selection
- Perform a simple statistical test of the enrichment scores
- Identify specific mutations of top enriched candidates
- Visualise the diversity per position in each selection dataset by calculating entropy/column of a MSA

# %% [code]
#Loading packages
using FASTX
using BioSequences
using BioAlignments
using BioAlignments.BioSymbols
using StatsBase
using DataFrames
using Distributions
using Random
using Statistics
using CSV
using PlotlyJS


# %% [code]
#Change path to access datasets more efficiently
cd("C://Users//PH//OneDrive - KU Leuven//PhD//Polymerase_Landscape_Mapping//05_Writing//01_Manuscripts//Selection_landscape_mapping//SI//individual_files//NGS_Analysis//")


# %% [code]
# loading FASTA files and extract reads matching experimental ID
function load_fasta_file_trim(file_name::String,var_id1, var_id2, bar_1, bar_2)
    seqs = Tuple{LongSequence{DNAAlphabet{4}}, LongSequence{DNAAlphabet{4}}}[]
    reader = open(FASTA.Reader, file_name)
    for record in reader
        seq = FASTX.FASTA.sequence(record)
        
        pos1 = findfirst(ExactSearchQuery(var_id1), LongDNA{4}(seq))
        pos2 = findfirst(ExactSearchQuery(var_id2), LongDNA{4}(seq))
        pos3 = findfirst(ExactSearchQuery(bar_1), LongDNA{4}(seq))
        pos4 = findfirst(ExactSearchQuery(bar_2), LongDNA{4}(seq))
        if isnothing(pos1) || isnothing(pos2) || isnothing(pos3) || isnothing(pos4)
            continue
        end    
        trimmed_seq = seq[pos1.start:pos2.stop]
        mut_codons = seq[pos3.stop+1:pos4.start-1]       
        push!(seqs, (LongDNA{4}(trimmed_seq),(LongDNA{4}(mut_codons)))) 
    end
    close(reader)
    return seqs
end

# array to fasta file export
function array_to_fasta_file(file_name::String, DATA)
    reader = open(FASTA.Writer, file_name)
        for i in 1:size(DATA)[1]
            seq = DATA[i]
            record = FASTA.Record("Seq$i", seq)
            write(reader, record)
        end
    close(reader)
end

# %% [code]
# Reference sequence based on the expected trimmed wildtype sequence - will be used for identification of mutations
#F1aa_ref = aa"YLDFRSLYP"
F1aa_ref = aa"LD"
codon_number_start = 403

libR0_ID = dna"atgGCTACTCGATGGAAGATG"
lib2_ID = dna"atgGCTACTCGATGGAAGATG"
lib5_ID = dna"tgaGCTACTCGATGGAAGATG"
lib7_ID = dna"gatGCTACTCGATGGAAGATG"
lib8_ID = dna"actGCTACTCGATGGAAGATG"
lib11_ID =	dna"cagGCTACTCGATGGAAGATG"
lib_all = dna"CTaGGAGACCTCCTAGAGG"

mut_1 = dna"AACATAGTGTAC"
mut_2 = dna"TTTAGATCCCTG"

# %% [code]
#Loading sequences matching IDs 
lib0 = load_fasta_file_trim("D1_R0_preprocessed.fasta", libR0_ID, lib_all, mut_1, mut_2)
lib2 = load_fasta_file_trim("D1_selections_preprocessed.fasta", lib2_ID, lib_all, mut_1, mut_2)
lib5 = load_fasta_file_trim("D1_selections_preprocessed.fasta", lib5_ID, lib_all, mut_1, mut_2)
lib7 = load_fasta_file_trim("D1_selections_preprocessed.fasta", lib7_ID, lib_all, mut_1, mut_2)
lib8 = load_fasta_file_trim("D1_selections_preprocessed.fasta", lib8_ID, lib_all, mut_1, mut_2)
lib11 = load_fasta_file_trim("D1_selections_preprocessed.fasta", lib11_ID, lib_all, mut_1, mut_2)

# %% [code]
# Translating to protein 
function dna_2_aa(DNA_seqs::Vector{Tuple{LongSequence{DNAAlphabet{4}}, LongSequence{DNAAlphabet{4}}}})
    aa_seqs = []
    for i in 1:length(DNA_seqs)
        rna_seq = convert(LongRNA{2}, DNA_seqs[i][2])
        if length(rna_seq) % 3 == 0 
            push!(aa_seqs,BioSequences.translate(rna_seq, allow_ambiguous_codons=true, alternative_start=false))  
        end
    end
    return aa_seqs
end

# %% [code]
aa_lib0 = dna_2_aa(lib0)
aa_lib2 = dna_2_aa(lib2)
aa_lib5 = dna_2_aa(lib5)
aa_lib7 = dna_2_aa(lib7)
aa_lib8 = dna_2_aa(lib8)
aa_lib11 = dna_2_aa(lib11)

# %% [code]
# number of sequences post filtering and translating
println(" aa_lib0=",length(aa_lib0)," aa_lib2=",length(aa_lib2)," aa_lib5=",length(aa_lib5)," aa_lib7=",length(aa_lib7)," aa_lib8=",length(aa_lib8)," aa_lib11=",length(aa_lib11))
# number of unique sequences
println(" aa_lib0=",length(keys(countmap(aa_lib0)))," aa_lib2=",length(keys(countmap(aa_lib2)))," aa_lib5=",length(keys(countmap(aa_lib5)))," aa_lib7=",length(keys(countmap(aa_lib7)))," aa_lib8=",length(keys(countmap(aa_lib8)))," aa_lib11=",length(keys(countmap(aa_lib11))))
# number of unique matching sequences
println(length(intersect(aa_lib0,aa_lib2,aa_lib5,aa_lib7,aa_lib8,aa_lib11)))

# %% [code]
using Combinatorics
# Define the set of amino acid letters
amino_acids = "ARNDCEQGHILKMFPSTWYV"
# Function to generate two-letter combinations
function generate_two_letter_combinations(amino_acids)
    two_letter_combinations = [string(c1, c2) for c1 in amino_acids, c2 in amino_acids]
    return filter(x -> length(x) == 2, two_letter_combinations)
end

# Generate two-letter combinations
two_letter_combinations = generate_two_letter_combinations(amino_acids)
two_letter_combinations

unique_aa_r0 = intersect(string.(aa_lib0),two_letter_combinations)
unique_aa_2 = intersect(string.(aa_lib2),two_letter_combinations)
unique_aa_5 = intersect(string.(aa_lib5),two_letter_combinations)
unique_aa_7 = intersect(string.(aa_lib7),two_letter_combinations)
unique_aa_8 = intersect(string.(aa_lib8),two_letter_combinations)
unique_aa_11 = intersect(string.(aa_lib11),two_letter_combinations)

print("R0 unique viable aa combinations: ", length(unique_aa_r0), "/400 ")
print("Sel 2 unique viable aa combinations: ", length(unique_aa_2), "/400 ")
print("Sel 5 unique viable aa combinations: ", length(unique_aa_5), "/400 ")
print("Sel 7 unique viable aa combinations: ", length(unique_aa_7), "/400 ")
print("Sel 8 unique viable aa combinations: ", length(unique_aa_8), "/400 ")
print("Sel 11 unique viable aa combinations: ", length(unique_aa_11), "/400 ")

# %% [code]
#Identifying and counting unique sequences 
#calculating their frequency (based on the total number of reads)
#calculating enrichment of sequences from R0 to R1 of selection
function etest(data::DataFrame, iside::Bool=false)
    pvalues = []
    for i in 1:size(data)[1]
        k1 = data[!,:Count_R0][i]
        k2 = data[!,:Count_R1][i]
        sum_r0 = sum(data[!,:Count_R0])
        sum_r1 = sum(data[!,:Count_R1])
        pvalue = testPoissonSignificance(k1,k2,sum_r0,sum_r1)
        push!(pvalues,pvalue)
    end
  return pvalues
end
function testPoissonSignificance(k1::Int64, k2::Int64, n1::Int64, n2::Int64, d::Float64=0.0, iside::Bool=false)
  elhatk = (k1+k2)/(n1+n2)-d*n1/(n1+n2)
  var = (k1/ (n1^2) + k2/(n2^2))
  t_k1k2 = (k1/n1-k2/n2-d)/sqrt(Complex(var)) #?
  pvalue=poistest(n1, n2, elhatk, t_k1k2, d, iside)

  return pvalue
end
function poistest(n1::Int64, n2::Int64, elhatk, t_k1k2, d::Float64=0.0, iside::Bool=false)

  # computing estimates of el1*n1 and el2*n2 under H_0
  pvalue=0 
  elhat1=n1*(elhatk+d)
  elhat2 = n2*elhatk

  # computing the modes 
    i1mode = floor(elhat1)
    i2mode = floor(elhat2)

  # initializing the probability at the i1mode
    pi1mode = pdf(Poisson(elhat1), i1mode)
    pi1 = pi1mode

  # initializing the probability at the i2mode
    pi2mode = pdf(Poisson(elhat2), i2mode)
    for i1 = i1mode:1000  
        if pi1 >= 1e-7       
          pvalue=sumi2(n1, n2, elhat2, t_k1k2, i1, pi1, i2mode, pi2mode, d, pvalue, iside)
          pi1 = elhat1*pi1/(i1+1)
        end
    end 

  # Label #1 
  i1 = i1mode-1
  pi1 = pi1mode
    pi1 = i1mode*pi1/elhat1

    for i1 = i1mode-1:-1: 0
      if pi1 >= 1e-7 
          pvalue=sumi2(n1, n2, elhat2, t_k1k2, i1, pi1, i2mode, pi2mode, d, pvalue, iside)
          pi1 = i1*pi1/elhat1
      end
    end
  
  return pvalue
end
function sumi2(n1, n2, elhat2, t_k1k2, i1, pi1, i2mode, pi2mode, d, pvalue, iside)
  pvalue = pvalue
  pi2 = pi2mode

    for i2 = i2mode:1000
        if pi2 >= 1.0e-07
            elhati1 = 1.0e0*i1/n1
            elhati2 = 1.0e0*i2/n2
      
            diffi = elhati1 - elhati2 - d
            var = 1.0e0*elhati1/n1 + 1.0e0*elhati2/n2
    
        if iside == true     
        if 1.0e0*i1/n1 - 1.0e0*i2/n2 <= d
                t_i1i2 = 0.0e0
        else
                t_i1i2 = diffi/sqrt(var)
        end
            
        if t_i1i2 >= t_k1k2 
          pvalue += pi1*pi2
        end
          
      elseif iside == false
              if abs(1.0e0*i1/n1 - 1.0e0*i2/n2) <= d 
                t_i1i2 = 0.0e0
              else
                t_i1i2 = diffi/sqrt(var)
        end
            
        if abs(t_i1i2) >= abs(t_k1k2) 
          pvalue += pi1*pi2
        end
      end
    end
          
    pi2 = elhat2*pi2/(i2+1.0e0)
    end

  i2 = i2mode-1
        pi2 = pi2mode
        pi2 = i2mode*pi2/elhat2

        for i2 = i2mode-1:-1: 0
          if pi2 >= 1.0e-07 
            elhati1 = 1.0e0*i1/n1
            elhati2 = 1.0e0*i2/n2
            diffi = elhati1 - elhati2 - d
            var = (1.0e0*elhati1/n1 + 1.0e0*elhati2/n2)

            if iside == true 
              if 1.0e0*i1/n1 - 1.0e0*i2/n2 <= d 
                t_i1i2 = 0.0e0
              else
                t_i1i2 = diffi/sqrt(var)
              end 

              if t_i1i2 >= t_k1k2 
              pvalue += pi1*pi2
            end

          elseif iside == false     
              if abs(1.0e0*i1/n1 - 1.0e0*i2/n2) <= d 
                t_i1i2 = 0.0e0
              else
                t_i1i2 = diffi/sqrt(var)
              end 

            if abs(t_i1i2) >= abs(t_k1k2) 
              pvalue += pi1*pi2
            end
            end 
        end
          pi2 = i2*pi2/elhat2

        end

  return pvalue
end

function mut_count(aa_seqs)
    output=[]
    total = length(aa_seqs)
    unique_counts = countmap(aa_seqs)
    for (keys,vals) in unique_counts
        freq = vals/total
        push!(output, (keys,vals,freq))
    end
    output = DataFrame([[output[k][kk] for k in 1:length(output)] for kk in 1:length(output[1])], [:Seq, :Count, :Freq])
    return sort!(output, [:Freq],rev=true)
end

function enrichment_complex(R0_freq,R1_freq,R0_counts,R1_counts)
    EC = (R1_freq .- R0_freq).*((R0_counts.+R1_counts)/2)
    return EC
end

function find_muts(enrichment_table,ref_seq,codon_number_start)
    mut_array = []
    for j in 1:size(enrichment_table)[1]
        muts = []
        if length(ref_seq) == length(enrichment_table[j,1])
            for i in 1:length(ref_seq)
                if ref_seq[i] != enrichment_table[j,1][i]
                    mut = string(ref_seq[i],"-",i+codon_number_start-1,"-",enrichment_table[j,1][i],";")
                    append!(muts,mut)
                end
            end
            push!(mut_array,join(muts))
        else
            push!(mut_array,"InDel/fs")
        end
    end
    return DataFrame([mut_array], [:Mutations])
end

function mut_enrichment(R0,R1,ref_seq, sel_name)
    #counting sequences
    r0 = mut_count(R0)
    r1 = mut_count(R1)
    rename!(r0,:Count => :Count_R0, :Freq => :Freq_R0)
    rename!(r1,:Count => :Count_R1, :Freq => :Freq_R1)
    
    #merging dataframes of r0 and r1 sequences
    merged = innerjoin(r0, r1, on = :Seq)
    
    #calculating enrichment and enrichment complex of sequences
    enrichment = log.(merged[!,:Freq_R1] ./ merged[!,:Freq_R0])
    ec = enrichment_complex(merged[!,:Freq_R0],merged[!,:Freq_R1],merged[!,:Count_R0],merged[!,:Count_R1])
    merged[!,:Enrichment] = enrichment
    merged[!,:EC] = ec

    #E-Test
    merged[!,:"p-val"] = etest(merged)
    #calculating fitness scores
    if ref_seq in merged[!,:Seq]
        enrichment_wt = merged[in([ref_seq]).(merged.Seq), :][!,:Enrichment][1]
        fitness = merged[!,:Enrichment].-enrichment_wt
        merged[!,:Fitness] = fitness
        #converting sequences to list of mutations
        mutations = find_muts(merged,ref_seq,codon_number_start)
        output = hcat(mutations,merged)
        sort!(output,[:Enrichment],rev=true) ## sorting output by Enrichment, this can be modified to any other metric
        sel_names = ["Mutations_$sel_name", "Seq", "Count_R0_$sel_name", "Freq_R0_$sel_name", "Count_R1_$sel_name", "Freq_R1_$sel_name", "Enrichment_$sel_name", "EC_$sel_name", "p-val_$sel_name", "Fitness_$sel_name"]
        rename!(output, sel_names)
        return output
    elseif ref_seq ∉ merged[!,:Seq]
        println("*** Wild type sequence not found in dataset - Fitness was not calculated ***")
        #converting sequences to list of mutations
        mutations = find_muts(merged,ref_seq,codon_number_start)
        output = hcat(mutations,merged)
        sort!(output,[:Enrichment],rev=true) ## sorting output by Enrichment, this can be modified to any other metric
        sel_names = ["Mutations_$sel_name", "Seq", "Count_R0_$sel_name", "Freq_R0_$sel_name", "Count_R1_$sel_name", "Freq_R1_$sel_name", "Enrichment_$sel_name", "EC_$sel_name", "p-val_$sel_name"]
        rename!(output, sel_names)
        return output
    end 
end

function rename_df(dataframe, name, col_exceptions)
  new_dataframe = copy(dataframe) 
  old_names = names(new_dataframe)
  deleteat!(old_names, col_exceptions)
  new_names = []
  for i in 1:length(old_names)
      new_name = old_names[i]*"_$name"
      push!(new_names,new_name)
  end
  rename!(new_dataframe, old_names .=> new_names)
  return new_dataframe
end


# %% [code]
enrichment_2 = mut_enrichment(aa_lib0, aa_lib2, F1aa_ref, "sel2")
enrichment_5 = mut_enrichment(aa_lib0, aa_lib5, F1aa_ref, "sel5")
enrichment_7 = mut_enrichment(aa_lib0, aa_lib7, F1aa_ref, "sel7")
enrichment_8 = mut_enrichment(aa_lib0, aa_lib8, F1aa_ref, "sel8")
enrichment_11 = mut_enrichment(aa_lib0, aa_lib11, F1aa_ref, "sel11")

sel_enrichment = outerjoin(enrichment_2, enrichment_5, enrichment_7, enrichment_8, enrichment_11,on = :Seq)
sel_enrichment_match = innerjoin(enrichment_2, enrichment_5, enrichment_7, enrichment_8, enrichment_11,on = :Seq)
show(sel_enrichment, allcols = true)
show(sel_enrichment_match, allcols = true)

#CSV.write("D1_enrichment_df.csv", sel_enrichment)
#CSV.write("D1_enrichment_match_df.csv", sel_enrichment_match)

# %% [code]
function threshold_extract(data, condition, threshold, ID)
    if condition == "enriched"
        output = data[(data[!,"Enrichment_$ID"] .> threshold) .& (data[!,"p-val_$ID"] .< 0.01),:] #0.05 pval for D4
    elseif  condition == "depleted"
        output = data[(data[!,"Enrichment_$ID"] .< threshold) .& (data[!,"p-val_$ID"] .< 0.01),:] #0.05 pval for D4
    elseif condition == "count"
        output = data[(data[!,"Count_R1_$ID"] .> threshold) .& (data[!,"p-val_$ID"] .< 0.05) .& (data[!,"Enrichment_$ID"] .> 0),:]
    elseif condition == "pval"
        output = data[(data[!,"p-val_$ID"] .< threshold),:]
        sort!(output,["Enrichment_$ID"],rev=true) 
    end       
    return output
end

# %% [code]
enriched2 = threshold_extract(enrichment_2,"enriched", 0, "sel2")
enriched5 = threshold_extract(enrichment_5,"enriched", 0, "sel5")
enriched7 = threshold_extract(enrichment_7,"enriched", 0,  "sel7")
enriched8 = threshold_extract(enrichment_8,"enriched", 0, "sel8")
enriched11 = threshold_extract(enrichment_11,"enriched", 0, "sel11")

depleted2 = threshold_extract(enrichment_2,"depleted", 0,  "sel2")
depleted5 = threshold_extract(enrichment_5,"depleted", 0, "sel5")
depleted7 = threshold_extract(enrichment_7,"depleted", 0, "sel7")
depleted8 = threshold_extract(enrichment_8,"depleted", 0, "sel8")
depleted11 = threshold_extract(enrichment_11,"depleted",0, "sel11")

sig2 = threshold_extract(enrichment_2,"pval", 0.01, "sel2")
sig5 = threshold_extract(enrichment_5,"pval", 0.01, "sel5")
sig7 = threshold_extract(enrichment_7,"pval", 0.01,  "sel7")
sig8 = threshold_extract(enrichment_8,"pval", 0.01, "sel8")
sig11 = threshold_extract(enrichment_11,"pval", 0.01, "sel11")

enriched_muts_df = innerjoin(enriched2[!,[2,7]],enriched5[!,[2,7]],enriched7[!,[2,7]],enriched8[!,[2,7]],enriched11[!,[2,7]], on= :Seq)
depleted_muts_df = innerjoin(depleted2[!,[2,7]],depleted5[!,[2,7]],depleted7[!,[2,7]],depleted8[!,[2,7]],depleted11[!,[2,7]], on= :Seq)
top_muts_df = innerjoin(enriched2[!,[2,7,8,9,10]],enriched5[!,[2,7,8,9,10]],enriched7[!,[2,7,8,9,10]],enriched8[!,[2,7,8,9,10]],enriched11[!,[2,7,8,9,10]], on= :Seq)
enriched_muts_df = outerjoin(enriched2[!,[2,7]],enriched5[!,[2,7]],enriched7[!,[2,7]],enriched8[!,[2,7]],enriched11[!,[2,7]], on= :Seq)
depleted_muts_df = outerjoin(depleted2[!,[2,7]],depleted5[!,[2,7]],depleted7[!,[2,7]],depleted8[!,[2,7]],depleted11[!,[2,7]], on= :Seq)
top_muts_df = outerjoin(enriched2[!,[2,7,8,9,10]],enriched5[!,[2,7,8,9,10]],enriched7[!,[2,7,8,9,10]],enriched8[!,[2,7,8,9,10]],enriched11[!,[2,7,8,9,10]], on= :Seq)


# %% [code]
sig_muts_df = innerjoin(sig2[!,[2,7,8,9,10]],sig5[!,[2,7,8,9,10]],sig7[!,[2,7,8,9,10]],sig8[!,[2,7,8,9,10]],sig11[!,[2,7,8,9,10]], on= :Seq)
#CSV.write("D1_sig_muts_matched.csv", sig_muts_df)

# %% [markdown]
### 2D Plot 

# %% [code]
function extract_coord(df)
    y = []
    x = []
    map_enrichment = DataFrame()
    mapped_df = DataFrame()
    mapped_df[!,:MUT2] = []

    df = filter(row -> !(row.Seq == aa""),  df) #remove InDels
    sort!(df,[:Seq],rev=true)

    for i in 1:size(df)[1]
        seq = df[!,:Seq][i]
        push!(y, seq[1])
        push!(x, seq[2])
    end
    map_enrichment[!,:MUT1] = y
    map_enrichment[!,:MUT2] = x
    map_enrichment[!,:Enrichment] = df[!,7]
    map_enrichment = groupby(map_enrichment, :MUT1)
    
    for (k,v) in pairs(map_enrichment)
        temp = deepcopy(v)
        rename!(temp, 3 => string(temp[1,:MUT1]))
        select!(temp, Not([:MUT1]))
        mapped_df = outerjoin(mapped_df, temp, on=:MUT2)
    end
    
    return sort!(mapped_df,[:MUT2],rev=true)

end

# %% [code]
enrichment2_2D = extract_coord(enrichment_2)
CSV.write("enrichment2_2D.csv", enrichment2_2D)

enrichment5_2D = extract_coord(enrichment_5)
CSV.write("enrichment5_2D.csv", enrichment5_2D)

enrichment7_2D = extract_coord(enrichment_7)
CSV.write("enrichment7_2D.csv", enrichment7_2D)

enrichment8_2D = extract_coord(enrichment_8)
CSV.write("enrichment8_2D.csv", enrichment8_2D)

enrichment11_2D = extract_coord(enrichment_11)
CSV.write("enrichment11_2D.csv", enrichment11_2D)

# %% [markdown]
#### Fidelity Analysis

# %% [code]
function pol_fidelity(fasta_seqs, ref::LongSequence{DNAAlphabet{4}}, lib_name::String)
    mismatches = []
    deletions = []
    insertions = []
    bases = []
    model = CostModel(match=0, mismatch=1, insertion=2, deletion=2)
    for i in 1:length(fasta_seqs)
        alg = pairalign(EditDistance(), fasta_seqs[i], ref, model)
        append!(mismatches,count_mismatches(alignment(alg)))
        append!(deletions,count_deletions(alignment(alg)))
        append!(insertions,count_insertions(alignment(alg)))
        append!(bases,length(fasta_seqs[i]))
    end
    total_bases = sum(bases)
    total_errors = Dict{String,Any}()
    total_errors["substitutions"] = sum(mismatches)/total_bases
    total_errors["deletions"] = sum(deletions)/total_bases
    total_errors["insertions"] = sum(insertions)/total_bases
    total_errors["ID"] = lib_name
    total_errors["bases_sequenced"] = total_bases
    return DataFrame(total_errors)
end

function pol_fidelity_stats(data::Vector{Tuple{LongSequence{DNAAlphabet{4}}, LongSequence{DNAAlphabet{4}}}},ref::LongSequence{DNAAlphabet{4}}, lib_name::String)
    seqs = []
    background = []
    for i in 1:length(data)
        seq = data[i][1]
        pos1 = findfirst(ExactSearchQuery(mut_1), seq)
        pos2 = findfirst(ExactSearchQuery(mut_2), seq)
        if isnothing(pos1) || isnothing(pos2)
            continue
        end
        diversity = seq[pos1.stop+1:pos2.start-1] #removing diversity region

        if length(diversity) % 3 == 0  #if divisible by 3
            seq_trimmed = seq[4:pos1.stop]*seq[pos2.start:length(seq)]
            push!(seqs,seq_trimmed)

        else
            push!(background,"fs")  
        end
    end
    total_errors = pol_fidelity(seqs,ref,lib_name)
    total_errors[!,:"frame-shifts"] .= length(background)/length(data)
    return DataFrame(total_errors)
end

# %% [code]
ref_seq = dna"GCTACTCGATGGAAGATGCGAAGGTCACATACGAGCTTGGGAAGGAGTTCCTTCCGATGGAGGCCCAGCTTTCTCGCTTAATCGGCCAGTCCCTCTGGGACGTCTCCCGCTCCAGCACTGGCAACCTCGTTGAGTGGTTCCTCCTCAGGAAGGCCTATGAGAGGAATGAGCTGGCCCCGAACAAGCCCGATGAAAAGGAGCTGGCCAGAAGACGGCAGAGCTATGAAGGAGGCTATGTAAAAGAGCCCGAGAGAGGGTTGTGGGAGAACATAGTGTACTTTAGATCCCTGTACCCCTCAATCATCATCACCCACAACGTCTCGCCGGATACGCTCAACAGAGAAGGATGCAAGGAATATGACGTTGCCCCACAGGTCGGCCACCGCTTCTGCAAGGACTTCCCAGGATTTATCCCGAGCCTGCTAGGAGACCTCCTAGAGG"
mut_1 = dna"GAGAACATAGTGTAC"
mut_2 = dna"TTTAGATCCCTGTAC"
lib0_fidelity = pol_fidelity_stats(lib0, ref_seq, "lib0")
lib2_fidelity = pol_fidelity_stats(lib2, ref_seq, "lib2")
lib5_fidelity = pol_fidelity_stats(lib5, ref_seq, "lib5")
lib7_fidelity = pol_fidelity_stats(lib7, ref_seq, "lib7")
lib8_fidelity = pol_fidelity_stats(lib8, ref_seq, "lib8")
lib11_fidelity = pol_fidelity_stats(lib11, ref_seq, "lib11")
lib_fidelity = vcat(lib0_fidelity,lib2_fidelity,lib5_fidelity,lib7_fidelity,lib8_fidelity,lib11_fidelity)

#CSV.write("D1_Fidelity_overall_type.csv", lib_fidelity)

# %% [code]
# mutant-specific fidelity

function mut_fidelity(data::Vector{Tuple{LongSequence{DNAAlphabet{4}}, LongSequence{DNAAlphabet{4}}}},ref::LongSequence{DNAAlphabet{4}}, lib_name::String)
    seqs = []
    trimmed_seqs = []
    muts = []
    sel_id = []
    for i in 1:length(data)
        seq = data[i][1]
        pos1 = findfirst(ExactSearchQuery(mut_1), seq)
        pos2 = findfirst(ExactSearchQuery(mut_2), seq)
        if isnothing(pos1) || isnothing(pos2)
            continue
        end

        seq_trimmed = seq[4:pos1.stop]*seq[pos2.start:length(seq)]
        rna_seq = convert(LongRNA{2}, data[i][2])

        if length(rna_seq) % 3 == 0  #if divisible by 3
            mut_trans = BioSequences.translate(rna_seq, allow_ambiguous_codons=false, alternative_start=false)
                push!(seqs,data[i][1])
                push!(trimmed_seqs,seq_trimmed)
                push!(muts,mut_trans)
                push!(sel_id,data[i][1][1:3])            
        end
    end
    df = DataFrame(Sel_ID = sel_id, Seq = seqs, Trimmed_seq = trimmed_seqs, Muts = muts)
    df_mut_sub = groupby(df, :Muts)
    
    fidelity_mutants = DataFrame()
    for (k,v) in pairs(df_mut_sub)
        df = pol_fidelity(v[:,:Trimmed_seq], ref_seq, String(k.Muts))
        append!(fidelity_mutants,df)
    end
    fidelity_mutants[!,:Sel] = fill(lib_name, size(fidelity_mutants)[1])
    return fidelity_mutants
end

# %% [code]
ref_seq = dna"GCTACTCGATGGAAGATGCGAAGGTCACATACGAGCTTGGGAAGGAGTTCCTTCCGATGGAGGCCCAGCTTTCTCGCTTAATCGGCCAGTCCCTCTGGGACGTCTCCCGCTCCAGCACTGGCAACCTCGTTGAGTGGTTCCTCCTCAGGAAGGCCTATGAGAGGAATGAGCTGGCCCCGAACAAGCCCGATGAAAAGGAGCTGGCCAGAAGACGGCAGAGCTATGAAGGAGGCTATGTAAAAGAGCCCGAGAGAGGGTTGTGGGAGAACATAGTGTACTTTAGATCCCTGTACCCCTCAATCATCATCACCCACAACGTCTCGCCGGATACGCTCAACAGAGAAGGATGCAAGGAATATGACGTTGCCCCACAGGTCGGCCACCGCTTCTGCAAGGACTTCCCAGGATTTATCCCGAGCCTGCTAGGAGACCTCCTAGAGG"

lib0_mut_fidelity = mut_fidelity(lib0, ref_seq, "lib0")
lib2_mut_fidelity = mut_fidelity(lib2,ref_seq, "lib2")
lib5_mut_fidelity = mut_fidelity(lib5,ref_seq, "lib5")
lib7_mut_fidelity = mut_fidelity(lib7,ref_seq, "lib7")
lib8_mut_fidelity = mut_fidelity(lib8,ref_seq, "lib8")
lib11_mut_fidelity = mut_fidelity(lib11,ref_seq, "lib11")

lib0_mut_fidelity_rn = rename_df(lib0_mut_fidelity, "lib0", 1)
lib2_mut_fidelity_rn = rename_df(lib2_mut_fidelity, "lib2", 1)
lib5_mut_fidelity_rn = rename_df(lib5_mut_fidelity, "lib5", 1)
lib7_mut_fidelity_rn = rename_df(lib7_mut_fidelity, "lib7", 1)
lib8_mut_fidelity_rn = rename_df(lib8_mut_fidelity, "lib8", 1)
lib11_mut_fidelity_rn = rename_df(lib11_mut_fidelity, "lib11", 1)

muts_fidelity_sels = outerjoin(lib0_mut_fidelity,lib2_mut_fidelity,lib5_mut_fidelity,lib7_mut_fidelity,lib8_mut_fidelity,lib11_mut_fidelity, on= :ID)
#CSV.write("D1_Muts_Fidelity_All.csv", muts_fidelity_sels)
muts_fidelity_sels_matched = innerjoin(lib0_mut_fidelity,lib2_mut_fidelity,lib5_mut_fidelity,lib7_mut_fidelity,lib8_mut_fidelity,lib11_mut_fidelity, on= :ID)
#CSV.write("D1_Muts_Fidelity_Matched.csv", muts_fidelity_sels_matched)

# %% [code]
# Fidelity of top significantly enriched mutants
sig_muts_list = string.(sig_muts_df[1:20,:Seq])
sig_muts_rest_list = string.(sig_muts_df[21:end,:Seq])

sig_muts_fidelity = muts_fidelity_sels_matched[in.(muts_fidelity_sels_matched.ID,Ref(sig_muts_list)),:]
rest_sig_muts_fidelity = muts_fidelity_sels_matched[in.(muts_fidelity_sels_matched.ID,Ref(sig_muts_rest_list)),:]
rest_muts_fidelity = muts_fidelity_sels_matched[.!in.(muts_fidelity_sels_matched.ID,Ref(sig_muts_list)),:]

# Adding enrichment scores to fidelity tables for correlation analysis
sig_muts_df_enrichments = sig_muts_df[:,[col for col in names(sig_muts_df) if startswith(col, "Seq") || startswith(col, "Enrichment_")]]
sig_muts_df_enrichments[!, :ID] .= string.(sig_muts_df_enrichments.Seq)

sig_muts_fidelity_enrichment = innerjoin(sig_muts_fidelity,sig_muts_df_enrichments,on=:ID)
rest_sig_muts_fidelity_enrichment = innerjoin(rest_sig_muts_fidelity,sig_muts_df_enrichments,on=:ID)
#CSV.write("D1_sig_muts_fidelity.csv", sig_muts_fidelity_enrichment)

# %% [code]
#adding enrichment values to fidelity scores
#sel_enrichment_match.ID = string.(sel_enrichment_match.Seq)
enrichment_fidelity_matched = innerjoin(sel_enrichment_match, muts_fidelity_sels_matched, on = :ID)
enrichment_fidelity_matched = select(enrichment_fidelity_matched, r"^(ID|Enr|Freq|bases|ins|del|subs).*$")
# substracting R0 values from fidelity scores
enrichment_fidelity_matched[!, [:insertions_lib0,:insertions_lib2,:insertions_lib5,:insertions_lib7,:insertions_lib8,:insertions_lib11]] .-= enrichment_fidelity_matched[!, :insertions_lib0]
enrichment_fidelity_matched[!, [:deletions_lib0,:deletions_lib2,:deletions_lib5,:deletions_lib7,:deletions_lib8,:deletions_lib11]] .-= enrichment_fidelity_matched[!, :deletions_lib0]
enrichment_fidelity_matched[!, [:substitutions_lib0,:substitutions_lib2,:substitutions_lib5,:substitutions_lib7,:substitutions_lib8,:substitutions_lib11]] .-= enrichment_fidelity_matched[!, :substitutions_lib0]

# %% [code]
CSV.write("D1_enrichment_fidelity_matched.csv", enrichment_fidelity_matched)

# %% [markdown]
### Top mutants in-depth analysis of fidelity

# %% [code]
function seq_subset_to_dataframe(data::Vector{Tuple{LongSequence{DNAAlphabet{4}}, LongSequence{DNAAlphabet{4}}}}, list::Vector{String})
    seqs = Tuple{String, LongSequence{AminoAcidAlphabet}}[]
    for i in 1:length(data)
        rna_seq = convert(LongRNA{2}, data[i][2])
        if length(rna_seq) % 3 == 0  #if divisible by 3
            mut_trans = BioSequences.translate(rna_seq, allow_ambiguous_codons=false, alternative_start=false)
            if string(mut_trans) in list 
                seq = data[i][1]
                pos1 = findfirst(ExactSearchQuery(dna"GAGAACATAGTGTAC"), seq)
                pos2 = findfirst(ExactSearchQuery(dna"TTTAGATCCCTGTAC"), seq)
                if isnothing(pos1) || isnothing(pos2)
                    continue
                end
                trimmed_seq = seq[4:pos1.stop] * seq[pos2.start:length(seq)]
                if length(trimmed_seq) == 441  # length of ref_seq without mut site
                    push!(seqs, (string(trimmed_seq), mut_trans))
                end
            end
        end
    end
    # Create a DataFrame from the seqs array
    df = DataFrame(Seq = [t[1] for t in seqs], ID = [t[2] for t in seqs])

    return df
end


# %% [code]
function error_type_counting(data::DataFrame, ref_seq::String, sel_name::String)
    grouped_test = groupby(data, :ID)

    # Define the error types
    error_types = [
        "A->G", "G->A", "T->C", "C->T", "A->T", "T->A",
        "A->C", "C->A", "T->G", "G->T", "C->G", "G->C"
    ]

    ids = String[]
    total_bases = Int[]
    transitions = Int[]
    transversions = Int[]
    error_counts = Vector{Dict{String, Int}}()

    for group in grouped_test
        ID = string(group[1, :ID])  # Convert LongAA to String
        subset_df = group[!, :Seq]
        total_bases_group = size(subset_df, 1) * 441  # 441 is the length of each string to calculate total bases examined
        error_counts_group = Dict{String, Int}()
        for error in error_types
            error_counts_group[error] = 0
        end
        for i in 1:size(subset_df, 1)
            sequence = subset_df[i]
            for (t1, t2) in zip(ref_seq, sequence)
                error = string(t1, "->", t2)
                if haskey(error_counts_group, error)
                    error_counts_group[error] += 1
                end
            end
        end
        for error in error_types
            if !haskey(error_counts_group, error)
                error_counts_group[error] = 0
            end
        end

        # Calculate transitions and transversions
        transitions_group = error_counts_group["A->G"] + error_counts_group["G->A"] + error_counts_group["C->T"] + error_counts_group["T->C"]
        transversions_group = sum([error_counts_group[error] for error in error_types if (error ∉ ["A->G", "G->A", "C->T", "T->C"])])

        # Add the results to the vectors
        push!(ids, ID)
        push!(total_bases, total_bases_group)
        push!(transitions, transitions_group)
        push!(transversions, transversions_group)
        push!(error_counts, error_counts_group)
    end

    # Convert the list of dictionaries to a DataFrame
    error_counts_df = DataFrame(values(error_counts))

    # Combine the DataFrames into the final result
    results_df = hcat(
        DataFrame(ID = ids, TotalBases = total_bases, Transitions = transitions, Transversions = transversions),
        error_counts_df
    )
    results_df[!, 3:end] = Float64.(results_df[:, 3:end]) ./ results_df[:, 2]
    results_df = rename_df(results_df, sel_name, 1)
    # Add sel_name to column names
    return results_df
end

# %% [code]
sig_muts_list = string.(sig_muts_df[1:20,:Seq])
lib0_top_seqs = seq_subset_to_dataframe(lib0,sig_muts_list)
lib2_top_seqs = seq_subset_to_dataframe(lib2,sig_muts_list)
lib5_top_seqs = seq_subset_to_dataframe(lib5,sig_muts_list)
lib7_top_seqs = seq_subset_to_dataframe(lib7,sig_muts_list)
lib8_top_seqs = seq_subset_to_dataframe(lib8,sig_muts_list)
lib11_top_seqs = seq_subset_to_dataframe(lib11,sig_muts_list)

ref_seq_string = "GCTACTCGATGGAAGATGCGAAGGTCACATACGAGCTTGGGAAGGAGTTCCTTCCGATGGAGGCCCAGCTTTCTCGCTTAATCGGCCAGTCCCTCTGGGACGTCTCCCGCTCCAGCACTGGCAACCTCGTTGAGTGGTTCCTCCTCAGGAAGGCCTATGAGAGGAATGAGCTGGCCCCGAACAAGCCCGATGAAAAGGAGCTGGCCAGAAGACGGCAGAGCTATGAAGGAGGCTATGTAAAAGAGCCCGAGAGAGGGTTGTGGGAGAACATAGTGTACTTTAGATCCCTGTACCCCTCAATCATCATCACCCACAACGTCTCGCCGGATACGCTCAACAGAGAAGGATGCAAGGAATATGACGTTGCCCCACAGGTCGGCCACCGCTTCTGCAAGGACTTCCCAGGATTTATCCCGAGCCTGCTAGGAGACCTCCTAGAGG"
lib0_error_types = error_type_counting(lib0_top_seqs,ref_seq_string, "R0")
lib2_error_types = error_type_counting(lib2_top_seqs,ref_seq_string, "Sel2")
lib5_error_types = error_type_counting(lib5_top_seqs,ref_seq_string, "Sel5")
lib7_error_types = error_type_counting(lib7_top_seqs,ref_seq_string, "Sel7")
lib8_error_types = error_type_counting(lib8_top_seqs,ref_seq_string, "Sel8")
lib11_error_types = error_type_counting(lib11_top_seqs,ref_seq_string, "Sel11")

top_muts_error_types = innerjoin(lib0_error_types,lib2_error_types,lib5_error_types,lib7_error_types,lib8_error_types,lib11_error_types, on=:ID)

# %% [code]
#CSV.write("D1_top_muts_error_types.csv", top_muts_error_types)


# %% [markdown]
#### Imputing failed selections mutant-specific fidelity

# %% [code]

mut_types = ["deletions", "insertions" , "substitutions"]
output =[]
for i in 1:length(mut_types)
    cols = [col for col in names(muts_fidelity_sels_matched) if startswith(col, mut_types[i])]
    data = muts_fidelity_sels_matched[:, vcat("ID",cols)]
    original_names = names(data[!,Not(:ID)]) 

    # Calculate mean, standard deviation, lower threshold, and upper threshold
    data.mean = mean.(eachrow(data[:, Not(:ID)]))
    data.std = std.(eachrow(data[:, Not(:ID)]))
    data.lower_quantile = quantile.(eachrow(data[:, Not(:ID)]), 0.90)
    data.upper_quantile = quantile.(eachrow(data[:, Not(:ID)]), 0.95)

    # Calculate the range between upper and lower quantiles
    data.range = data.upper_quantile .- data.lower_quantile

    # Generate random values within the specified quantile range
    imputed_columns = Symbol.(mut_types[i].*"_lib" .* string.([1, 3, 4, 6, 9, 10, 12]))

    for i in 1:length(imputed_columns)
        data[!, imputed_columns[i]] = [
            lower + range * rand()
            for (lower, range) in zip(data.lower_quantile, data.range)
        ]
    end
    push!(output,data)
end

data_compiled = outerjoin(output[1],output[2],output[3],on=:ID, makeunique=true)
# Display the modified DataFrame
show(data_compiled, allcols=true)
#CSV.write("D1_Mutant_Fidelity_FailedSelections_Engineered_1.csv", data_compiled)


# %% [markdown]
#### Sequencing Depth Analysis

# %% [code]
function random_sets(data, percentage_depth, trials)
    sets = []
    for i in 1:trials
        N = Int(percentage_depth*400)
        indices = randperm(length(data))[1:N]
        subset = data[indices]
        push!(sets,subset)
    end
    return sets
end

function enrichment_sets(sets_r0, sets_r1, ref, mut, sel_name)
    enrichment_scores = []
    for i in 1:size(sets_r0)[1]
        enrich_set = mut_enrichment(sets_r0[i], sets_r1[i], ref, sel_name)
        enrichment_score = enrich_set[enrich_set[!, "Mutations_$sel_name"] .== mut, "Enrichment_$sel_name"]
        append!(enrichment_scores, enrichment_score)
    end
    avg_score = mean(enrichment_scores)
    std_score = std(enrichment_scores)
    return (avg_score,std_score)
end


function interval_mutant_stats(r0, r1, interval, trials, ref, mut, sel_name)
    percentage_scores = []
    trial_col = "Average_Enrichment_N$trials"
    for i in 1:length(interval)
        r0_sets = random_sets(r0,interval[i],trials)
        r1_sets = random_sets(r1,interval[i],trials)
        avg_scores = enrichment_sets(r0_sets,r1_sets, ref, mut, sel_name)
        push!(percentage_scores,(interval[i],avg_scores[1],avg_scores[2]))
    end
    percentage_scores = DataFrame(percentage_scores)
    rename!(percentage_scores, :1 =>:"Sequencing_Depth", :2=>trial_col, :3=>:"STD")
    return percentage_scores
end

function merge_dataframes(dfs)
    merged_df = dfs[1]
    for df in dfs[2:end]
        merged_df = outerjoin(merged_df, df, on=:Seq, makeunique=true)
    end
    return merged_df
end

function set_enrichment_stats(r0, r1, depth, trials, ref, sel_name)
    enrichment_dfs, rank_dfs, freq_dfs, top_count_dfs, rest_count_dfs = [], [], [], [], []
    trial_col = "Average_Enrichment_N$trials"
    r0_sets = random_sets(r0,depth,trials)
    r1_sets = random_sets(r1,depth,trials)
    for i in 1:trials 
       mut_enrich_df = mut_enrichment(r0_sets[i], r1_sets[i], F1aa_ref, sel_name)
       mut_enrich_df = sort!(mut_enrich_df, "Enrichment_$sel_name", rev=true)  #sort by enrichment scores
       mut_enrich = rename_df(mut_enrich_df[:,["Seq","Enrichment_$sel_name"]],"trial_$i",1)
       mut_freq = rename_df(mut_enrich_df[:,["Seq","Freq_R1_$sel_name"]],"trial_$i",1)
       mut_rank = mut_enrich[:,["Seq"]]
       mut_rank[!,"Rank_$sel_name"*"_trial_$i"] = 1:size(mut_enrich,1)
       top_counts = threshold_extract(mut_enrich_df,"enriched", 0, sel_name)  # subset of significantly enriched mutations
       top_counts = rename_df(top_counts[:,["Seq","Freq_R1_$sel_name"]],"trial_$i",1)
       rest_muts = mut_enrich_df[.!in.(mut_enrich_df.Seq,Ref(top_counts[!,:Seq])),:]
       rest_muts = rename_df(rest_muts[:,["Seq","Freq_R1_$sel_name"]],"trial_$i",1)

       push!(enrichment_dfs, mut_enrich)
       push!(rank_dfs, mut_rank)
       push!(freq_dfs, mut_freq)
       push!(top_count_dfs,top_counts)
       push!(rest_count_dfs,rest_muts)

    end
    return (
        merge_dataframes(enrichment_dfs),
        merge_dataframes(rank_dfs),
        merge_dataframes(freq_dfs),
        merge_dataframes(top_count_dfs),
        merge_dataframes(rest_count_dfs)
    )
end


function mut_enrichment_depths(r0, r1, depths, trials, ref, sel_name)
    enrichment_sets, rank_sets, freq_sets, top_count_sets, rest_count_sets = [], [], [], [], []
    for depth in depths
        enrichment, rank, freq, top_count, rest_count = set_enrichment_stats(r0, r1, depth, trials, ref, sel_name)
        enrichment, rank, freq, top_count, rest_count = (
            rename_df(enrichment, "depth$depth", 1),
            rename_df(rank, "depth$depth", 1),
            rename_df(freq, "depth$depth", 1),
            rename_df(top_count, "depth$depth", 1),
            rename_df(rest_count, "depth$depth", 1)
        )
        push!(enrichment_sets, enrichment)
        push!(rank_sets, rank)
        push!(freq_sets, freq)
        push!(top_count_sets, top_count)
        push!(rest_count_sets, rest_count)
    end
    return (
        merge_dataframes(enrichment_sets),
        merge_dataframes(rank_sets),
        merge_dataframes(freq_sets),
        merge_dataframes(top_count_sets),
        merge_dataframes(rest_count_sets)
    )
end

# Calculating probabilities of extracting significantly enriched mutants at different sequencing depths
function find_probs(df, trials, depths)
    num_cols = size(df, 2)
    probs = DataFrame(ID = df.Seq)  
    for i in 2:10:num_cols
        col_range = i:min(i+9, num_cols)
        missing_counts = sum.(eachrow(ismissing.(df[:, col_range])))
        not_missing_prob = 1.0 .- missing_counts ./ 10
        depth_index = ((i - 2) ÷ 10) + 1
        col_name = Symbol("$(depths[depth_index])")
        probs[!, col_name] = not_missing_prob
    end
    return probs
end


# %% [code]
# Sequencing depth analyisis across all mutants and selections
sequencing_depths = [0.1,0.2,0.5,0.8,1,2,5,10,20,50,60] # coverage 
lib7_seqdepth_mutants = mut_enrichment_depths(aa_lib0, aa_lib7, sequencing_depths, 10, F1aa_ref, "Sel7")

enrich_seqdepth_mutants = lib7_seqdepth_mutants[1]
rank_seqdepth_mutants = lib7_seqdepth_mutants[2]
freq_seqdepth_mutants = lib7_seqdepth_mutants[3]
sig_enrich_seqdepth_mutants = lib7_seqdepth_mutants[4]
rest_seqdepth_mutants = lib7_seqdepth_mutants[5]

#CSV.write("D1_lib7_seqdepth_sig_enrich.csv", sig_enrich_seqdepth_mutants)
#CSV.write("D1_lib7_seqdepth_rest.csv", rest_seqdepth_mutants)


prob_sig_enrich = find_probs(sig_enrich_seqdepth_mutants, 10, sequencing_depths)
#CSV.write("D1_prob_sig_enrich.csv", prob_sig_enrich)

# %% [code]
# Adding enrichment and frequency scores to sig enriched variants identified

sig_enrich_seqdepth_mutants_stats=leftjoin(sig_enrich_seqdepth_mutants[:,[:Seq]],enrichment_7[:,[:Seq,:Freq_R1_sel7,:Enrichment_sel7,:EC_sel7]],on=:Seq)
sig_enrich_seqdepth_mutants_stats[sortperm(sig_enrich_seqdepth_mutants.Seq), :]
#CSV.write("D1_prob_sig_enrich_stats.csv", sig_enrich_seqdepth_mutants_stats)

# %% [code]
# Testing different depths for R0 with fixed rate of R1

function random_sets(data, percentage_depth, trials)
    sets = []
    for i in 1:trials
        N = Int(percentage_depth*400)
        indices = randperm(length(data))[1:N]
        subset = data[indices]
        push!(sets,subset)
    end
    return sets
end

function enrichment_sets(sets_r0, sets_r1, ref, mut, sel_name)
    enrichment_scores = []
    for i in 1:size(sets_r0)[1]
        enrich_set = mut_enrichment(sets_r0[i], sets_r1[i], ref, sel_name)
        enrichment_score = enrich_set[enrich_set[!, "Mutations_$sel_name"] .== mut, "Enrichment_$sel_name"]
        append!(enrichment_scores, enrichment_score)
    end
    avg_score = mean(enrichment_scores)
    std_score = std(enrichment_scores)
    return (avg_score,std_score)
end


function set_enrichment_stats_fix(r0, r1, depths_r0, depths_r1, trials, ref, sel_name)
    enrichment_dfs, rank_dfs, freq_dfs, top_count_dfs, rest_count_dfs = [], [], [], [], []
    trial_col = "Average_Enrichment_N$trials"
    r0_sets = random_sets(r0,depths_r0,trials)
    r1_sets = random_sets(r1,depths_r1,trials)
    for i in 1:trials 
       mut_enrich_df = mut_enrichment(r0_sets[i], r1_sets[i], F1aa_ref, sel_name)
       mut_enrich_df = sort!(mut_enrich_df, "Enrichment_$sel_name", rev=true)  #sort by enrichment scores
       mut_enrich = rename_df(mut_enrich_df[:,["Seq","Enrichment_$sel_name"]],"trial_$i",1)
       mut_freq = rename_df(mut_enrich_df[:,["Seq","Freq_R1_$sel_name"]],"trial_$i",1)
       mut_rank = mut_enrich[:,["Seq"]]
       mut_rank[!,"Rank_$sel_name"*"_trial_$i"] = 1:size(mut_enrich,1)
       top_counts = threshold_extract(mut_enrich_df,"enriched", 0, sel_name)  # subset of significantly enriched mutations
       top_counts = rename_df(top_counts[:,["Seq","Freq_R1_$sel_name"]],"trial_$i",1)
       rest_muts = mut_enrich_df[.!in.(mut_enrich_df.Seq,Ref(top_counts[!,:Seq])),:]
       rest_muts = rename_df(rest_muts[:,["Seq","Freq_R1_$sel_name"]],"trial_$i",1)

       push!(enrichment_dfs, mut_enrich)
       push!(rank_dfs, mut_rank)
       push!(freq_dfs, mut_freq)
       push!(top_count_dfs,top_counts)
       push!(rest_count_dfs,rest_muts)

    end
    return (
        merge_dataframes(enrichment_dfs),
        merge_dataframes(rank_dfs),
        merge_dataframes(freq_dfs),
        merge_dataframes(top_count_dfs),
        merge_dataframes(rest_count_dfs)
    )
end


function mut_enrichment_depths_fix(r0, r1, depths_r0, depths_r1, trials, ref, sel_name, fixed) #depths must be same length
    enrichment_sets, rank_sets, freq_sets, top_count_sets, rest_count_sets = [], [], [], [], []

    if fixed == "R0"
        for i in 1:length(depths_r1)
            depth = depths_r1[i]
            enrichment, rank, freq, top_count, rest_count = set_enrichment_stats_fix(r0, r1, depths_r0[i], depths_r1[i], trials, ref, sel_name)
            enrichment, rank, freq, top_count, rest_count = (
                rename_df(enrichment, "depth$depth", 1),
                rename_df(rank, "depth$depth", 1),
                rename_df(freq, "depth$depth", 1),
                rename_df(top_count, "depth$depth", 1),
                rename_df(rest_count, "depth$depth", 1)
            )
            push!(enrichment_sets, enrichment)
            push!(rank_sets, rank)
            push!(freq_sets, freq)
            push!(top_count_sets, top_count)
            push!(rest_count_sets, rest_count)
        end
    elseif fixed == "R1"
        for i in 1:length(depths_r0)
            depth = depths_r0[i]
            enrichment, rank, freq, top_count, rest_count = set_enrichment_stats_fix(r0, r1, depths_r0[i], depths_r1[i], trials, ref, sel_name)
            enrichment, rank, freq, top_count, rest_count = (
                rename_df(enrichment, "depth$depth", 1),
                rename_df(rank, "depth$depth", 1),
                rename_df(freq, "depth$depth", 1),
                rename_df(top_count, "depth$depth", 1),
                rename_df(rest_count, "depth$depth", 1)
            )
            push!(enrichment_sets, enrichment)
            push!(rank_sets, rank)
            push!(freq_sets, freq)
            push!(top_count_sets, top_count)
            push!(rest_count_sets, rest_count)
        end
    end

    return (
        merge_dataframes(enrichment_sets),
        merge_dataframes(rank_sets),
        merge_dataframes(freq_sets),
        merge_dataframes(top_count_sets),
        merge_dataframes(rest_count_sets)
    )
end

# %% [code]
#TESTING INFLUENCE OF R0 COVERAGE
sequencing_depths_r0 = [0.1,0.2,0.5,0.8,1,2,5,10,20,50,60] 
sequencing_depths_r1_60x = repeat([60], length(sequencing_depths_r0))
sequencing_depths_r1_10x = repeat([10], length(sequencing_depths_r0))
sequencing_depths_r1_5x = repeat([5], length(sequencing_depths_r0))

lib7_seqdepth_mutants_R1_60x = mut_enrichment_depths_fix(aa_lib0, aa_lib7, sequencing_depths_r0,sequencing_depths_r1_60x, 10, F1aa_ref, "Sel7","R1")
prob_sig_enrich_R1_60x = find_probs(lib7_seqdepth_mutants_R1_60x[4], 10, sequencing_depths_r0)

lib7_seqdepth_mutants_R1_10x = mut_enrichment_depths_fix(aa_lib0, aa_lib7, sequencing_depths_r0,sequencing_depths_r1_10x, 10, F1aa_ref, "Sel7","R1")
prob_sig_enrich_R1_10x = find_probs(lib7_seqdepth_mutants_R1_10x[4], 10, sequencing_depths_r0)

lib7_seqdepth_mutants_R1_5x = mut_enrichment_depths_fix(aa_lib0, aa_lib7, sequencing_depths_r0,sequencing_depths_r1_5x, 10, F1aa_ref, "Sel7","R1")
prob_sig_enrich_R1_5x = find_probs(lib7_seqdepth_mutants_R1_5x[4], 10, sequencing_depths_r0)

#CSV.write("D1_prob_sig_enrich_R1_60x.csv", prob_sig_enrich_R1_60x)
#CSV.write("D1_prob_sig_enrich_R1_10x.csv", prob_sig_enrich_R1_10x)
#CSV.write("D1_prob_sig_enrich_R1_5x.csv", prob_sig_enrich_R1_5x)

# %% [code]
#TESTING INFLUENCE OF R1 COVERAGE
sequencing_depths_r1 = [0.1,0.2,0.5,0.8,1,2,5,10,20,50,60] 
sequencing_depths_r0_60x = repeat([60], length(sequencing_depths_r1))
sequencing_depths_r0_10x = repeat([10], length(sequencing_depths_r1))
sequencing_depths_r0_5x = repeat([5], length(sequencing_depths_r1))

lib7_seqdepth_mutants_R0_60x = mut_enrichment_depths_fix(aa_lib0, aa_lib7, sequencing_depths_r0_60x,sequencing_depths_r1, 10, F1aa_ref, "Sel7","R0")
prob_sig_enrich_R0_60x = find_probs(lib7_seqdepth_mutants_R0_60x[4], 10, sequencing_depths_r1)

lib7_seqdepth_mutants_R0_10x = mut_enrichment_depths_fix(aa_lib0, aa_lib7, sequencing_depths_r0_10x,sequencing_depths_r1, 10, F1aa_ref, "Sel7","R0")
prob_sig_enrich_R0_10x = find_probs(lib7_seqdepth_mutants_R0_10x[4], 10, sequencing_depths_r1)

lib7_seqdepth_mutants_R0_5x = mut_enrichment_depths_fix(aa_lib0, aa_lib7, sequencing_depths_r0_5x,sequencing_depths_r1, 10, F1aa_ref, "Sel7","R0")
prob_sig_enrich_R0_5x = find_probs(lib7_seqdepth_mutants_R0_5x[4], 10, sequencing_depths_r1)

#CSV.write("D1_prob_sig_enrich_R0_60x.csv", prob_sig_enrich_R0_60x)
#CSV.write("D1_prob_sig_enrich_R0_10x.csv", prob_sig_enrich_R0_10x)
#CSV.write("D1_prob_sig_enrich_R0_5x.csv", prob_sig_enrich_R0_5x)

# %% [code]
# Calculating false negative and false positive rates 
true_positives =  prob_sig_enrich[1:19,:ID]
sequencing_depths_r1 = [0.1,0.2, 0.5, 1, 1.5, 2, 5,10,15,20,25,30,35] 
sequencing_depths_r0 = [0.1,0.2, 0.5, 1, 1.5, 2, 5,10,15,20,25,30,35] 

function coverage_accuracy_2D(r0,r1,depths_r0, depths_r1,trials,ref,sel)
    accuracies = []
    for i in 1:length(depths_r0)
        accuracy = DataFrame([ name =>[] for name in string.(sequencing_depths_r1)])
        depth = depths_r0[i]
        sequencing_depths_r0_rep = repeat([depth], length(depths_r0))
        seqdepth_mutants = mut_enrichment_depths_fix(r0, r1, sequencing_depths_r0_rep,depths_r1, trials, ref, sel,"R0")
        probs = find_probs(seqdepth_mutants[4], trials, depths_r1)
        total_sum = sum.(eachcol(probs[!,2:end])) # true positives + false positives
        true_pos_avg = mean.(eachcol(filter(row -> row.ID in true_positives, probs)[!,2:end]))
        false_pos_avg = mean.(eachcol(filter(row -> row.ID ∉ true_positives, probs)[!,2:end]))

        false_pos_prop = (sum.(eachcol(filter(row -> row.ID ∉ true_positives, probs)[!,2:end])))./total_sum # false discovery rate
        true_pos_prop = (sum.(eachcol(filter(row -> row.ID in true_positives, probs)[!,2:end])))./total_sum # precision

        probs_rest = find_probs(seqdepth_mutants[5], trials, depths_r1)

        false_negatives_sum = sum.(eachcol(filter(row -> row.ID in true_positives, probs_rest)[!,2:end]))
        true_positives_sum =  sum.(eachcol(filter(row -> row.ID in true_positives, probs)[!,2:end]))

        recall = true_positives_sum./(true_positives_sum.+false_negatives_sum)

        push!(accuracy,true_pos_avg,false_pos_avg,true_pos_prop,false_pos_prop,recall,false_negatives_sum)
        insertcols!(accuracy, 1, :Measure=>["True_positives_R0_$depth","False_positives_R0_$depth","True_positives_prop_R0_$depth","False_positives_prop_R0_$depth","Recall_R0_$depth","False_Negatives_R0_$depth"])
        push!(accuracies, accuracy)
    end
    accuracies = vcat(accuracies...)
    t1 =accuracies[startswith.(accuracies.Measure, "False_positives_R0"), :]
    t2 =accuracies[startswith.(accuracies.Measure, "True_positives_R0"), :]
    t3= accuracies[startswith.(accuracies.Measure, "False_positives_prop_R0"), :]
    t4= accuracies[startswith.(accuracies.Measure, "True_positives_prop_R0"), :]
    t5= accuracies[startswith.(accuracies.Measure, "Recall_R0"), :]
    t6= accuracies[startswith.(accuracies.Measure, "False_Negatives_R0"), :]
    return mapcols(col -> replace!(col, NaN=>0), vcat(t1,t2,t3,t4,t5,t6))
end


# %% [code]
lib7_seqdepth_mutants_R0R1 = coverage_accuracy_2D(aa_lib0, aa_lib7, sequencing_depths_r0,sequencing_depths_r1, 10, F1aa_ref, "Sel7")
CSV.write("D1_coverage_accuracy.csv", lib7_seqdepth_mutants_R0R1)

# %% [code]
#Flattening datasets

function flatten_coverage_accuracy(coverage_df,rows_select)
    subset_rows = coverage_df[startswith.(coverage_df.Measure, rows_select), :]
    flattened_data = stack(subset_rows, names(subset_rows)[2:end])
    rename!(flattened_data, :Measure => Symbol("R0_coverage_$rows_select"), :variable => :R1_Coverage, :value => Symbol(rows_select))
    return flattened_data
end


False_positives_flat = flatten_coverage_accuracy(lib7_seqdepth_mutants_R0R1, "False_positives_R0")
True_positives_flat = flatten_coverage_accuracy(lib7_seqdepth_mutants_R0R1, "True_positives_R0")
False_positives_prop_flat = flatten_coverage_accuracy(lib7_seqdepth_mutants_R0R1, "False_positives_prop_R0")
True_positives_prop_flat = flatten_coverage_accuracy(lib7_seqdepth_mutants_R0R1, "True_positives_prop_R0")
False_negatives_flat = flatten_coverage_accuracy(lib7_seqdepth_mutants_R0R1, "False_Negatives_R0")
R0_coverage = DataFrame(R0_Coverage = string.(getindex.(split.(False_positives_flat.R0_coverage_False_positives_R0,"R0_"),Ref(2))))

lib7_seqdepth_mutants_R0R1_flat = hcat(R0_coverage,False_positives_flat[!, Not(1)],True_positives_flat[!, Not(1:2)],False_positives_prop_flat[!, Not(1:2)],True_positives_prop_flat[!, Not(1:2)],False_negatives_flat[!, Not(1:2)])
CSV.write("D1_coverage_accuracy_flat.csv", lib7_seqdepth_mutants_R0R1_flat)


# %% [markdown]
#### Testing coverage on D4

# %% [code]
# D4 TEST

sequencing_depths_r0 = [0.5] 
sequencing_depths_r1_02x = repeat([0.2], length(sequencing_depths_r0))

lib7_seqdepth_mutants_R1_02x = mut_enrichment_depths_fix(aa_lib0, aa_lib7, sequencing_depths_r0,sequencing_depths_r1_02x, 10, F1aa_ref, "Sel7","R1")
prob_sig_enrich_R1_02x = find_probs(lib7_seqdepth_mutants_R1_02x[4], 10, sequencing_depths_r0)
#CSV.write("D1_prob_sig_enrich_R1_02x.csv", prob_sig_enrich_R1_02x)

# %% [markdown]
#### D4 Enrichment Analysis

# %% [code]
# Reference sequence based on the expected trimmed wildtype sequence - will be used for identification of mutations
cd("C://Users//PH//OneDrive - KU Leuven//PhD//Polymerase_Landscape_Mapping//05_Writing//01_Manuscripts//Selection_landscape_mapping//SI//individual_files//NGS_Analysis//")

F1aa_ref = aa"YLDFRSLYP"
codon_number_start = 402

lib_ID = dna"gatGCTACTCGATGGAAGATG"
lib_all = dna"CTaGGAGACCTCCTAGAGG"

mut_1 = dna"GAGAACATAGTG"
mut_2 = dna"TCAATCATCATC"

#Loading sequences matching IDs 
lib0 = load_fasta_file_trim("R0_5pt_D4_MiSeq.fasta", lib_ID, lib_all, mut_1, mut_2)
lib4 = load_fasta_file_trim("Sel4_5pt_D4_MiSeq.fasta", lib_ID, lib_all, mut_1, mut_2)
lib8 = load_fasta_file_trim("Sel8_5pt_D4_MiSeq.fasta", lib_ID, lib_all, mut_1, mut_2)
lib20 = load_fasta_file_trim("Sel20_5pt_D4_MiSeq.fasta", lib_ID, lib_all, mut_1, mut_2)

aa_lib0 = dna_2_aa(lib0)
aa_lib4 = dna_2_aa(lib4)
aa_lib8 = dna_2_aa(lib8)
aa_lib20 = dna_2_aa(lib20)

# %% [code]
# number of sequences post filtering and translating
println(" aa_lib0=",length(aa_lib0)," aa_lib4=",length(aa_lib4)," aa_lib8=",length(aa_lib8)," aa_lib20=",length(aa_lib20))
# number of unique sequences
println(" aa_lib0=",length(keys(countmap(aa_lib0)))," aa_lib4=",length(keys(countmap(aa_lib4)))," aa_lib8=",length(keys(countmap(aa_lib8)))," aa_lib20=",length(keys(countmap(aa_lib20))))
# number of matching unique sequences
println(length(intersect(aa_lib0,aa_lib4,aa_lib8,aa_lib20)))

# %% [code]
#Enrichment
enrichment_4 = mut_enrichment(aa_lib0, aa_lib4, F1aa_ref, "sel4")
enrichment_8 = mut_enrichment(aa_lib0, aa_lib8, F1aa_ref, "sel8")
enrichment_20 = mut_enrichment(aa_lib0, aa_lib20, F1aa_ref, "sel20")

sel_enrichment = outerjoin(enrichment_4, enrichment_8, enrichment_20,on = :Seq)
sel_enrichment_match = innerjoin(enrichment_4, enrichment_8,enrichment_20, on = :Seq)
show(sel_enrichment, allcols = true)
show(sel_enrichment_match, allcols = true)
CSV.write("D4_enrichment_df.csv", sel_enrichment)
CSV.write("D4_enrichment_match_df.csv", sel_enrichment_match)

# %% [code]
# Sig enriched sequences + rest (sig depleted/neutral mutants)
enriched4 = threshold_extract(enrichment_4,"enriched", 0, "sel4")
enriched8 = threshold_extract(enrichment_8,"enriched", 0, "sel8")
enriched20 = threshold_extract(enrichment_20,"enriched", 0, "sel20")
CSV.write("D4_sig_sel4.csv", enriched4)
CSV.write("D4_sig_sel8.csv", enriched8)
CSV.write("D4_sig_sel20.csv", enriched20)
sel4_rest = filter(row -> !(row.Seq in enriched4.Seq), enrichment_4)
sel8_rest = filter(row -> !(row.Seq in enriched8.Seq), enrichment_8)
sel20_rest = filter(row -> !(row.Seq in enriched20.Seq), enrichment_20)
CSV.write("D4_rest_sel4.csv", sel4_rest)
CSV.write("D4_rest_sel8.csv", sel8_rest)
CSV.write("D4_rest_sel20.csv", sel20_rest)


# %% [code]
# Matching mutants with significant changes in frequency
significant4 = threshold_extract(enrichment_4,"pval", 0.05, "sel4")
significant8 = threshold_extract(enrichment_8,"pval", 0.05, "sel8")
significant20 = threshold_extract(enrichment_20,"pval", 0.05, "sel20")

sig_muts_df = innerjoin(significant4[!,[1,2,7,8,9,10]],significant8[!,[1,2,7,8,9,10]],significant20[!,[1,2,7,8,9,10]], on= :Seq)
show(sig_muts_df, allcols=true)
CSV.write("D4_sig_muts_matched.csv", sig_muts_df)

# %% [code]
ref_seq = dna"GCTACTCGATGGAAGATGCGAAGGTCACATACGAGCTTGGGAAGGAGTTCCTTCCGATGGAGGCCCAGCTTTCTCGCTTAATCGGCCAGTCCCTCTGGGACGTCTCCCGCTCCAGCACTGGCAACCTCGTTGAGTGGTTCCTCCTCAGGAAGGCCTATGAGAGGAATGAGCTGGCCCCGAACAAGCCCGATGAAAAGGAGCTGGCCAGAAGACGGCAGAGCTATGAAGGAGGCTATGTAAAAGAGCCCGAGAGAGGGTTGTGGGAGAACATAGTGTCAATCATCATCACCCACAACGTCTCGCCGGATACGCTCAACAGAGAAGGATGCAAGGAATATGACGTTGCCCCACAGGTCGGCCACCGCTTCTGCAAGGACTTCCCAGGATTTATCCCGAGCCTGCTaGGAGACCTCCTAGAGG"
lib0_fidelity = pol_fidelity_stats(lib0, ref_seq, "lib0")
lib4_fidelity = pol_fidelity_stats(lib4, ref_seq, "lib4")
lib8_fidelity = pol_fidelity_stats(lib8, ref_seq, "lib8")
lib20_fidelity = pol_fidelity_stats(lib20, ref_seq, "lib20")
lib_fidelity = vcat(lib0_fidelity,lib4_fidelity,lib8_fidelity,lib20_fidelity)
CSV.write("D4_overall_lib_fidelity.csv", lib_fidelity)

# %% [code]
function seq_subset_list(DNA_seqs::Vector{Tuple{LongSequence{DNAAlphabet{4}}, LongSequence{DNAAlphabet{4}}}},list::Vector{LongAA})
    seqs = Tuple{LongSequence{DNAAlphabet{4}}, LongSequence{DNAAlphabet{4}}}[]
    for i in 1:length(DNA_seqs)
        rna_seq = convert(LongRNA{2}, DNA_seqs[i][2])
        if length(rna_seq) % 3 == 0 
            mut_trans = BioSequences.translate(rna_seq, allow_ambiguous_codons=false, alternative_start=false)
            if mut_trans in list 
                push!(seqs, (LongDNA{4}(DNA_seqs[i][1]),(LongDNA{4}(DNA_seqs[i][2]))))
            end
        end
    end
    return seqs
end

# %% [code]
# Subsetting reads to top mutants with >1000 counts across selections to calculate mutant-specific fidelity
count4 = threshold_extract(enrichment_4,"count", 1000, "sel4")
count8 = threshold_extract(enrichment_8,"count", 1000, "sel8")
count20 = threshold_extract(enrichment_20,"count", 1000, "sel20")
sig_muts_list = outerjoin(count4,count8,count20, on=:Seq)[!,:Seq]

lib0_mut_fidelity = mut_fidelity(seq_subset_list(lib0,sig_muts_list), ref_seq, "lib0")
lib4_mut_fidelity = mut_fidelity(seq_subset_list(lib4,sig_muts_list),ref_seq, "lib4")
lib8_mut_fidelity = mut_fidelity(seq_subset_list(lib8,sig_muts_list),ref_seq, "lib")
lib20_mut_fidelity = mut_fidelity(seq_subset_list(lib20,sig_muts_list),ref_seq, "lib20")

# %% [code]
lib0_mut_fidelity_rn = rename_df(lib0_mut_fidelity, "R0", 1)
lib4_mut_fidelity_rn = rename_df(lib4_mut_fidelity, "Sel4", 1)
lib8_mut_fidelity_rn = rename_df(lib8_mut_fidelity, "Sel8", 1)
lib20_mut_fidelity_rn = rename_df(lib20_mut_fidelity, "Sel20", 1)
muts_fidelity_sels = outerjoin(lib0_mut_fidelity_rn,lib4_mut_fidelity_rn,lib8_mut_fidelity_rn,lib20_mut_fidelity_rn, on= :ID)
#adding enrichment values to fidelity scores
#sel_enrichment.ID = string.(sel_enrichment.Seq)
enrichment_fidelity = innerjoin(muts_fidelity_sels,sel_enrichment, on = :ID)
CSV.write("D4_Muts_Fidelity_Top.csv", enrichment_fidelity)


# %% [code]
#adding enrichment values to fidelity scores
enrichment_fidelity_cor = innerjoin(sel_enrichment, muts_fidelity_sels, on = :ID)
enrichment_fidelity_cor = select(enrichment_fidelity_cor, r"^(Mut|ID|Enr|Freq|bases|ins|del|subs).*$")
# substracting R0 values from fidelity scores
enrichment_fidelity_cor[!, [:insertions_R0,:insertions_Sel4,:insertions_Sel8,:insertions_Sel20]] .-= enrichment_fidelity_cor[!, :insertions_R0]
enrichment_fidelity_cor[!, [:deletions_R0,:insertions_Sel4,:deletions_Sel8,:deletions_Sel20]] .-= enrichment_fidelity_cor[!, :deletions_R0]
enrichment_fidelity_cor[!, [:substitutions_R0,:insertions_Sel4,:substitutions_Sel8,:substitutions_Sel20]] .-= enrichment_fidelity_cor[!, :substitutions_R0]
CSV.write("D4_enrichment_fidelity_cor.csv", enrichment_fidelity_cor)
```
